# *In vitro* plasticity between ureteric epithelial and distal nephron identity and maturity is controlled by extracellular signals

**DOI:** 10.1101/2025.06.03.657609

**Authors:** Pamela Kairath, Pei Xuan Er, Sean Wilson, Irene Ghobrial, Jessica M. Vanslambrouck, Chen Yi-Hsien, Shondra M. Pruett-Miller, Sanjay Jain, Melissa H. Little

## Abstract

Several studies have described human pluripotent stem cell (hPSC)-derived ureteric epithelium, which in the embryo arises from the nephric duct and forms the collecting ducts of the kidney. However, hPSC-derived distal nephron epithelium can also adopt a ureteric phenotype, despite this not occurring during embryogenesis. In this study, RET^tdTomato^ and GATA3^mCherry^ reporter lines were used to further investigate this plasticity. Induction of anterior intermediate mesoderm resulted in the spontaneous formation of an epithelial plexus with a nephric duct-like identity. Subsequent addition of RSPO1 induced patterning of distalized nephrons, including distal convoluted tubule and thick ascending limb of loop of Henle but lacking proximal segments or glomeruli. This epithelium showed a capacity to adopt ureteric epithelial or nephric duct-like states in *ex vivo* co-culture in response to external cues. The same epithelium seeded as single cells in Matrigel formed epithelial spheroids and adopted a RET+ ureteric tip identity. This *in vitro* continuum between nephric duct, ureteric epithelium and distal nephron illustrates the role of the microenvironment in cellular identity.

**Highlights and eTOC:** *Highlights:*

- Distal renal epithelium shows substantial plasticity *in vitro*
- Identity can be shifted towards distal nephron, including distal convoluted tubule and loop of Henle, in response to prolonged RSPO1
- Distal epithelial identity can shift in *ex vivo* culture in response to surrounding signals
- Collecting duct is supported by Matrigel culture

*eTOC blurb:* Kairath and colleagues investigate the plasticity of induced renal epithelium in response to external signalling cues *in vitro* and *ex vivo*. Initial patterning generates a plastic epithelium able to pattern to specific distal nephron. The same epithelium, cultured in Matrigel patterns to ureteric epithelium.

## Introduction

A key component of the kidney is the system of collecting ducts to which each nephron is connected and through which the urinary filtrate passes to exit the kidney. This branched epithelial network is unique to the metanephros, the third pair of excretory organs to form during mammalian development. Unlike the nephrons of the metanephros, which arise from a nephrogenic mesenchyme^1^, the collecting ducts form via the dichotomous branching of an ingrowing ureteric epithelium (UE)^2^. This UE is itself formed as a side branch of the nephric duct (ND), also referred to as the Wolffian duct or the mesonephric duct^3^. The ND arises from the anterior intermediate mesoderm (IM) and then extends caudally along with the forming body axis. In some vertebrates, including fish and frog, a metanephric kidney does not form with the pronephros and mesonephros providing larval and adult excretory function respectively. Despite this, the patterning and segmentation of pronephric, mesonephric and metanephric nephrons is highly conserved^4, 5, 6^. Around 10dpc in the mouse, the ureteric epithelium (UE) arises as an outpouching of epithelial cells from the distal ND called the ureteric bud (UB). This side-branch grows towards the metanephric mesenchyme in response to tropic factors from this domain. Direct interaction between the tips of the UE (ureteric buds) and the surrounding nephron progenitor mesenchyme results in dichotomous branching to form a ureteric epithelial tree comprised of proliferating RET-expressing ureteric tip cells and less proliferative ureteric stalks^2^. The ureteric stalks pattern to form distinct cortical, outer medullary and inner medullary collecting ducts through which the urinary filtrate passes to exit the kidney. The distal portions of the nephrons themselves, including the distal convoluted tubule (DCT) and distal straight tubule (thick ascending limb of the Loop of Henle; TAL) connect to the ureteric tips via the connecting segment (CS). While in the mouse, lineage tracing shows a metanephric mesenchymal origin for the CS^7^, there is considerable congruence in gene expression between ureteric epithelium and distal nephron elements including the CS, DCT and TAL^8, 9^. In human, there is clear evidence for the presence of both water reabsorbing AQP2+ Principal cells and acid/base regulating intercalated cells within regions of the collecting duct tree as well as the DCT and CS and clear evidence that the relative ratio of these component cell types can shift with injury or drug use^10, 11, 12^.

Our ability to dissect kidney development in a human setting has significantly improved in recent years with the development of protocols for the stepwise directed differentiation of human pluripotent stem cells to adopt a renal fate (reviewed in^13^). The resulting kidney organoids contain patterning and segmenting nephrons, comprised of glomeruli, proximal and distal nephron as evidenced by gene expression, protein production and morphology. However, initial kidney organoid protocols did not show evidence of RET^+^ ureteric tips or a coordinated branching ureteric epithelium. Our own recent reanalysis also showed poor representation of ureteric epithelium and limited maturation of distal nephron elements^14, 15^. Based on the embryonic origin of the UE from the ND, several groups have now reported the distinct generation of induced UE from pluripotent stem cells using protocols that mimic a more anterior intermediate mesoderm^16, 17, 18, 19, 20^. All such studies have cultured these populations in Matrigel, in which they variably survive, proliferate and bud. Indeed, such induced-UE (iUE) cultures have been used to model autosomal dominant polycystic kidney disease ^21^. More recently, we have shown that such a induced UE identity can be achieved *in vitro* via the selective propagation of GATA3^+^ distal nephron epithelium in the presence of UE-supportive growth factors, suggesting a epithelial plasticity *in vitro* apparently not present *in vivo* ^14^. This plasticity evident between distal nephron and UE *in vitro* is complicated by the lack of transcriptional data on ND in both human or mouse. This represents a ‘blindspot’ for our ability to classify early ND within such pluripotent stem cell-derived structures.

In this study, we investigate the plasticity of induced renal epithelium generated from multiple human induced pluripotent stem cell (iPSC) reporter lines, including both RET^tdTomato^ (Supplementary Figure 1) and GATA3^mCherry 22^. By applying existing knowledge of early embryonic patterning, we initially generate an induced nephric duct-like epithelium (iND) within a surrounding mesenchyme both via 2D monolayer and 3D organoid cultures. This resulted in the spontaneous formation of LHX1^+^ PAX2^+^ epithelial tubules with evidence of both tdTomato and Cherry reporter expression. In 3D culture, this iND self-organised to form extending and branching RET^+^ epithelial tips within a surrounding mesenchyme. The addition of RSPO1 to such organoids induced the formation of regions of SLC12A3^+^GATA3^+^ distal convoluted tubule (DCT) and SLC12A1^+^GATA3^−^ thick ascending limb (TAL), suggesting induced distal nephron (iDN) patterning and maturation. The same epithelium showed considerable plasticity *in vitro*, adopting induced UE (iUE) and iND fates in co-culture assays. Indeed, individual epithelial cells isolated from such organoids displayed clonal formation of RET^+^ epithelial cysts in Matrigel. These observations indicate a continuum of cellular identity between iND, iUE and iDN and an ability for cell fate to shift in response to the extracellular matrix environment and growth factor signalling.

## Results

### Patterning to a GATA3^+^ RET^+^ epithelium

Cells that migrate out of the presomitic mesoderm early contribute to the nascent mesoderm of the anterior body axis, including the ND which arises from the anterior IM. The ND arises via the collaborative action of mediolateral gradients of BMP/Activin in combination with high retinoic acid signalling. In mouse, correct patterning, localization, and orientation of the ND requires co-expression of a core gene network composed of *Lim1(Lhx1)/Pax2/Pax8* ^3,23^. The Pax2^+^ Lhx1^+^ intermediate mesoderm forms a Gata3^+^ ND which in turn induces the expression of the receptor tyrosine kinase, *RET* ^24, 25, 26^. While there have been extensive studies into the transcriptional profile and morphogenesis of the dichotomously branching ureteric tree, much less is understood about the ND. However, early markers of ND include *Gata3* and *Ret* in mouse ^27^ with human orthologs of these genes also expressed in the UE of mouse and human. *Gata3* is also expressed in the early DN of both mouse^8^ and human^28^ with expression ultimately restricted to the distal straight and distal convoluted tubules in mouse^9^ and GATA3 mutations in humans resulting in Barakat syndrome (HDR syndrome)^29^. However, the DN does not express *RET in vivo*.

In this study, we used both *GATA3^mCherry+^* ^22^ and RET^tdTomato^ (https://www.rebuildingakidney.org/chaise/record/#2/Cell_Line:Reporter_Cell_Line/RID=Q-2CW0) (Suppl. Figure 1A) human iPSC reporter lines to generate induced ND, UE and DN (iND, iUE, iDN) epithelial states *in vitro*. We have previously reported an association between the duration of Wnt signalling and the initiation of an anterior (short duration) versus posterior (long duration) intermediate mesoderm^30^. As the ND arises from the anterior IM, monolayer cultures of reporter lines were exposed to a short (2 days) duration and low (4 to 6 µM) concentration of the glycogen synthase kinase GSK-3 inhibitor, CHIR99021, compared to previous kidney organoid protocols^13^ (Figure 1A). From day 3 to day 7, APEL2 media was supplemented with growth factors reported to promote early migration from the posterior primitive streak and/or support iND^18, 31^. This included FGF9 (200 ng/ml), heparin (HA) (1 µg/ml), GDNF (100 ng/ml), All-*trans* retinoic acid (0.2µM) (tRA), 9-*cis*-retinoic acid (0.2µM) (9-*cis*RA), and the BMP inhibitor LDN (0.1µM). Within the monolayer cultures, areas of high cellular density arose from day 5 with robust mCherry or tdTomato fluorescence by day 7, with 24.31% and 83.63% of the culture FACS-sorted as RET^tdTomato+^ or GATA3^mCherry+^ respectively (Figure 1BC). qPCR of monolayer cultures versus FACS-sorted GATA3^mCherry+^ cells confirmed enrichment for the ND markers *LHX1, RET* and *PAX2* as well as the downstream RET target genes, *GFRA1* and *ETV5*, in the sorted populations (Figure 1D). At day 7, these regions did not represent epithelial tubules, hence cultures were either extended for a further 7 days in monolayer format (Figure 1 EF) or dissociated and reaggregated to form micromasses which were cultured on Transwell filters for a further 7 days (Figure 1 GH). Antibody staining at day 14 revealed the spontaneous formation of a distinct epithelial network in both monolayer (Figure 1F, Suppl. Figure 1B) and organoid (Figure 1H, Suppl. Fig. 1C) cultures. Native tdTomato fluorescence from reporter lines suggested the induction of RET (Suppl. Figure 1BC) and GATA3 in a PAX2^+^ ECAD^+^ epithelium which was surrounded by a non-epithelial stroma (Figure 1FH). In monolayer cultures, there was evidence of small PAX2^+^ mesenchymal aggregates in monolayer cultures which appeared to be fusing to the epithelium as would a neo-nephron in the fish (Figure 1F).

**Figure 1.**
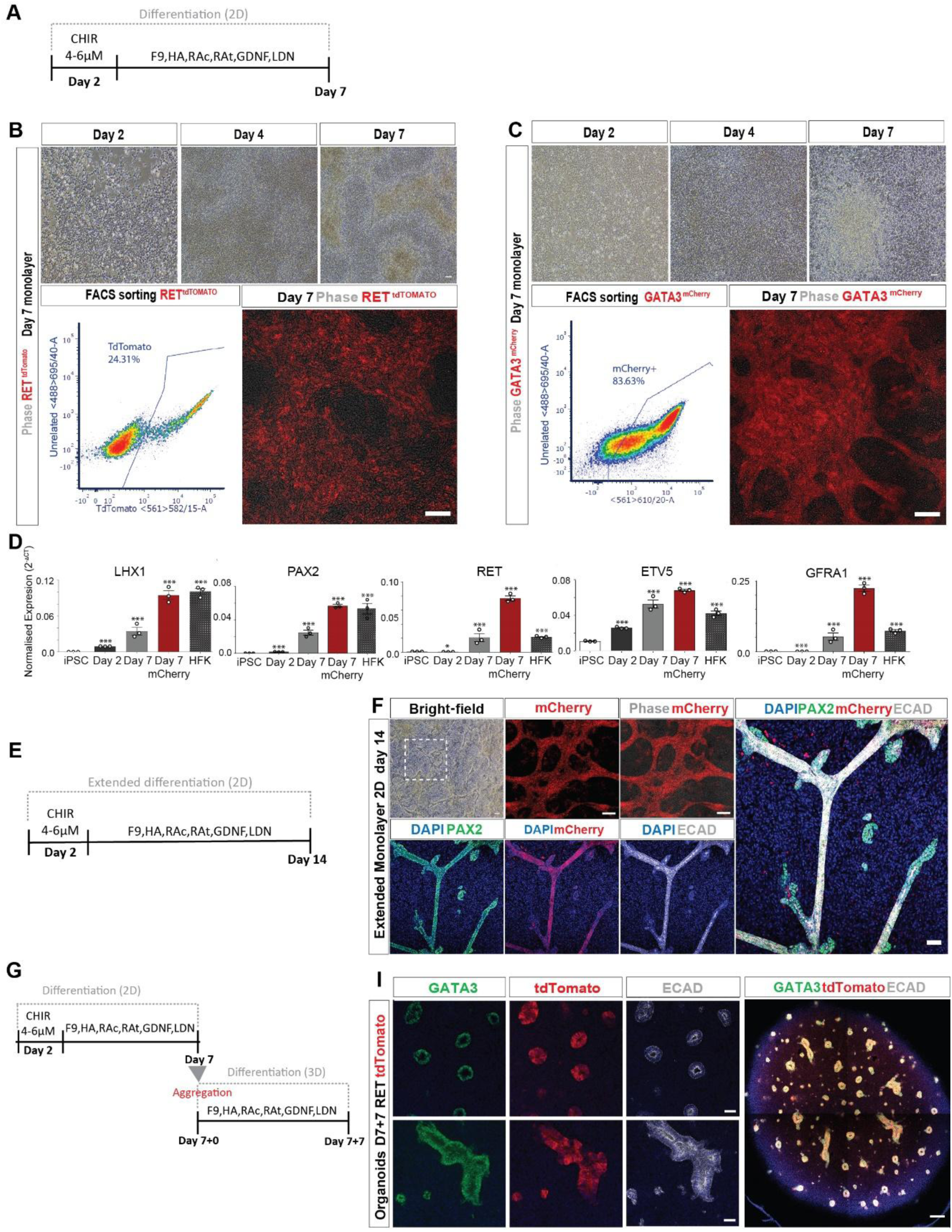
Directed differentiation towards anterior intermediate mesoderm results in the spontaneous formation of a nephric duct-like epithelial plexus in either monolayer or organoid culture formats. **A.** Diagram of 7 day monolayer culture for differentiation of human PSCs to AIM. CHIR, CHIR99021 (GSK3b antagonist), F9, FGF9; HA, heparin; RAc, Cis-retinonic acid; Rat, Trans-retinoic acid; GDNF, glial derived neurotrophic factor; LDN, LDN193189 (ALK2/3 inhibitor). **BC.** Differentiation timecourse using RET^tdTomato^ (**B**) and GATA3^mCherry^ (**C**) reporter lines. Top: Brightfield images at day 0, 2 and 7. Bottom left. FACS sorting revealed the presence of tdTomato (24.31%, 3wells of one independent experiment pooled (**B**)) and mCherry (83.63%, 3 wells pooled from two independent experiments (**C**)) cell populations at day 7. See supplemental Figure 1D and 1E for negative controls. Bottom right: native fluorescence of monolayer cultures at Day 7. Scale bars: 100uM **D.** qPCR time-course for expression of key nephric duct / ureteric epithelial markers in RNA extracted from whole monolayer cultures (day 2 and day7) and FACs-purified GATA3^mCherry^ (day 7) compared to undifferentiated iPSC and human fetal kidney (HFK). Error bars represent SEM and significance was determined using two tailed unpaired *t-*test (*P≤0.05; **P≤0.01; ***P≤0.001), n=3 independent wells representing parallel differentiation experiments. **E.** Diagram of 14day monolayer culture for differentiation of human PSCs (extended differentiation). **F.** Live fluorescence imaging (top row) and immunofluorescence staining of extended 2D culture of GATA3^mCherry^ reporter line imaged at day 14. Scale bar: 100 and 50 uM respectively **G.** Diagram of extended differentiation with an initial 7 days of monolayer culture and a further 7 days after organoid formation (aggregation). **H.** Immunofluorescence staining of organoid cultures of RET^tdTomato^ reporter line imaged at day 7+7. Scale bar: 50 and 200uM

### Single cell profiling identifies an induced nephric duct-like epithelial population

The induction of a spontaneously arising epithelium showing reporter expression from both the RET and GATA3 loci did not distinguish identity between induced ND and UE identities. Previous protocols for the generation of iUE have proposed a transition through ND from which UE progenitors have been FACS sorted via specific cell surface markers^18, 19^. To investigate the identity of the epithelium within these organoid cultures, single cell transcriptional profiling was performed on day 7 + 7 organoids generated from the RET^tdTomato^ reporter line (Supplementary Figure 1) using the 10X platform. 5,098 cells passing quality control were grouped into nine distinct clusters (Figure 2A, Supplementary Table 1). The dominant cluster (cluster 0) represented 29.7% of the cells within the organoids. Cells in this cluster were epithelial (*CDH1*^+^*EPCAM*^+^) and displayed differential expression (DE) of nephric duct genes including *PAX2/8*, *GATA3* and *LHX1* (Figure 2B,C). They also showed DE expression of many genes associated with ureteric epithelial identity, including *HNF1B, EMX2, COL18A1, WFDC2* and *KRT8* (Figure 2C). The expression of downstream targets of the GDNF/RET pathway, including *GFRA1, ETV4/5* and *SPRY1*, while also expressed in this cluster, were more broadly expressed in non-epithelial clusters consistent with a much wider cellular response to the presence of GNDF within the media (Figure 2B,C). Notably, the epithelial cluster also showed marked DE gene expression of genes previously identified by Taguchi *et al.*^18^ as UE progenitor genes, including *CXCR4* and *KIT* (Figure 2B,C, Suppl. Figure 1B). While these markers have previously been used in combination to isolate double positive progenitors, reanalysis of this cluster alone resolved five subpopulations (Figure 2D, Supplementary Table 2) showing differential co-expression of *GATA3, KIT* and *LGR4* versus *CXCR4* and *LGR5* (Figure 2E). While there was overlap in *KIT* and *CXCR4* expression, *KIT* more strongly associated with *GATA3* expression. While transcriptional profiling of human ND is not available, we also understand little about the associated stroma that forms alongside the ND. However, Clusters 2, 3 and 5 revealed DE for *WT1* and *HOXA10*, both of which mark the nephrogenic mesenchyme.

**Figure 2.**
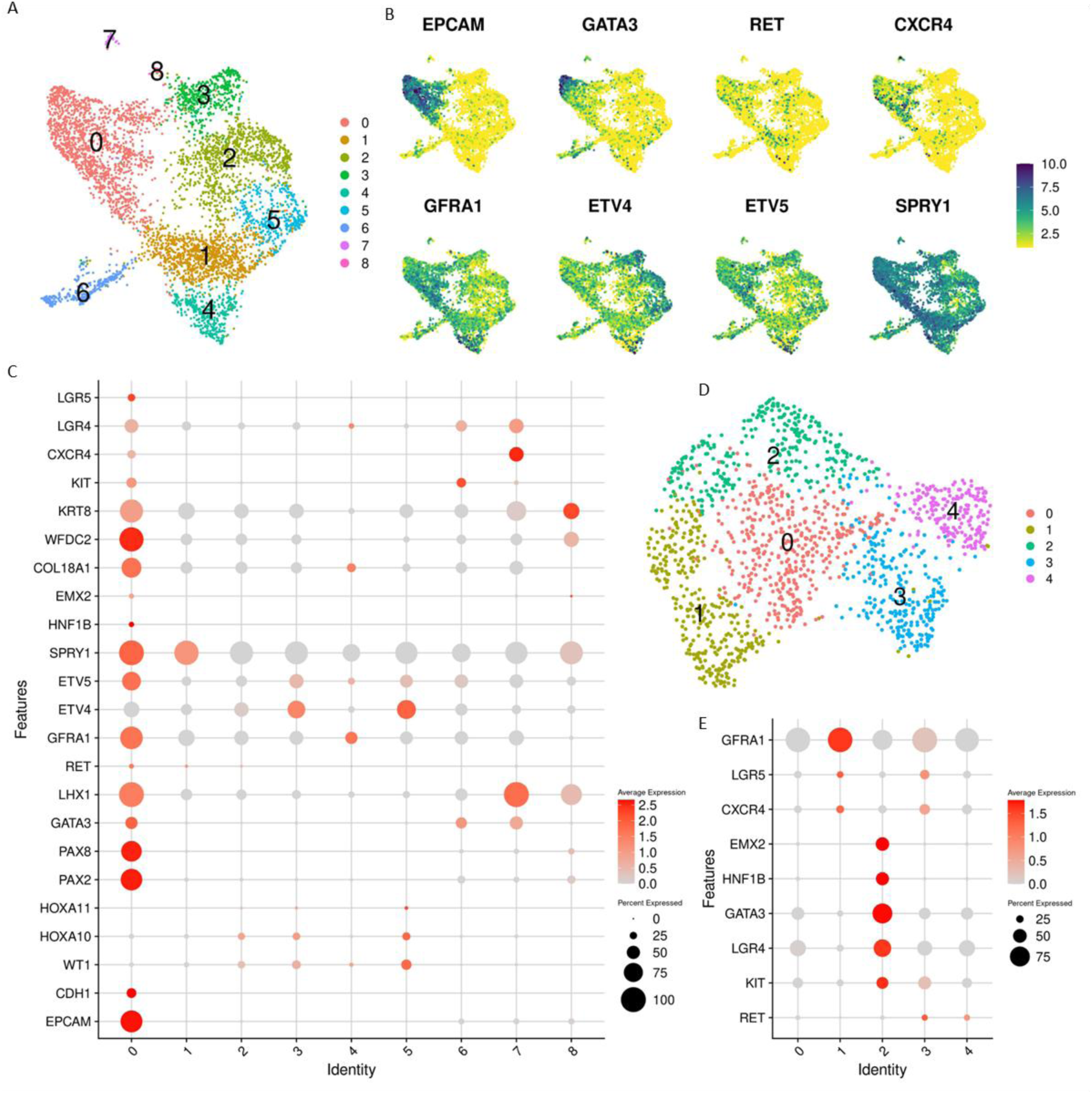
Analysis of cellular identity at a single cell level suggests an indeterminate induced nephric duct-like phenotype. **A.** Single cell RNA-sequencing of pooled day 14 organoids combined from a single differentiation experiment. Results are plotted in the first two UMAP dimensions, showing 9 clusters. **B.** Distribution of key gene expression plotted on the first two UMAP dimensions. **C.** DotPlot showing the expression of key markers of distal nephron and ureteric epithelium within the clusters shown in **D.** Cluster 0 re-analysed and plotted in the first two UMAP dimensions, showing 5 clusters. **E.** DotPlot showing the expression of key markers within the clusters.

Reporter expression confirmed the presence of *RET* promoter activity within these organoids. However, *RET* expression was only detected in a few cells and not localised to a specific cluster (Figure 2E). Surprisingly, *RET* expression was highest in subclusters 3 and 4, whereas the highest *GATA3* expression was observed in subcluster 3 along with *KIT* and *LGR4* expression (Figure 2E). Conversely, subcluster 2 showed highest expression of *GFRA1*, *LGR5* and *CXCR4*, suggesting distinct cellular states as marked by *KIT* and *CXCR4*. Without the overt presence of branching or evidence of specific mature ureteric cell types, we describe this epithelium as nephric duct (ND)-like or induced ND (iND).

### RSPO1 mediates the effect of WNT signalling supporting distalized epithelial patterning

There is considerable evidence for the supporting role of canonical WNT signalling in the maintenance of stem cell progenitors involved in kidney development. While in mouse, Wnt9b and Wnt4 have been classically regarded as crucial ligands for initiating nephron formation^32, 33^, more recent studies have shed some light on the importance of canonical WNT signalling on the UE tip^34, 35^ and/or the distal nephron niche^36,37^. Amplification of Wnt signals via R-spondin signalling is also involved in kidney development. RSPO signalling via Lgr4 has been reported to be required for ureteric branching morphogenesis with a loss of Lgr4 resulting in reduced GATA3 expression and premature differentiation^38^. Yuri et al^31^ described the expression of both *Rspo1* and *Rspo3* in the metanephric mesenchyme, while *Lgr4* was expressed here and in the underlying UE tips. Here, the synergistic action of RSPO and FGF signalling favoured proliferation and branching of E11.5 murine ureteric bud (UB), upregulating the ureteric tip markers *Ret*, *Wnt11* and *Etv5*. In mouse, *Lgr5* is a distinct marker of the distal nephron giving rise to the thick ascending limb of Henle’s loop and the distal convoluted tubule as a result of a clonal expansion^36^.

Based on differential expression of *LGR4* and *LGR5* in distinct subclusters of our iND (Fig 2), we investigated whether the subsequent addition of RSPO1 (300ng/ml) induced a UE identity or conversely enhanced a distal nephron epithelial fate (Figure 3, Suppl. Figure 3). In response to RSPO1, GATA3^mCherry^ epithelial cells began to coalesce to form budding epithelial elements which rapidly unified as a connected epithelial plexus across the entire organoid (Figure 3B, left panel) with 33.68% of the organoid FACS sorted as GATA3^mCherry+^ (Figure 3B, right panel). Extended regions of GATA3^+^ epithelium was observed but no evidence of proximal nephron segments or glomeruli (Figure 3BCD; Suppl. Figure 3A). Within the core of the organoids, the epithelium adopted an extensive epithelial field morphology from which tubular ‘fingers’ arose with distinct KRT8^+^ECAD^+^ staining, a clear lumen and apical aPKC protein (Figure 3E). Organoids displayed segmented epithelia with clear co-expression of UMOD and mCherry. However, they also contained, distinct regions of SLC12A3^+^mCherry^+^ epithelium, suggestive of distal convoluted tubule (DCT) and SLC12A1^+^mCherry^−^ epithelium suggestive of TAL patterning (Figure 3E). The same patterning was evident when generated using a parental hPSC line (Suppl. Figure 3C) with clear evidence of ECAD^+^ distal epithelium beyond the region of GATA3^+^ epithelium. Hence, epithelial segmentation within organoids showed the formation of defined distal nephron segments. This is consistent with previous reports showing that the synchronous activation of Wnt signalling and BMP inhibition induces a complete distalization of the nephron epithelium within embryonic mouse kidney explants^37^. We define these as ‘distalized’ kidney organoids with the induced DN epithelium capable of segmenting into distinct distal epithelial segments. While evidence of loop of Henle segments has previously been identified within kidney organoids^30, 39, 40^, this is the first report of patterning to SLC12A3+ DCT. This iDN shows the initial iND represents a plastic epithelium capable of adopting cellular identities not generated from the ND *in vivo*.

**Figure 3.**
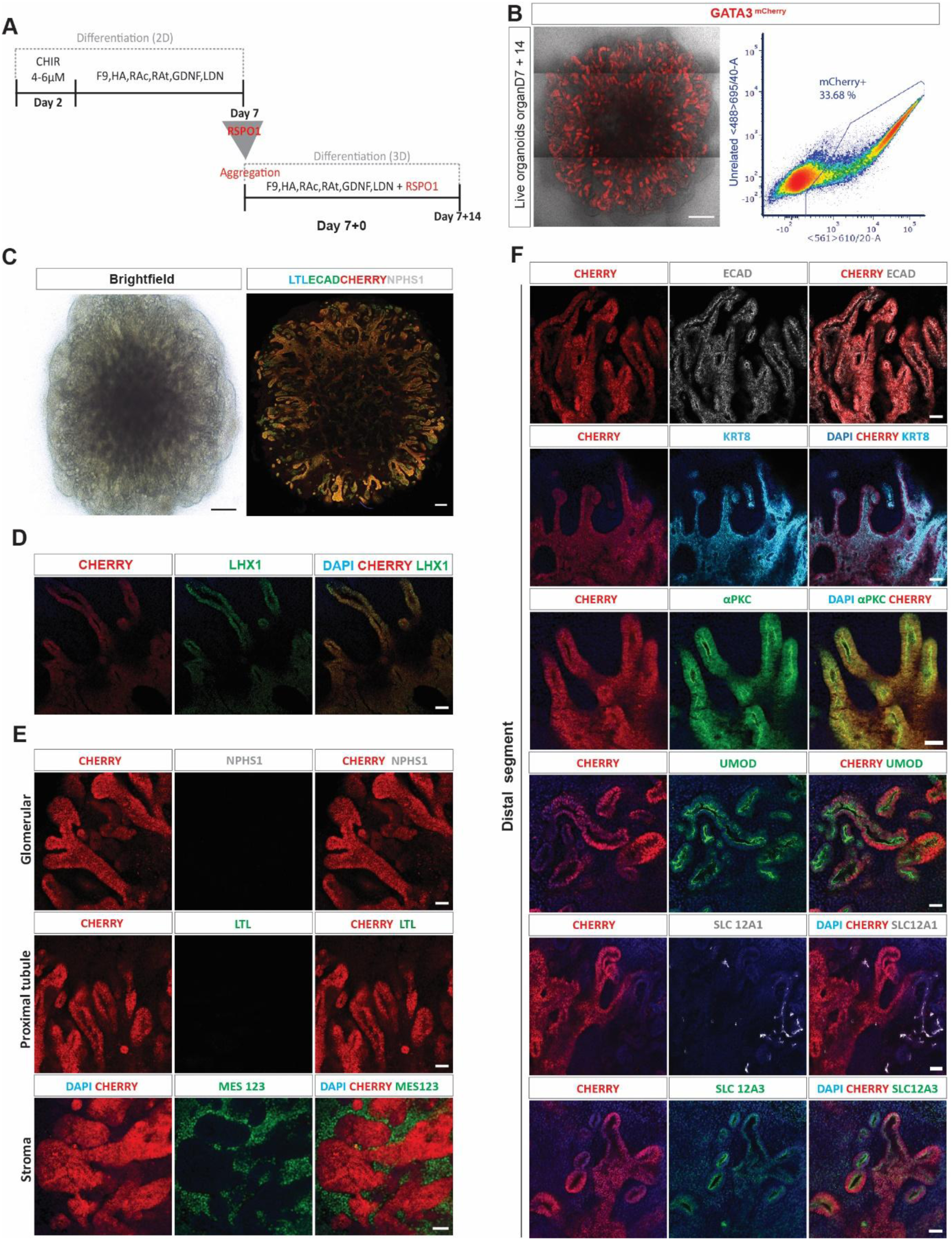
Generation of renal distal tubule, including distal convoluted tubule and thick ascending limb of the Loop of Henle, in response to RSPO1. **A.** Diagram of differentiation approach using micromass organoid culture from day 7 together with the addition of RSPO1 at organoid aggregation. **B.** Detection of endogenous mCherry in live organoids (Scale bar: 500uM) and FACS plot showing the presence of mCherry-positive cell population (33.68%) isolated at day 7+14 (data from pooled organoids from a single differentiation experiment). **C.** Brightfield (Scale bar: 500uM) and immunofluorescence (Scale bars: 200 uM) images of an entire organoid at day 7+14 generated using the GATA3^mCherry^ reporter line. Immunostaining was performed for proximal tubule (LTL, blue), distal epithelium (ECAD, green), mCherry and podocytes (NPHS, grey). See also Supplementary Figure 3. **D.** High magnification images of mCherry^+^ epithelium within these distalized organoids showing staining for the nephric duct marker, LHX1. **E.** Evidence within 3D distalized kidney organoids for the presence of stroma (MEIS123), but no evidence for proximal tubule segments (LTL) or glomeruli (NPHS1). **F.** Immunostaining of distalized kidney organoids for markers of distal nephron components and their relative expression of mCherry. Staining with CDH1, KRT8 and aPKC illustrates the presence of clear tubular lumens coincident with mCherry. Evidence is also seen for the formation of distal convoluted tubule (SLC12A3) and loop of Henle (SLAC12A1) segments which are not mCherry^+^. Scale bars: 50uM. All immunofluorescence images (**B-F**) are representative examples of multiple repeated differentiation experiments.

### Transcriptional evidence of distal nephron fate in response to extended RSPO1

As previously noted, the overlap in gene expression between the ND, UE and particular regions of the distal nephron makes definitive identification of these distinct epithelial subtypes difficult *in vitro*. There is distinct congruence of gene expression observed between UE and connecting segment, including the expression of *CALB1*. However, there are distinct expression patterns identified within other regions of the distal epithelium, including the expression of *SLC12A3* in the DCT. To further investigate the distalization of the epithelium within the RSPO1-patterned organoids, we again performed single cell transcriptional profiling. Organoids were profiled at day 21, one sample having RSPO added at day 7 while another at day 14. 6012 and 5715 cells passed quality control respectively (Figure 4). These samples were integrated using Seurat prior to dimensional reduction, which showed little difference between the component cell types in each dataset (Suppl. Figure 4). The graph-based clustering identified four main clusters (Figure 4A, Supplementary Table 3), the largest having an epithelial identity with expression of *GATA3*, *LHX1*, *PAX2* and *MAL*, and hence having high congruence with both ureteric epithelium / distal tubule (Figure 4B). Re-clustering of these epithelial cells identified 4 sub-clusters (Figure 4C, Supplementary Table 4), one of which was cell cycling related. Clusters 0 and 2 had considerable *GATA3* expression while co-expressing distal nephron markers such as *IRX1*, *FXYD2*, *TMEM52B* and *KCNJ1* as well as the Loop of Henle marker *SLC12A1* (Figure 4D, Suppl. Figure 4). Using *DevKidCC*, an unbiased method for cell identification within developing kidney tissue (Wilson et al., 2021), we compared the cellular identity profile of cells within the initial Day 7+7 organoids (Figure 2), extended RSPO1-treated distalized organoids and previously reported human pluripotent stem cell-derived UE defined by Mae *et al.*^16^ (Figure 4EF). While the Mae dataset had a ∼27% contribution of ureteric epithelium with almost no nephron, both day 7+7 and extended RSPO1-treated organoids showed the presence of nephron epithelia but no ureteric epithelia. Application of the next tier of cellular classification showed the nephron population within distalized organoids was predominantly early distal nephron, while the ureteric epithelium within Mae et al^16^ was predominantly ureteric tip cells. This may suggest that our initial iND was simply an early distal nephron progenitor.

**Figure 4.**
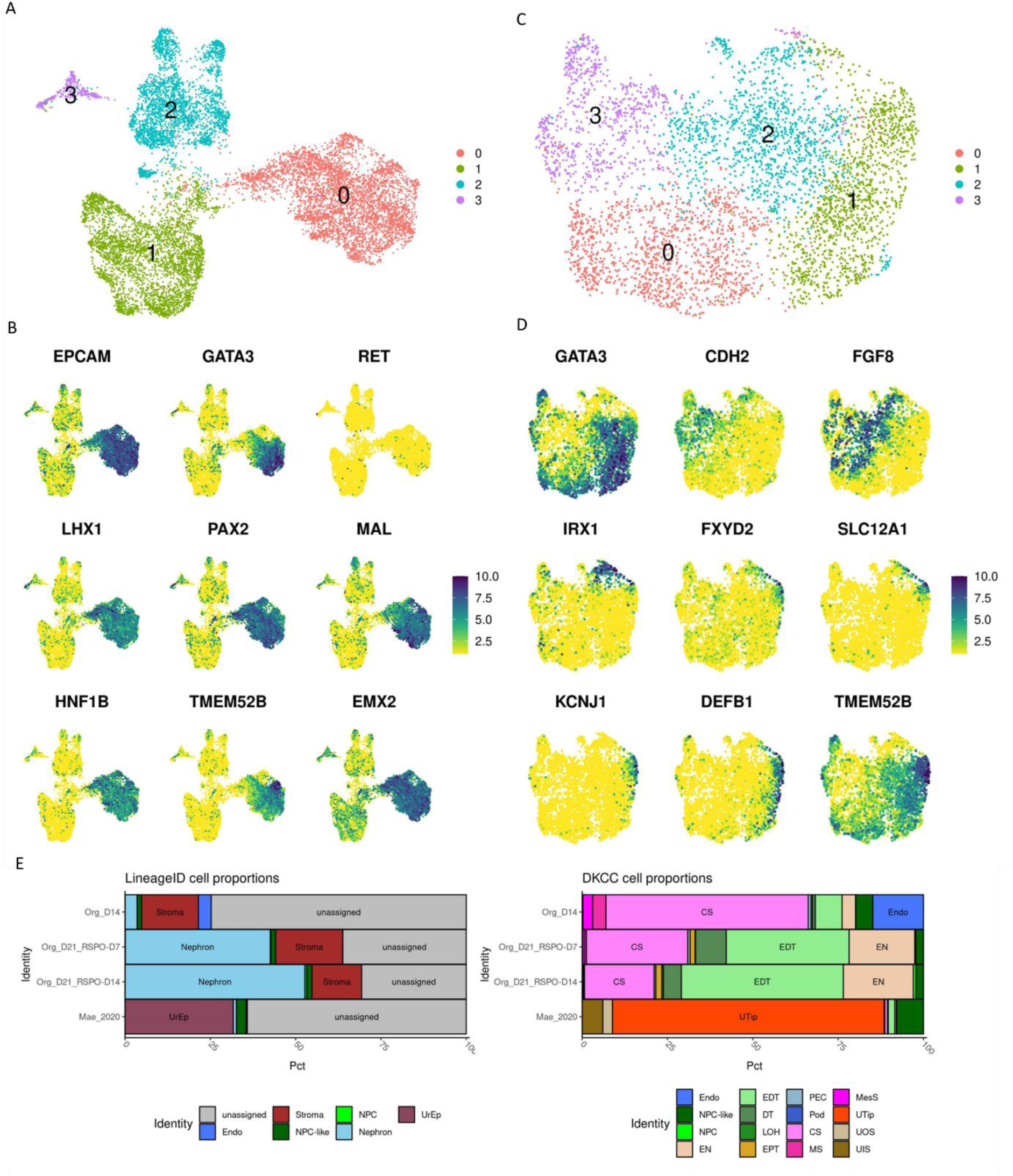
Single cell transcriptional profiling of distalized kidney organoids after short (7 days) or long (14 days) addition of RSPO1 identifies conversion to distal nephron. **A.** Single cell RNA-sequencing of two organoids samples of day 21 organoids with RSPO added at day 7 or day 14, integrated and plotted in the first two UMAP dimensions grouped as 4 main clusters. **B.** Expression distribution of key gene markers of distal nephron and ureteric epithelium. **C.** Re-analysis of the epithelial cluster (cluster 0) plotted in the first two UMAP dimensions, grouped as 4 main clusters. **D.** Expression of key markers show populations of early distal nephron (GATA3, CDH2, FGF8), Loop of Henle (IRX1, FXYD2, SLC12A1) and other distal nephron (KCNJ1, DEFB1, TMEM52B) markers. **E.** Kidney cell classification tool DevKidCC identified the lineage identities of cells in our organoid datasets in comparison to that of Mae et al^16^, highlighting a difference in the epithelium present being distal nephron in our datasets while being ureteric epithelial-like in Mae.

While we have focussed on the identity of the epithelium within these organoids, the surrounding stroma is likely to have played both an instructive role in epithelial identity as well as responding to the growth factor environment provided. We sought to identify the nature of this surrounding stroma based upon transcriptional profile. Dimensional reduction and cell clustering identified 3 other broad cell classifications in this data, a *TBX2/3/5* stromal cluster, and two *SOX2* positive mesenchymal clusters. *SOX2* is known to mark the neuromesodermal precursors in the tailbud region able to adopt both neural and mesodermal tissues^41^. The *TBX2/3/5* cluster showed distinct subclusters of *LHX9*^+^ and *GATA6*^+^ cells which may represent gonadal and adrenal progenitors respectively. Notably absent was any evidence of nephron progenitor or endothelial populations, as have been described in kidney organoids. Despite the presence of GDNF in the media, there was no evidence of a *SOX10*^+^ neural crest population. The Hox code of all clusters did not extend beyond *HOX9*, suggesting an anterior identity, and there was no evidence of the *TBX18* mesenchyme reported to arise along the ureteric bud as it grows into the metanephros, although a small region of *FOXD1* positive cells was present within a *TCF15*^+^*PAX3*^+^ cluster. A recent study of mouse ND differentiation identified the expression of *TCFAP2A/B* as marking a subtype of nephric duct progenitors^42^. These genes were expressed in the *GATA3*^+^ epithelial cluster, but also marked a small distinct cluster (cluster 3) also expressing *PAX3, SLIT2, ROBO2, EMX2, SOX2* and *EPCAM*.

In summary, the majority of the epithelium present under these conditions has adopted a distal nephron identity, including features of CS, DCT and TAL. No evidence of glomeruli or proximal nephron is present. In addition, the surrounding stroma suggests a more anterior region that the metanephros.

### Distalized organoid epithelium adopts a ureteric epithelial identity within mouse embryonic kidney

Distalized organoids contained a complex epithelium with distal nephron subsegments. While such elements do not give rise to collecting duct *in vivo*, we have previously shown that iUE can be generated from nephron epithelium^14^. To assess the identity and plasticity of the renal epithelium within distalized organoids, these were recombined with dissociated E13.5 embryonic mouse kidney^43, 44, 45, 46^. The incorporation of undifferentiated GATA3^mCherry^ hPSC resulted in random mCherry^+^ cells within the explant stroma (Suppl. Figure 5A). However, when mCherry^+^ cells FACS-sorted from distalized organoids were combined with dissociated Hoxb7GFP embryonic kidneys at 5 or 10% of total cells (Figure 5A), these adopted a clear ureteric epithelial fate (Figure 5B, Suppl. Figure 5B-D). At 5% of total cells, the mCherry^+^ cells could be seen interdigitating with GFP^+^ UE while when present at 10% of total cells, extended segments of branched mCherry^+^ epithelium was evident, as anticipated for UE. As such, this environment supported the adoption of a UE identity; iUE.

**Figure 5.**
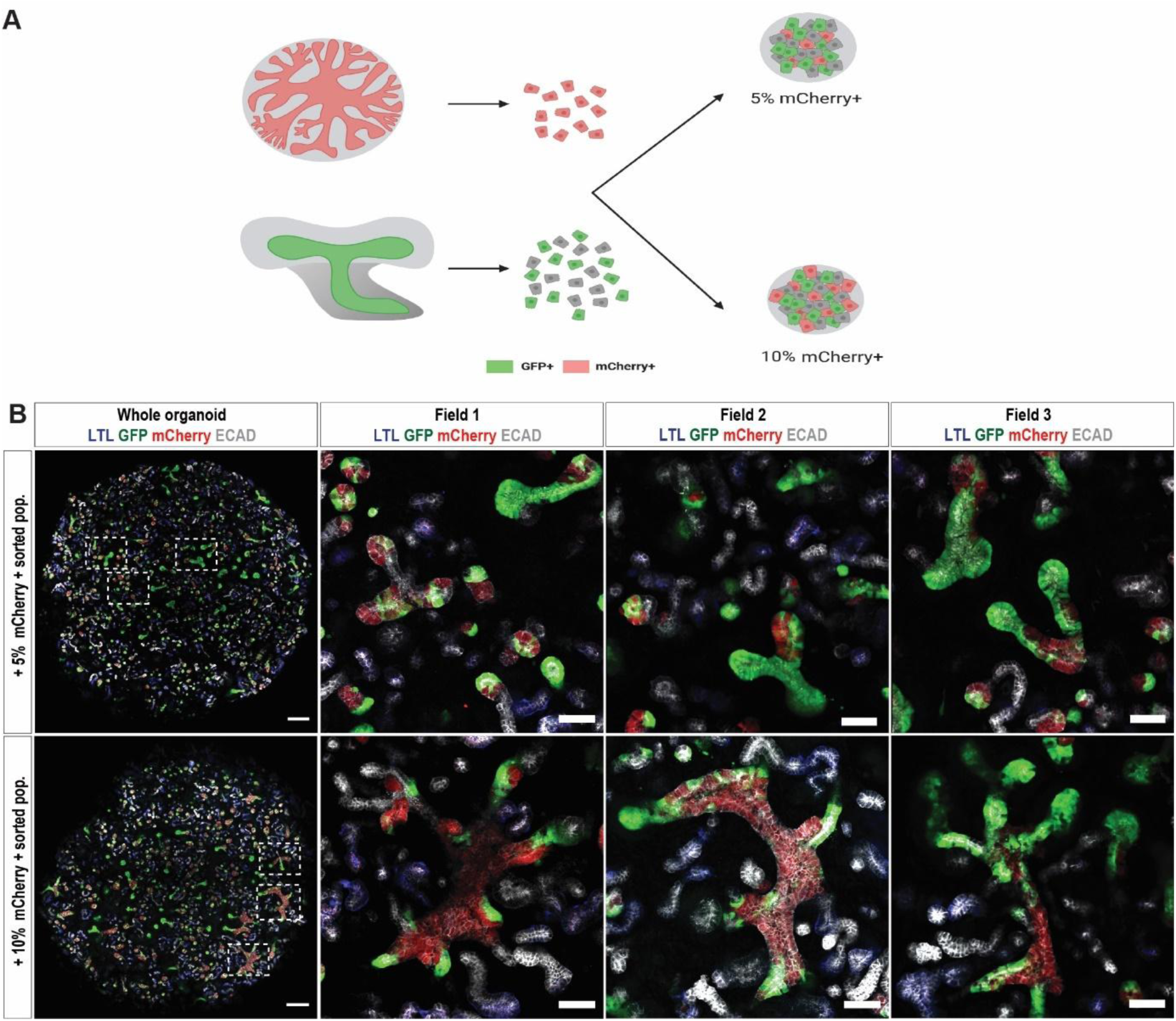
Selective integration of FACS-isolated mCherry^+^ epithelium into embryonic mouse ureteric epithelium. **A.** Schematic diagram depicting the recombination assay for assessing renal potential of mCherry^+^ expressing cells. **B.** Confocal images of recombinations between dissociated 12.5dpc Hoxb7GFP mouse embryonic kidney and 5% (top panels) or 10% (bottom panels) of dissociated distalized kidney organoids. At least 3 organoids evaluated for each condition with presented images extracted from one example of each % of reaggregated cells. Images were acquired 6 days after reaggregation. Immunostaining was performed to identify proximal tubule (LTL, blue), GFP (green), mCherry (red) and ECAD (grey). Right panels show higher magnification examples of mCherry integration from regions boxed at left. Scale bars: 200 and 50uM

### Epithelial plasticity in response to surrounding embryonic stroma

A recent study, using iUE differentiated from mouse embryonic stem cells, demonstrated that the identity of this iUE shifted in response to the surrounding tissue, presumably via stromal or ECM signals^47^. This was achieved by grafting iUE into the cortex of embryonic mouse kidney explants, resulting induction of nephron formation around this iUE. Conversely, iUE adopted a nephric duct-like phenotype when placed into the peri-Wolffian mesenchyme^47^.

Using this co-culture approach, we investigated the plasticity of our distalized organoid epithelium in response to nephrogenic versus peri-Wolffian stroma. Mouse 11.5-12.5 Hoxb7-EGFP embryonic kidneys, dissected to include the adjacent nephric duct and surrounding peri-Wolffian stroma, were chosen to facilitate visualisation of the EGFP^+^ murine ureteric tree and nephric duct. Manually dissected mCherry^+^ epithelium from distalized organoids was transplanted at different locations within respect to the embryonic tissue (Figure 6). Co-cultures were grown on DMEM/F12 for a further 3-4 days and imaged daily. When placed adjacent to the nephrogenic mesenchyme, Six2-expressing cells actively migrated towards the human mCherry^+^ epithelia, suggesting a ureteric tip identity (Figure 6AB). mCherry^+^ distal organoid epithelia transplanted into the peri-Wolffian mesenchyme showed evidence of dilation, the induction of the urothelial protein UPK3A and the formation of surrounding α-smooth muscle actin (Figure 6CD), similar to the reported response of iUB to this environment^47^. Finally, when grafted adjacent to an injured HOXB7-GFP tree, mCherry^+^ distalized epithelial grafts connected and formed a continuum with Hoxb7GFP mouse UE (Figure 6F-H).

**Figure 6.**
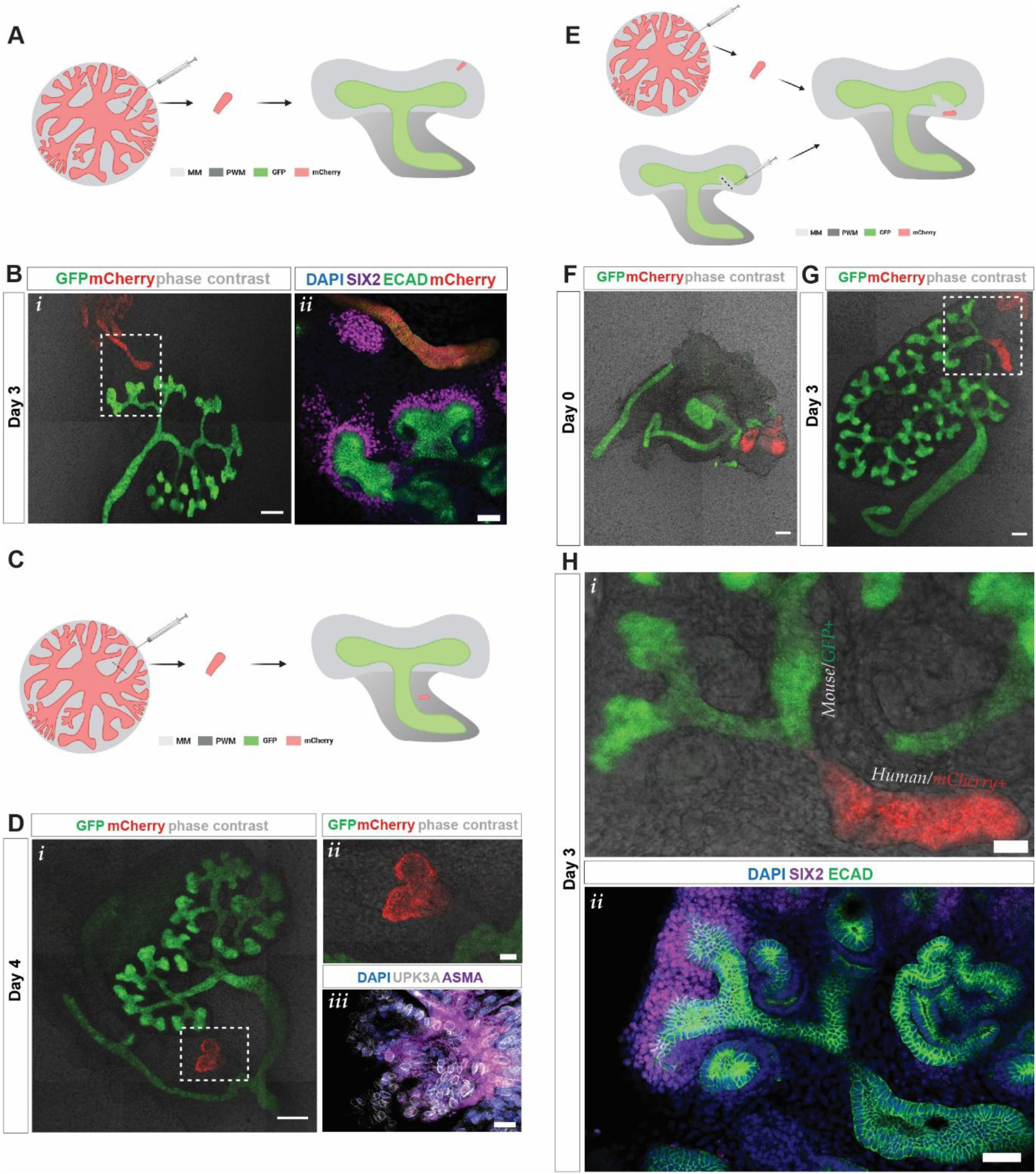
Dissected mCherry^+^ epithelium from distalized organoids can adopt the identity of the surrounding microenvironment. **A**. Diagram illustrating placement of dissected DN within nephrogenic field. All approaches were performed at least 2 times **B.** Example 3 day co-culture of dissected mCherry^+^ epithelium and explanted 11.5dpc Hoxb7GFP mouse kidney with placement adjacent to the MM. Left panel (i): low magnification showing relative positioning of mCherry+ epithelium with kidney explant. Scale bar: 200 uM. Right panel (ii): Immunostaining illustrating the presence of Six2^+^ mouse cap mesenchyme adjacent to the mCherry^+^ epithelium. Scale bars: 50 uM. **C**. Diagram illustrating placement of dissected DN within the peri-Wolffian mesenchyme. **D.** Example 4-day co-culture of dissected mCherry+ epithelium and explanted Hoxb7GFP mouse kidney with placement within the PWM. i) low magnification showing relative positioning of mCherry+ epithelium with kidney explant. Scale bar: 200 uM; ii) high magnification live imaging of boxed region from i); iii) Immunostaining of the mCherry+ epithelium with Uroplakin 3a (UPK3A), alpha smooth muscle actin (ASMA) and DAPI. Scale bars: 20uM **E**. Diagram illustrating placement of dissected DN within nephrogenic field in which ureteric tips have been damaged. **F.** Live imaging at Day 0 of co-culture showing the proximity of the mCherry+ epithelium and the damaged GFP+ ureteric tree. Scale bar: 100uM **G.** Live imaging at Day 3 of coculture showing connectivity between mCherry+ epithelium and GFP+ ureteric epithelium. Scale bar: 100uM **H**. High magnification images of the region boxed in G. Top panel (i) shows live imaging of mouse ureteric epithelium connecting human mCherry+ epithelium. Lower panel (ii) is subsequent immunostaining of the same material showing the relative location of the Six2^+^ cap mesenchyme around the mouse ureteric tips. Scale bars: 50 uM.

### Formation and maturation of ureteric epithelium from single epithelial progenitors supports plasticity of cellular identity

These reaggregation and co-culture studies suggested that distalized organoids contain epithelial cells able to adopt a ND/UE-like phenotype with the identity of these cells influenced by the surrounding environment. However, the profiling suggested that this epithelium does not commence with a UE identity. Previous studies describing the formation of UE from both mouse and human PSCs have isolated epithelial cells and cultured such progenitors in the presence of Matrigel free of the surrounding mesenchyme^16, 17, 18^. In mouse, Yuri et al^31^ previously showed that a single UE progenitor cell can clonally give rise to ureteric epithelial spheroids when cultured in Matrigel culture. No hPSC-derived iUE protocol has proven a capacity to generate epithelium from a single starting cell. We investigated the response of our induced distal epithelium to *in vitro* culture in Matrigel when plated at single cell seeding density. Individual FACS-sorted mCherry^+^ cells from day7+14 GATA3^mCherry^ reporter-derived organoids (Figure 7AB) were seeded at limiting dilution into 50% Matrigel and grown for 12 days using the following growth factor combination: Y-27632, FGF9 (200 ng/ml), and heparin (1 µg/ml), GDNF (100 ng/ml), All-trans retinoic acid (0.2µM) (tRA), 9-cis-retinoic acid (0.2µM) (9-cisRA), LDN (0.1µM), RSPO1(300 ng/ml). Individual mCherry^+^ cells were able to survive and develop colonies with spheroid-like structures which continued to expand across 12 days in culture (Figure 7C). qPCR revealed upregulation of the ureteric tip markers RET, GFRα1, ETV5, DUSP6 in response to the addition of Matrigel (Figure 7D). Immunostaining confirmed the presence of RET, ECAD, GATA3 and PAX2 and these induced UB-like structures exhibited epithelial lumen formation and apical-basal polarity, as revealed by αPKC staining (Figure7EF). The presence of tdTomato fluorescent epithelial clusters also arose from individually seeded cells isolated from distalized organoids generated using the RET^tdTomato^ reporter line (Figure 7G). Hence, distalized organoid epithelium contains progenitors able to generate induced UE spheroids in Matrigel.

**Figure 7.**
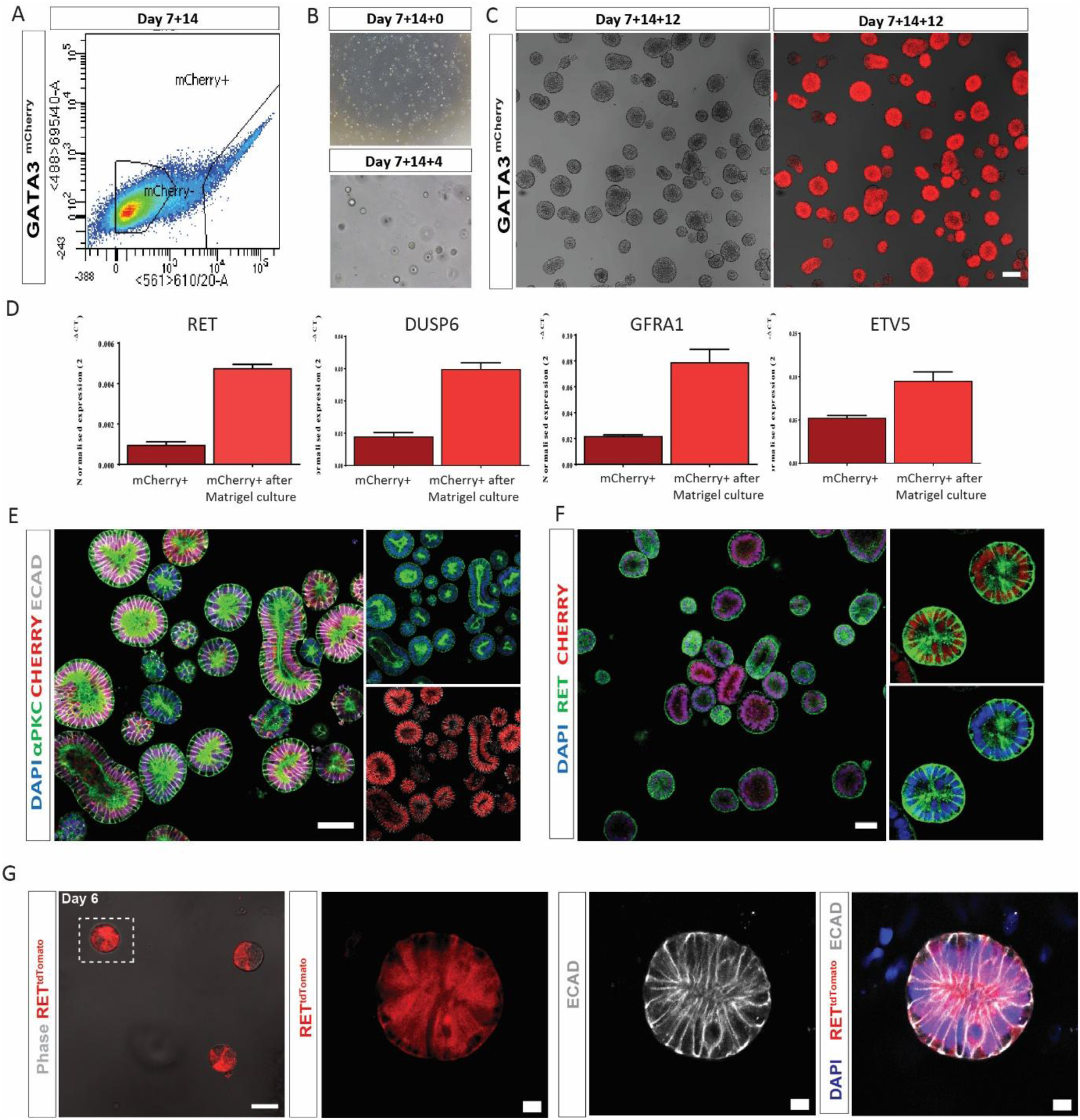
Maintenance of a ureteric epithelial phenotype is supported by culture in collagen/laminin rich extracellular matrix. **A.** FACS sorting from day7+14 distalized organoids **B.** Individual mCherry^+^ expressing single cells (Day 7+14+0; top)) display colony forming efficiency within Matrigel across four days (7+14+4; bottom). Single progenitor seeding was achieved via dilution as described previously^31^. **C.** Phase contrast and live fluorescence of mCherry^+^ 3D epithelial spheroids after 14 days in Matrigel. Scale bar = 100uM **D.** qPCR analysis of mCherry^+^ cells at isolation and after culture in Matrigel showing upregulation of ureteric tip markers *RET, DUSP6, GFRA1* and *ETV5*. Error bars represent SEM and significance was determined using two tailed unpaired *t-*test (*P≤0.05; **P≤0.01; ***P≤0.001), n=3 independent wells per condition from a single example differentiation experiment. **E.** Immunofluorescence of 3D epithelial spheroids in Matrigel showing staining for ECAD, GATA3 and αPKC proteins. Scale bar = 50uM **F.** Immunofluorescence of 3D epithelial spheroids in Matrigel showing staining for RET protein and mCherry. Scale bar = 50 uM. **G.** High magnification brightfield / live fluorescence imaging (top) and immunofluorescence (bottom) of individual epithelial spheroids at Day 6 of culture generated using the RET^tdTomato^ reporter line. Scale bar = 50 and 10uM

## Discussion

In this study, we have facilitated the spontaneous formation of an early distal renal epithelial plexus within a surrounding stroma via directed differentiation to anterior intermediate mesoderm using reporter iPSC lines for the nephric duct / ureteric epithelial genes RET and GATA3. This induced epithelium represents a plastic progenitor population able to spontaneously adopt distinct distal nephron patterning, including distal convoluted tubule and thick ascending limb of the loop of Henle, but no ability to pattern proximal nephron or glomerulus. The same epithelium, depending upon the stroma, matrix and/or growth factor environment, could also adopt a ureteric tip / collecting duct phenotype. This reaffirms our previous report of a capacity to transition between distal nephron and collecting duct in vitro^14^, providing new approaches for the generation of distinct renal epithelial cell types.

Several previous studies have described protocols reported to generate UE from pluripotent stem cells. The initial patterning approach used in this study is not dissimilar to previous approaches defined as generating an anterior mesoderm within which a nephric duct progenitor arises^16–20^. However, only one study other than this one demonstrated a capacity for individual progenitors to form a RET^+^ epithelial spheroid^19^. In that instance, UE progenitors were isolated by FACS based upon expression of *cKIT*. Culture in Matrigel resulted in the formation of induced ureteric buds (iUB) from which induced collecting duct (iCD) were generated. Our analysis of this early epithelial identity suggests an indeterminate expression profile similar to nephric duct, ureteric epithelium and distal nephron. Single cell analysis revealed distinct KIT or CXCR4 positive progenitors, in contrast to the double positive population previously observed^18^. A separation was also observed between the expression of LGR4 and LGR5, hence it is not clear if R-Spondin 1 selectively or uniformly expands these distinct populations. Of note, the inclusion of R-Spondin 1 is a feature of most adult epithelial stem cell expansion media.

The ability to definitively identify nephric duct has been challenging due to the lack of a transcriptional reference in either mouse or human. A recent comprehensive single cell and spatial transcriptomic analysis of the developing mouse urinary tract describes a series of nephric duct progenitor states proposed to represent distinct regions of the extending ND^42^. While all progenitors express *Gata3*, it was proposed that a Tcfap2b^+^ population (NdPr2) separated the most anterior ND progenitors (NdPr1) from the Ret^+^ tip cells (NdPr4). *Gata3* expression increased along the length of the ND with NdPr4 cells lost if Gata3 was deleted. While showing clear progenitor heterogeneity, *c-kit* expression was detected in NdPr2 and NdPr4 suggesting that FACS isolation using this marker was likely still isolating a mixed population. Our own analysis showed a separation between the expression of *KIT* and *CXCR4* with distinct *KIT^+^LGR4^+^GATA3^+^*and *CXCR4^+^LGR5^+^GFRA1^+^* cellular states evident in the ND-like organoids. A better understanding of the potential of each cell state may assist in directing to even more specific endpoints.

*In vivo*, the branching ureteric epithelium of the metanephric kidney does not form via transition from a distal nephron epithelium. The early epithelial state reported here is likely analogous to a nephric duct progenitor with this state capable of a variety of cellular endpoints. In the presence of the surrounding stroma within a distalized organoid, this remains distal nephron in nature. Most existing protocols reported to generate ureteric epithelium also commence with a short induction in CHIR to mimic canonical Wnt signalling followed by inhibitions of BMP activity. In most instances, this is followed by isolation of a specific cellular subset for subsequent culture without a surrounding stroma^16,17,18,19^. In this protocol, as well as that recently described by Shi et al^20^, all component cells within what is an heterogeneous differentiation were maintained. In this study, the resulting organoids patterned to distal nephrons while Shi et al embedded the entire spheroid culture into Matrigel^20^. Uniquely, we were able to seed single GATA3+ cells from these organoids into Matrigel to form iUE epithelial cysts, but these did not branch as was observed in Shi et al^20^.

Recent analyses of the developing human kidney suggested that the early nephron was not an aggregate of uniform progenitors but a spatially arranged set of distinct progenitors able to give rise to either podocyte, proximal nephron or distal nephron^49^. Indeed, Rinkevitch et al^50^ previously proposed the existence of segment-specific progenitors in mouse nephrons using lineage tracing. This predicted that *in vitro* you could induce the formation of a single specific progenitor for an individual nephron segment. Counter to this notion, we have generated a distal nephron progenitor that can adopt a ureteric tip / ureteric epithelium / collecting duct identity in response to external cues. This challenges the distinction between the origin of nephron and collecting duct. Indeed, the similarity in signature between nephric duct, ureteric epithelium and distal tubule, and the evidence of plasticity demonstrated in this study, suggests that this is a patterning continuum exquisitely poised to respond to surrounding signals. Ultimately, it will be the stability of this identity that will determine the utility of the cells generated.

Our hypothesis that the epithelial identity is regulated by the environment supports the data proposing distinct stromal signals defining nephron patterning^51^. Stromal-epithelial crosstalk has also been observed in adult tissues with epithelial identity changing in response to co-culture with a mismatched source of mesenchyme, including the adoption of an epithelial identity from another germ lineage^52^. Such epithelial transition is proposed to result from transdifferentiation of remaining adult epithelial progenitors. Indeed, Schutgens et al^53^ describe the presence of a TROY^+^ epithelial ‘stem cell’ within the collecting duct of the adult kidney and it is known that the relative ratio of Principal cells versus Intercalated cells can be altered in response to injury or toxicity, again suggesting prolonged epithelial plasticity. Of importance, this does suggest an ability to fine tune the epithelial cell state generated for the purposes of modelling disease in a specific nephron segment.

In summary, our modified human pluripotent stem cell differentiation protocol results in the formation of a plastic distalized renal epithelium within a surrounding stroma. For tissue engineering, there is a need to generate *bona fide* ureter and collecting duct. For disease modelling, there is a need to generate distinct regions of both the ureteric epithelium and the distal nephron. This method may provide appropriate cell types for the study of genetic conditions such as Gitelman syndrome, Bartter’s syndrome, Liddle’s syndrome, distal renal tubular acidosis and Autosomal recessive polycystic kidney disease. The protocol described here provides an opportunity to extend and refine further differentiation to each of these endpoints. It potentially also provides a model with which to further study the process of tubulogenesis during renal development.

## Experimental Procedures

### Pluripotent stem cell lines

1502.2^30^ and GATA3^mCherry+^ iPSC lines were previously reported^22^. RET ^tdTomato+^ was generated in the laboratory of Dr. Sanjay Jain as a part of the ReBuilding a Kidney consortium (https://www.rebuildingakidney.org/chaise/record/#2/Cell_Line:Reporter_Cell_Line/RID=Q-2CW0). All human iPSC lines were maintained and expanded at 37C, 5% CO2 and 5% O2 in Essential 8 medium (Thermo Fisher Scientific) on Matrigel-coated (Corning) plates with daily media changes and passaged every 3–4 days with EDTA in 1X PBS as previously described^54^. All the iPSC lines were tested for mycoplasma infection and the genomic integrity of iPSCs were also confirmed by molecular karyotyping using Infinium CoreExome-24 v1.1 SNP arrays (Illumina), performed by Victorian Clinical Genetics Services (Melbourne, Australia).

### Antibodies

The following primary antibodies were used for immunofluorescence at a concentration of 1:300: Goat polyclonal anti-GATA3 (Cat#AF2605; R&D Systems), Mouse monoclonal anti-KRT8 (Cat#AB115959; Abcam), Rabbit polyclonal anti-PAX2 (Cat#71-6000; Zymed Laboratories Inc.), Rat polyclonal anti-αPKC (Cat#sc-216; Santa Cruz), Rabbit monoclonal anti-GATA3 (Cat#5852; Cell Signalling Technology), Sheep polyclonal anti-NEPHRIN (Cat#AF4269; R&D Systems), Mouse monoclonal anti-E-CADHERIN (Cat#610181; BD Biosciences), Biotinylated Lotus tetragonolobus lectin (LTL) (Cat#B-1325; Vector Laboratories), Rabbit polyclonal anti-RFP (Cat#PM005; Medical & Biological Laboratories Co.), Goat polyclonal anti-LHX1 (Cat# sc-19341; Santa Cruz Biotechnology, Inc.), Mouse monoclonal anti-MES1/2/3 (Cat# 39795, ActiveMotif), Rabbit polyclonal anti-UMOD (Cat#BT-590; Biomedical Technologies Inc), Rabbit polyclonal SLC12A1 (Cat#18970-1-AP; Proteintech Group), Rabbit polyclonal SLC12A3 (Cat#LS-C662545; Sapphire Bioscience), Chicken polyclonal anti-GFP (Cat#ab13970; Abcam), Rabbit polyclonal anti-SIX2 (Cat#11562-1-AP; Proteintech Group), Rabbit polyclonal anti-UPK3A (Cat#HPA018415; Sigma-Aldrich), Mouse monoclonal anti-actin (ACTA2) (Cat#A2547; Sigma-Aldrich), Rabbit monoclonal anti-RET (Cat#3223, Cell Signalling Technology), Rabbit polyclonal anti-AQP2 (Cat#AB3274; Chemicon). Secondary antibodies conjugated with Alexa 488, 568 or 647 were used at 1:400 alongside with DAPI at 1:3000 (Life Technologies). Images were taken using a LSM 780 confocal microscope (Carl Zeiss).

### Quantitative RT-PCR analysis

Total RNA was extracted from cells using Purelink RNA Mini Kit (Cat# 12183020, ThermoFisher Scientific) as per manufacturer’s instructions. RNA was quantified using a NanoDrop 1000 spectrophotometer (ThermoFisher Scientific). Equal amounts of RNA were used to make cDNA using the GoScript reverse transcription system (Cat#A2800, Promega), following manufacturer’s instructions. Quantitative PCR were performed using GoTaq qPCR master mix (Cat#A6001, Promega). The reaction was conducted on an ABI Real-time PCR (Applied Biosystems) machine. Amplification was performed in triplicate in a MicroAmp Fast Optical 96-well optical reaction plate (Cat#4346906, Applied Biosystems, with 20 μl reaction samples. Primer sequences are listed in Table 1. cDNA levels of target genes were analysed using comparative Ct levels, normalized to glyceraldehyde-3-phosphate dehydrogenase and expressed as relative transcript abundance (ΔCt or 2−ΔCT), as well as fold change relative to control (ΔΔCT or 2–ΔΔCT)^55^. Error bars represent S.E.M. calculated from independent samples (wells or organoids or pooled organoids as stated); statistical significance was assessed using unpaired (two-tailed) Student’s t-test. P-values are represented in the Figures as *= P ≤ 0.05, **= P ≤ 0.01, ***= P ≤ 0.001 and ****= P ≤ 0.0001. Standard error of the mean is represented in error bars.

**Table 1.**
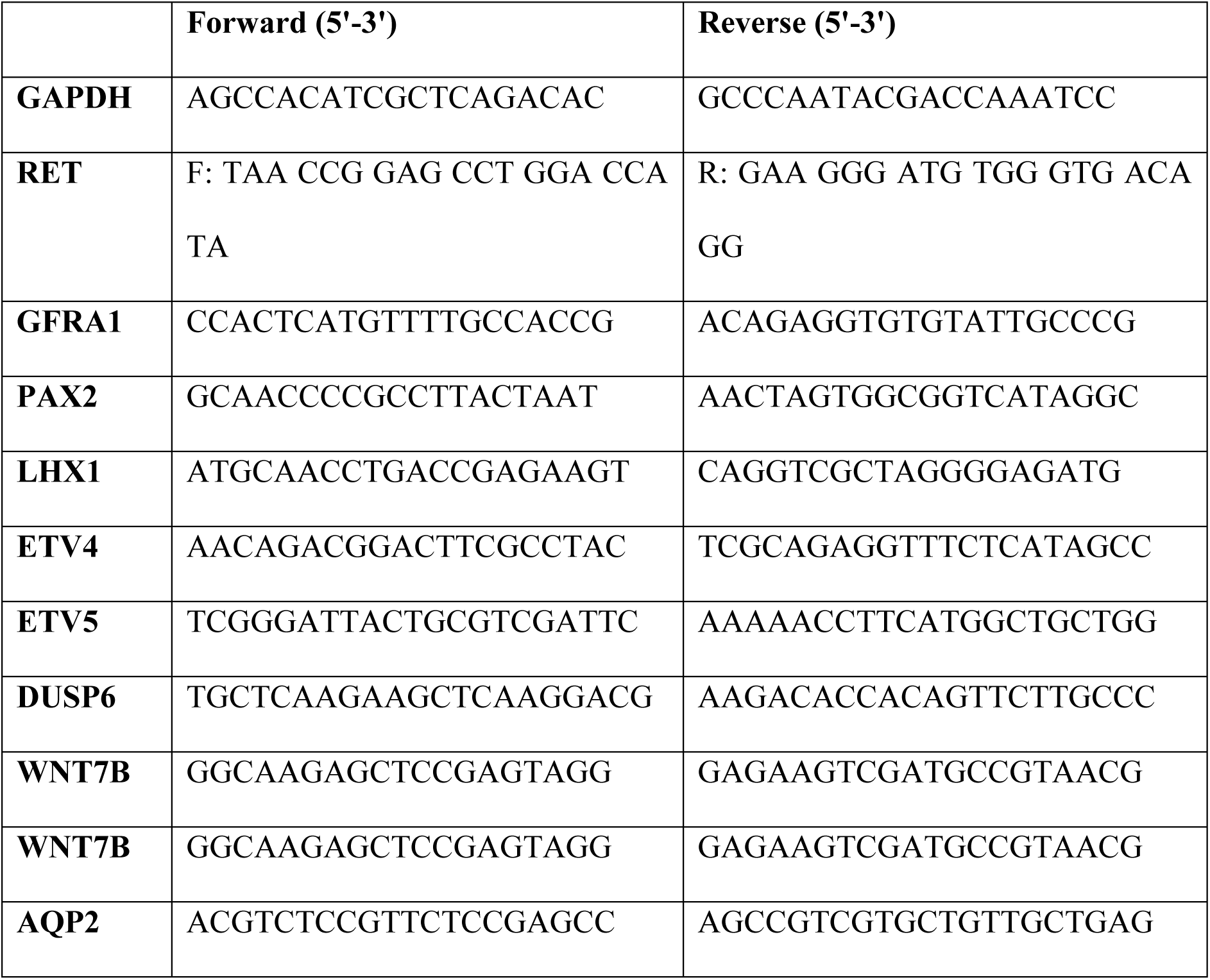
Primer sequences for QPCR.

### Flow cytometry

Kidney organoids were dissociated with 1:1 TrypLE/Accutase solution at 37°C for 10-15 min, with occasional pipetting until large clumps were no longer visible. Cells were then washed with FACS buffer (PBS 1X+ 2% FBS) before passing them through a 40-uM FACS tube cell strainer (Cat# 352235; Falcon). Flow cytometry was performed using a LSRFortessa Cell Analyzer (BD Biosciences). Data acquisition and analysis were performed using FACSDiva (BD Biosciences) and FlowLogic software (Inivai). Gating was performed on live cells based on forward and side-scatter analysis.

### Differentiation and kidney organoid generation

Briefly, undifferentiated iPSCs were dissociated using TrypLE-select enzyme (Cat# 12563029; ThermoFisher Scientific) and seeded onto Matrigel-coated (Cat#FAL354277; Corning) vessels at the cell density of 192 K cells/cm^2^ in Essential 8^TM^ medium with 5 μM Y27632 for 24 hours. The next day, cells were incubated with 4 to 6 μM CHIR99021 (Cat# 4423; Tocris Bioscience) in APEL 2 (Cat# 5275; StemCell Technologies) for 2 days. Then the medium was aspirated, and cells were treated with 200ng/mL FGF9 (Cat# 273-F9; R&D Systems), 1ug/mL heparin (Cat# H4784; Sigma Aldrich), 0.2uM ATRA (Cat#R2625; Sigma Aldrich), 0.2uM cis-RA (Cat#BML-GR101; Sapphire Biosciences), 100ng/ml GDNF (Cat#212-GD; R&D Systems) and 0.1uM LDN-193189 (Cat#04-0074; Stemgent) for five days, refreshing the media every two days. After seven days of differentiation, the cells were detached using Trypsin EDTA (0.05%) at 37°C for 3 minutes to make organoids according to Takasato *et al*.^29, 54^. Aggregates were made by centrifuging three times at 400xg for 3min and were gently picked up using a wide-bore pipette tip and were placed onto a transwell 0.4μm pore polyester membrane (Corning) in media containing 200ng/mL FGF9, 1ug/mL heparin, 0.2uM ATRA, 0.2uM cis-RA, 100ng/ml GDNF, 0.1uM LDN-193189 and 300ng/ml R-Spondin 1 (Cat#4645-RS, R&D Systems), refreshing the medium every two days until harvested.

### Spheroid culture assays

Organoids were dissociated with 1:1 Tryple/Accutase solution at 37°C for 10-15 min, with occasional pipetting until large clumps were no longer visible. Cell suspension was then washed with FACS buffer (PBS 1X+ 2% FBS) before passing them through a 40-lM FACS tube cell strainer (Falcon). Cells harboring a GATA3 promoter (driven by mCherry+) or RET (driven by tdTomato+) were resuspended in FACS buffer, and the mCherry+ and tdTomato+ fraction were isolated by fluorescent activated sorting. The sorted mCherry+ and/or tdTomato+ cells were resuspended in *Apel II* supplemented with 200ng/mL FGF9, 1ug/mL heparin, 0.2uM ATRA, 0.2uM cis-RA, 100ng/ml GDNF, 0.1uM LDN-193189 and 300ng/ml R-Spondin 1 (Cat#4645-RS, R&D Systems).

### Embryonic mouse recombination assay

Recombination assays were performed as previously described ^43, 44, 45^. Briefly, two different proportions of exogenous cells (5 and 10 % mCherry+ cells) FACS sorted from day 7+14 organoids, were recombined with dissociated embryonic kidneys ∼13.5 dpc. As a control, mouse embryonic dissociated kidney cells (∼13.5 dpc) were mixed with 10% iPSCs from the same parental line, and then cultured in the same way before being analysed. All recombination pellets were placed on a filter membrane and harvested after six days of culture for fixation and immunofluorescence.

### Grafting of eUBs into cultured kidney rudiments

Grafting assays were performed as described previously by Sallam et al.^47^, with modifications to structure isolation and environment evaluation. Briefly, mCherry+ ductal structures were manually dissected from day 7+14 GATA3^mCherry^ organoids, and then grafted into *ex fetu* mouse kidney rudiments obtained from Hoxb7-GFP+ transgenic mouse^56^. Human mCherry^+^ tissue was placed into near proximity of three different zones: MM, PWM as well as an injured Hoxb7^+^/GFP^+^ tree. Next, explants were monitored by live imaging of GFP+ (Hoxb7) and mCherry+ (GATA3) reporters within a humidified incubation chamber set at 37C and 5%0_2_ using a confocal ZEISS LSM 780, Carl Zeiss, Oberkochen, Germany. Finally, explants were harvested after three to four days for fixation and immunofluorescence.

### Single cell transcriptional profiling and data analysis

Three single cell RNA-sequencing datasets were generated in this study, one at 14 days of age (Day 7+7) and two at 21 days of age (D7+14) with RSPO1 added either at day 7 or day 14. Cells were dissociated following previously published methods (ref). Libraries were generated following the standard 10x Chromium Next GEM Single Cell 3ʹ Reagent Kits v3.1 protocol. Sequencing was performed using an Illumina Novoseq. 10x mRNA libraries were demultiplexed using CellRanger (3.1.0) to generate matrices of UMI counts per cell. All data were analysed using R (3.6.1) with the package Seurat (3.1.4) unless otherwise stated. Cells with mitochondrial content greater than 40%, number of UMIs per cell less than 8000 were removed to obtain filtered datasets. Data was log-normalised (factor 10,000) and scaled. The 21day datasets were integrated using Seurat’s anchor method. 30 component Principal Component Analysis (PCA) was performed using the 2000 most highly variable genes. Uniform Manifold Approximation and Projection (UMAP) were generated using PCA components. Seurat’s graph-based clustering approach was used to identify similar cell populations within the data, with resolutions of 0.25 and 0.15 used for 14 and 21 day old datasets respectively. Differential gene lists were generated using FindAllMarkers. Gene expression patterns were visualised using the FeaturePlot and DotPlot functions. The samples were independently annotated using the R package DevKidCC (v0.2.2). DevKidCC is a hierarchical set of machine-learning binary classifiers trained on a human fetal kidney reference dataset.

## Code and Data Availability

The data presented within this study is available within the Gene Expression Omnibus at the accession number GSE184634. Code is available at github.com/KidneyRegeneration/Kairath2021.

## Acknowledgements

This work was supported by the National Institutes of Health Rebuilding a Kidney consortium (UH3DK107344; U01DK107350), Dutch Kidney Foundation (RecordKID) and the National Health and Medical Research Council, Australia (GNT1156440). We acknowledge the Stafford Fox Medical Research Foundation MCRI genome editing facility for the generation of all pluripotent stem cell lines other than the Ret reporter line. For the latter, we are grateful to Dr. Bruce Conklin, Gladstone Institute, for the generous gift of the parent WTC11GCampf line that was used to engineer the Ret-reporter line. We thank the Alvin J. Siteman Cancer Center at Washington University School of Medicine and Barnes-Jewish Hospital in St. Louis, MO for the use of the Genome Engineering and Induced Pluripotent Stem Cell Center, which provided gene editing service for the RET^tdTomato^ iPSC reporter line. The Siteman Cancer Center is supported in part by an NCI Cancer Center Support Grant #P30 CA09184. MHL was an NHMRC Senior Principal Research Fellow (GNT1136085) and is supported by the Novo Nordisk Foundation Center for Stem Cell Medicine (<u>NNF21CC0073729</u>).

## Author contributions

Pamela Kairath^1^, Pei Xuan Er^1^, Sean Wilson^1^, Irene Ghobrial ^1^, Jessica M. Vanslambrouck^1^, Chen Y-H^2^, Miller S^2,3,^ Sanjay Jain^2^, Melissa H. Little^1,4,5#^

PK: conceptualisation, methodology, validation, formal analysis, investigation, writing, visualisation; PXE, IG: methodology, validation, investigation, manuscript review; SW: methodology, investigation, formal analysis, software, data curation, visualisation, writing, review and editing; JMV: methodology, investigation, resources, supervision, review and editing; CY-HM, SJ: resources, review and editing; MHL: Conceptualisation, formal analysis, writing, review and editing, visualisation, supervision, project administration, manuscript writing and submission, funding acquisition.

## Supplementary Figures

**Supplementary Figure 1.**
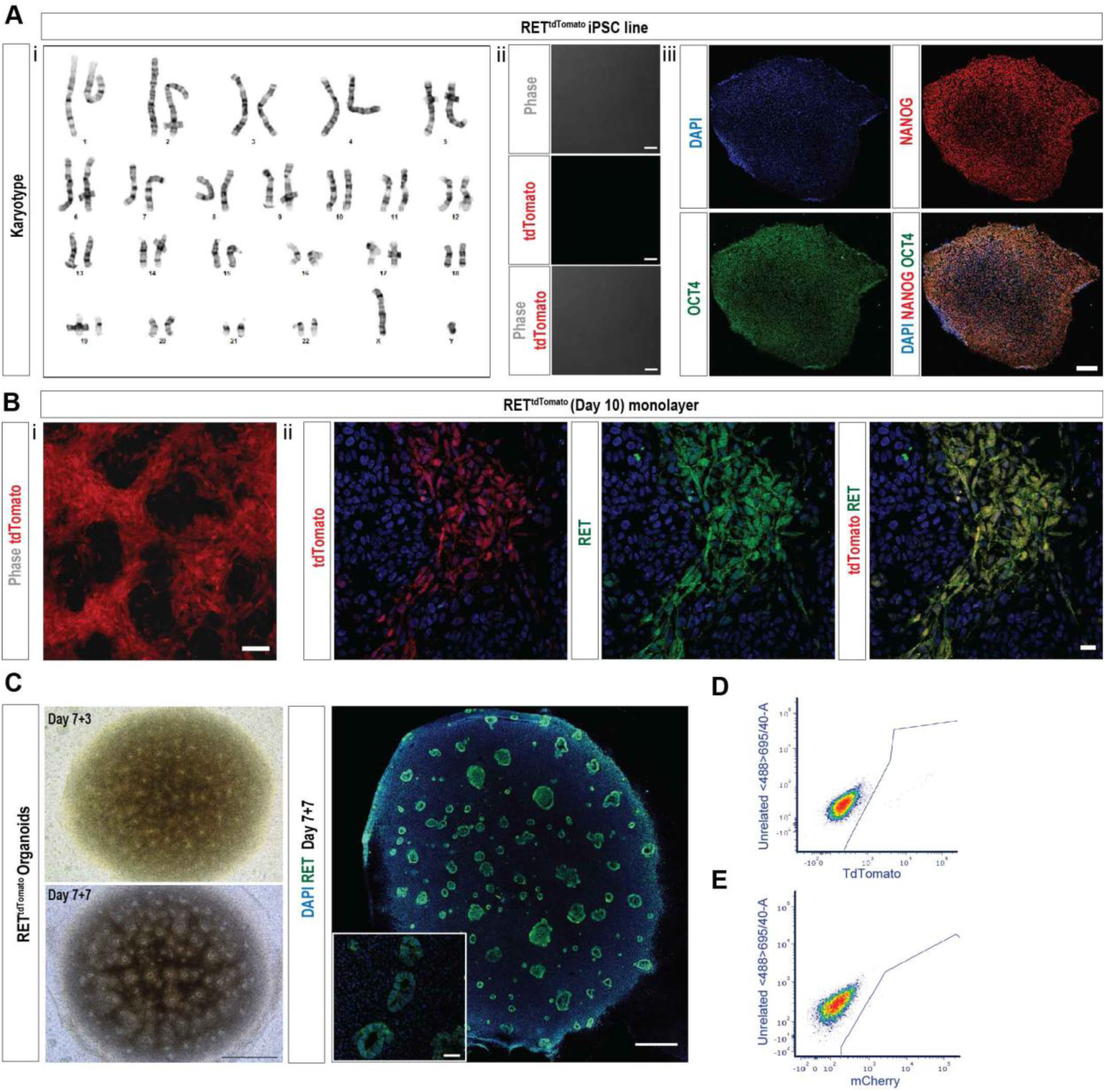
Generation, characterisation, and differentiation of the RET^tdTomato^ reporter line. **A.** Characterisation of RET^tdTomato^ reporter line including karyotype and immunostaining for pluripotency proteins NANOG and OCT4. Scale bar= 100uM. **B.** Day 10 monolayer culture of RET^tdTomato^ reporter line showing (i) native TdTomato expression and (ii) co-localisation of TdTomato and RET proteins via confocal immunofluorescence. Scale bar = 100 and 20 uM, respectively **C.** Brightfield (Day 7+5), immunostaining (Day 7+7), live fluorescence(i) and FACS-sorting(ii) of RET^tdTomato^ organoids. Scale bar: 500 uM.

**Supplementary Figure 3.**
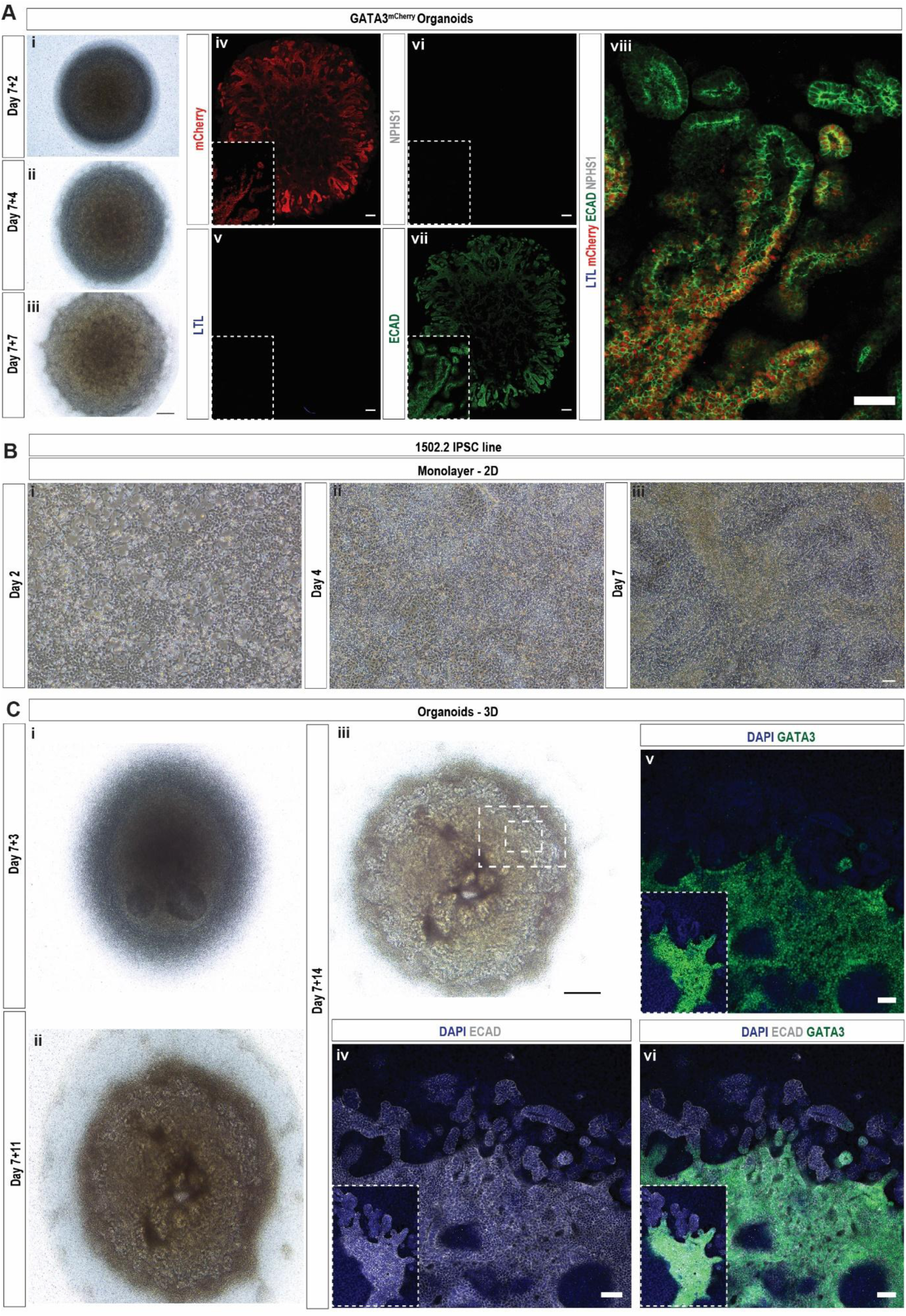
**A.** Time-course of organoids generated with the GATA3^mCherry^ reporter line. Brightfield and immunostaining against mCherry, LTL, ECAD, NPHS1 Scale bar= 500 and 50uM. **B.** Differentiation timecourse using a control human induced pluripotent stem cell line 1502.2. Scale bar = 100 uM. **C.** Characterisation of distalized organoids generated from control human induced pluripotent stem cell line 1502.2 showing formation of an extensive epithelial plexus including a GATA3^+^ECAD^+^ and GATA3^−^ECAD^+^ epithelial elements. Scale bar = 500uM.

**Supplementary Figure 4.**
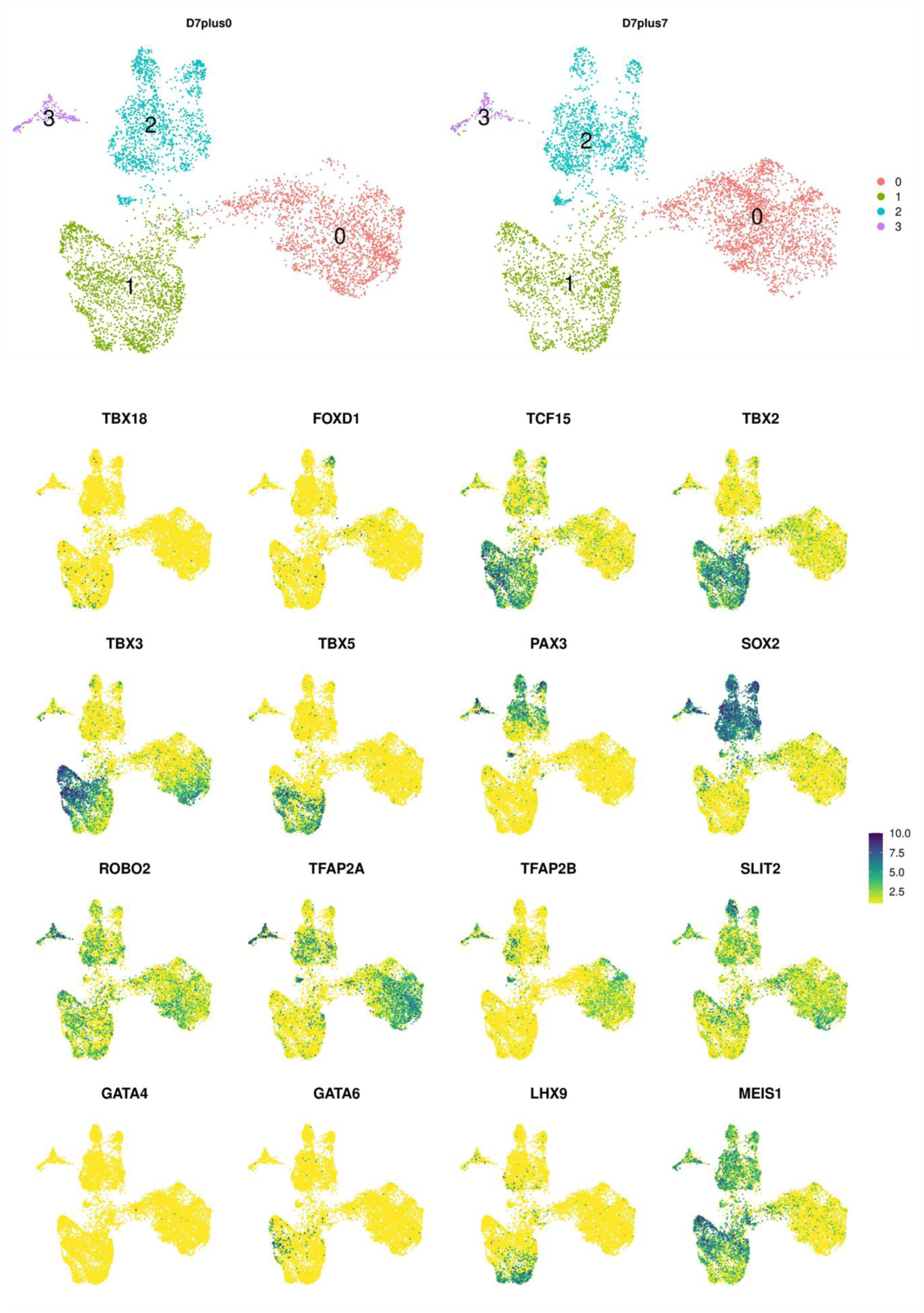
Feature plots of key genes in extended distalized organoids. Top: UMAP plot related to figure 4A showing the distribution of both integrated samples. Bottom: UMAP plot related to figure 4A showing various gene expression profiles.

**Supplementary Figure 5.**
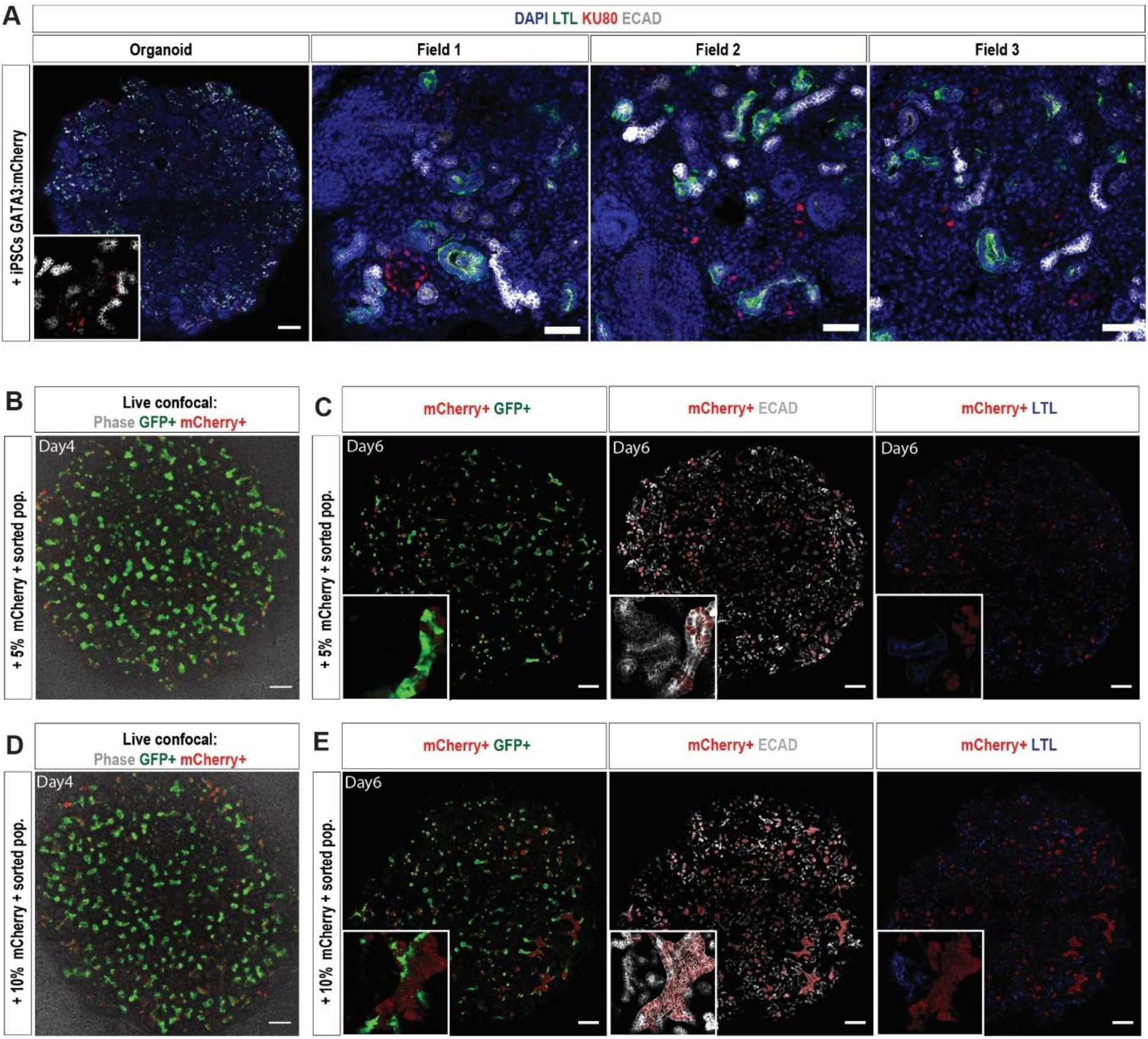
Analysis of recombination assays. **A.** Recombination between dissociated 12.5dpc mouse embryonic kidney and 5% undifferentiated induced pluripotent stem cells showing no integration into collecting duct segments. Human iPSC are detected using a human nuclear antibody (KU80, red). Staining for proximal tubules (LTL, green), distal epithelium / ureteric epithelium (ECAD, white) and nuclei (DAPI, blue) is also presented. Scale bars: 200 and 50 uM. Images representative of 2 independent aggregations. **B.** Image of entire reaggregation containing 5% sorted mCherry^+^ epithelium showing live imaging for GFP and mCherry. Scale bar: 200 uM. **C.** Immunostaining for GFP, ECAD (distal epithelium / UE) and LTL (proximal tubule marker) of organoid in B. Scale bars: 200 uM. **BC.** Images representative of two example aggregations. **D.** Image of entire reaggregation containing 5% sorted mCherry^+^ epithelium showing live imaging for GFP and mCherry. Scale bar: 200 uM**. E.** Immunostaining for GFP, ECAD (distal epithelium / UE) and LTL (proximal tubule marker) of organoid in D. Scale bar: 200 uM. **DE.** Images representative of two example aggregations.

## Supplementary Tables

**Supplementary Table 1.** DE genes for each cluster identified in tdTomato RET day 7 organoid data

**Supplementary Table 2.** DE gene lists for each subcluster within epithelial population of tdTomato RET day 7 organoid data

**Supplementary Table 3.** DE genes for each cluster identified in GATA3 mCherry day 7+14 organoids cultured with RSPO1

**Supplementary Table 4.** DE gene lists for epithelial subcluster within GATA3 mCherry day 7+14 organoids cultured with 7 or 14 days of RSPO1

