## Supplementary Table 1 for "*In vitro* plasticity between ureteric epithelial and distal nephron identity and maturity is controlled by extracellular signals"

DE genes for each cluster identified in tdTomato RET day 7 organoid data

Cluster 0

| p_val | avg_logFC | pct.1 | pct.2 | p_val_adj | cluster | gene |
|---|---|---|---|---|---|---|
| 3.8E-172 | 1.504956 | 0.584 | 0.052 | 1.7E-167 | 0 | PNOC |
| 0 | 1.30789 | 0.964 | 0.356 | 0 | 0 | WFDC2 |
| 0 | 1.243988 | 0.993 | 0.37 | 0 | 0 | LHX1 |
| 3.3E-274 | 1.094687 | 0.863 | 0.28 | 1.5E-269 | 0 | HBD |
| 9.8E-118 | 1.090536 | 0.493 | 0.034 | 4.4E-113 | 0 | SST |
| 0 | 0.98465 | 0.867 | 0.064 | 0 | 0 | EPCAM |
| 0 | 0.960282 | 0.854 | 0.12 | 0 | 0 | PAX2 |
| 2.5E-142 | 0.811448 | 0.548 | 0.053 | 1.1E-137 | 0 | SPP1 |
| 3.5E-164 | 0.793102 | 0.55 | 0.023 | 1.6E-159 | 0 | MAL |
| 0 | 0.781149 | 0.984 | 0.654 | 0 | 0 | SPINT2 |
| 7E-304 | 0.779209 | 0.834 | 0.129 | 3E-299 | 0 | KRT19 |
| 2.5E-157 | 0.74869 | 0.59 | 0.051 | 1.1E-152 | 0 | ALDH1A1 |
| 0 | 0.700144 | 0.811 | 0.065 | 0 | 0 | PAX8 |
| 0 | 0.695914 | 0.949 | 0.567 | 0 | 0 | KRT18 |
| 2.8E-241 | 0.659116 | 0.852 | 0.366 | 1.2E-236 | 0 | ATP1B1 |
| 3.4203573: | 0.618442 | 0.902 | 0.505 | 1.5E-225 | 0 | SAT1 |
| 0 | 0.61714 | 0.954 | 0.523 | 0 | 0 | TM7SF2 |
| 1E-303 | 0.603587 | 0.81 | 0.127 | 5E-299 | 0 | BMP7 |
| 0 | 0.601241 | 0.91 | 0.345 | 0 | 0 | TSTD1 |
| 0 | 0.593419 | 0.942 | 0.426 | 0 | 0 | DMKN |
| 1.6E-198 | 0.581984 | 0.918 | 0.58 | 7.2E-194 | 0 | APOE |
| 5.4E-232 | 0.576397 | 0.68 | 0.03 | 2.4E-227 | 0 | PCSK1N |
| 8.9E-285 | 0.549528 | 0.914 | 0.415 | 4.0319078: | 0 | IRX3 |
| 4.4E-213 | 0.548254 | 0.642 | 0.048 | 2E-208 | 0 | TFAP2A |
| 2E-304 | 0.54454 | 0.996 | 0.934 | 1E-299 | 0 | ITM2C |
| 4E-291 | 0.535222 | 0.797 | 0.126 | 1.8E-286 | 0 | UCP2 |
| 0 | 0.535093 | 0.97 | 0.689 | 0 | 0 | PCBD1 |
| 3.1E-197 | 0.532344 | 0.703 | 0.13 | 1.4E-192 | 0 | RDH10 |
| 1.5E-213 | 0.527099 | 0.741 | 0.16 | 6.9E-209 | 0 | BCAM |
| 1.2E-158 | 0.511169 | 0.982 | 0.85 | 5.3E-154 | 0 | GPC3 |
| 9.85E-51 | 0.50982 | 0.646 | 0.398 | 4.44E-46 | 0 | IGFBP7 |
| 2.31E-95 | 0.505538 | 0.621 | 0.201 | 1.0411405! | 0 | RGS5 |

Supplementary Table 1

### DE genes for each cluster identified in tdTomato RET day 7 organoid data

Cluster 1

| p_val | avg_logFC | pct.1 | pct.2 | p_val_adj | cluster | gene |
|---|---|---|---|---|---|---|
| 8.7E-261 | 0.743816 | 1 | 0.976 | 3.9E-256 | 1 | VIM |
| 7.9E-144 | 0.616512 | 0.974 | 0.838 | 3.5E-139 | 1 | SFRP1 |
| 2.8E-159 | 0.601933 | 0.933 | 0.729 | 1.3E-154 | 1 | CPE |
| 1.4E-201 | 0.59935 | 0.969 | 0.743 | 6.5E-197 | 1 | ALDH1A2 |
| 1.5E-226 | 0.575083 | 0.998 | 0.926 | 6.9E-222 | 1 | PTN |
| 6.5E-127 | 0.555869 | 0.933 | 0.69 | 2.9E-122 | 1 | NEFL |
| 1.0932899! | 0.531693 | 0.966 | 0.813 | 4.9E-136 | 1 | PLAT |
| 2.8E-142 | 0.519213 | 0.982 | 0.85 | 1.3E-137 | 1 | NEFM |
| 7.6E-142 | 0.504017 | 0.932 | 0.655 | 3.4E-137 | 1 | POSTN |

Supplementary Table 1

DE genes for each cluster identified in tdTomato RET day 7 organoid data

Cluster 2

| p_val | avg_logFC | pct.1 | pct.2 | p_val_adj | cluster | gene |
|---|---|---|---|---|---|---|
| 7.5E-266 | 1.553228 | 0.999 | 0.926 | 3.4E-261 | 2 | HIST1H4C |
| 0 | 1.053406 | 0.988 | 0.551 | 0 | 2 | TYMS |
| 0 | 0.99535 | 0.975 | 0.518 | 0 | 2 | KIAA0101 |
| 0 | 0.85843 | 1 | 0.979 | 0 | 2 | TUBA1B |
| 4.4E-269 | 0.809835 | 0.993 | 0.87 | 2E-264 | 2 | DUT |
| 8.5E-142 | 0.805589 | 0.741 | 0.299 | 3.8E-137 | 2 | HIST1H1D |
| 6.1E-132 | 0.707477 | 0.688 | 0.173 | 2.8E-127 | 2 | HIST1H1B |
| 5.5E-243 | 0.706357 | 0.961 | 0.602 | 2.5E-238 | 2 | PCNA |
| 0 | 0.703443 | 1 | 0.963 | 0 | 2 | H2AFZ |
| 2.4E-249 | 0.702151 | 0.983 | 0.744 | 1.1E-244 | 2 | CENPV |
| 2.9E-226 | 0.676927 | 0.929 | 0.473 | 1.3E-221 | 2 | GINS2 |
| 1.1E-164 | 0.646006 | 0.776 | 0.249 | 5.0226723! | 2 | RRM2 |
| 4.8E-203 | 0.63603 | 0.874 | 0.338 | 2.2E-198 | 2 | CLSPN |
| 7.2548158: | 0.62581 | 0.999 | 0.931 | 3.3E-285 | 2 | RANBP1 |
| 7.2E-275 | 0.622924 | 0.989 | 0.754 | 3.2586783! | 2 | TMSB15A |
| 0 | 0.618399 | 0.999 | 0.956 | 0 | 2 | NASP |
| 1.2509117 | 0.576227 | 0.896 | 0.386 | 5.6E-196 | 2 | CENPU |
| 1.3E-224 | 0.568158 | 0.982 | 0.766 | 5.8120415! | 2 | MCM7 |
| 5.6E-208 | 0.556818 | 0.925 | 0.44 | 2.5E-203 | 2 | ORC6 |
| 2.6E-177 | 0.548104 | 0.884 | 0.469 | 1.2E-172 | 2 | UBE2T |
| 8.5E-207 | 0.545577 | 0.957 | 0.627 | 3.8E-202 | 2 | HELLS |
| 1.0533486: | 0.545522 | 0.933 | 0.462 | 4.7E-206 | 2 | MAD2L1 |
| 1.6E-188 | 0.536079 | 0.899 | 0.456 | 7.3E-184 | 2 | ZWINT |
| 9.0190423! | 0.532031 | 0.925 | 0.602 | 4.1E-185 | 2 | RNASEH2A |
| 2.4E-157 | 0.524803 | 0.803 | 0.307 | 1.1E-152 | 2 | ATAD2 |
| 2.6E-154 | 0.523789 | 0.74 | 0.202 | 1.2E-149 | 2 | MYBL2 |
| 3.3E-238 | 0.516808 | 0.988 | 0.754 | 1.5E-233 | 2 | CKS1B |
| 5.6E-261 | 0.514077 | 1 | 0.978 | 2.5E-256 | 2 | HMGN2 |
| 2.3E-175 | 0.510253 | 0.924 | 0.552 | 1.0308652: | 2 | GMNN |

Supplementary Table 1

### DE genes for each cluster identified in tdTomato RET day 7 organoid data

Cluster 3

| p_val | avg_logFC | pct.1 | pct.2 | p_val_adj | cluster | gene |
|---|---|---|---|---|---|---|
| 4.9E-249 | 2.010124 | 0.993 | 0.455 | 2.2E-244 | 3 | UBE2C |
| 0 | 1.948564 | 0.998 | 0.554 | 4E-307 | 3 | CENPF |
| 7.2E-279 | 1.782184 | 0.998 | 0.4 | 3.2E-274 | 3 | TOP2A |
| 1.3E-234 | 1.706476 | 0.993 | 0.434 | 5.7324432! | 3 | CCNB1 |
| 2.9961299: | 1.659711 | 1 | 0.578 | 1E-305 | 3 | PTTG1 |
| 7.7E-212 | 1.636978 | 0.998 | 0.745 | 3.5E-207 | 3 | KPNA2 |
| 0 | 1.570465 | 1 | 0.821 | 0 | 3 | HMGB2 |
| 4.6E-243 | 1.556141 | 0.993 | 0.443 | 2.1E-238 | 3 | CCNB2 |
| 8.5E-208 | 1.444391 | 0.985 | 0.303 | 3.8E-203 | 3 | CDC20 |
| 2.4E-246 | 1.433348 | 0.996 | 0.359 | 1.1E-241 | 3 | TPX2 |
| 9.8E-203 | 1.410487 | 0.985 | 0.367 | 4.4E-198 | 3 | MKI67 |
| 6.1E-252 | 1.383463 | 0.998 | 0.867 | 2.7E-247 | 3 | TUBB4B |
| 6.5E-184 | 1.381144 | 0.969 | 0.338 | 2.9E-179 | 3 | ASPM |
| 2.4E-152 | 1.371541 | 0.958 | 0.521 | 1.1E-147 | 3 | TUBA1C |
| 8.9E-225 | 1.351948 | 0.996 | 0.729 | 4.0267997 | 3 | UBE2S |
| 1.7E-222 | 1.330256 | 0.989 | 0.292 | 7.9E-218 | 3 | DLGAP5 |
| 9.2E-194 | 1.32012 | 0.982 | 0.562 | 4.1E-189 | 3 | ARL6IP1 |
| 3.0218625 | 1.314026 | 0.98 | 0.667 | 1.4E-195 | 3 | CKS2 |
| 1.9E-172 | 1.270942 | 0.956 | 0.29 | 8.7E-168 | 3 | CENPE |
| 2.2E-204 | 1.2464 | 0.987 | 0.389 | 9.8369898 | 3 | NUSAP1 |
| 5.1E-237 | 1.227633 | 0.989 | 0.435 | 2.3E-232 | 3 | BIRC5 |
| 1.1E-161 | 1.100218 | 0.958 | 0.303 | 4.8E-157 | 3 | CDKN3 |
| 4.1E-158 | 1.099579 | 0.929 | 0.173 | 1.9E-153 | 3 | PLK1 |
| 1.9434211! | 1.095116 | 0.998 | 0.778 | 8.8E-256 | 3 | CKS1B |
| 2E-132 | 1.075076 | 0.905 | 0.371 | 8.9E-128 | 3 | CDK1 |
| 6.7E-158 | 1.049949 | 0.945 | 0.273 | 3E-153 | 3 | SGOL2 |
| 2.4E-139 | 1.012237 | 0.905 | 0.153 | 1.1E-134 | 3 | CENPA |
| 4.9E-142 | 1.004376 | 0.927 | 0.306 | 2.2E-137 | 3 | PRC1 |
| 1E-187 | 0.995912 | 0.969 | 0.332 | 4.6E-183 | 3 | AURKB |
| 1.1E-143 | 0.995834 | 0.951 | 0.466 | 5.1E-139 | 3 | CKAP2 |
| 1.4E-147 | 0.99497 | 0.923 | 0.292 | 6.4E-143 | 3 | NDC80 |
| 3.3E-142 | 0.971788 | 0.914 | 0.276 | 1.5E-137 | 3 | FAM64A |
| 1.4819665 | 0.951814 | 0.971 | 0.574 | 6.7E-156 | 3 | H2AFX |
| 3E-158 | 0.949732 | 0.94 | 0.256 | 1.4E-153 | 3 | KIF2C |
| 4.4E-132 | 0.93936 | 0.905 | 0.219 | 2E-127 | 3 | HMMR |
| 2.1E-188 | 0.937223 | 0.985 | 0.599 | 9.4E-184 | 3 | SMC4 |
| 2.7E-148 | 0.923378 | 0.934 | 0.257 | 1.2E-143 | 3 | GTSE1 |
| 1.4E-157 | 0.917176 | 0.978 | 0.571 | 6.5E-153 | 3 | MIS18BP1 |
| 1.3E-153 | 0.910224 | 0.932 | 0.258 | 6E-149 | 3 | PSRC1 |
| 5.8E-112 | 0.905783 | 0.839 | 0.161 | 2.6E-107 | 3 | AURKA |
| 4.1E-135 | 0.891357 | 0.936 | 0.414 | 1.8628475 | 3 | KIF20B |
| 5.7E-128 | 0.872828 | 0.892 | 0.247 | 2.6E-123 | 3 | CCNA2 |
| 2.8E-152 | 0.869715 | 0.949 | 0.312 | 1.3E-147 | 3 | TACC3 |
| 4E-155 | 0.868836 | 0.978 | 0.702 | 1.7913397 | 3 | BUB3 |
| 3.7E-127 | 0.858967 | 0.872 | 0.22 | 1.7E-122 | 3 | TTK |
| 2.2E-179 | 0.851615 | 0.982 | 0.509 | 1E-174 | 3 | MAD2L1 |

| | | | | | | |
|---|---|---|---|---|---|---|
| 1.1E-113 | 0.850274 | 0.85 | 0.15 | 4.8E-109 | 3 | NEK2 |
| 2.6E-128 | 0.849564 | 0.898 | 0.221 | 1.2E-123 | 3 | CDCA3 |
| 0 | 0.839198 | 1 | 0.982 | 0 | 3 | TUBA1B |
| 5.7E-118 | 0.802881 | 0.859 | 0.155 | 2.6E-113 | 3 | KIF20A |
| 2.2E-86 | 0.802488 | 0.737 | 0.105 | 9.93E-82 | 3 | PIF1 |
| 8.9E-182 | 0.794225 | 1 | 0.965 | 4E-177 | 3 | NUCKS1 |
| 1.7E-153 | 0.791319 | 0.982 | 0.72 | 7.8E-149 | 3 | SKA2 |
| 4.8E-132 | 0.790932 | 0.914 | 0.288 | 2.2E-127 | 3 | KIFC1 |
| 5.3E-128 | 0.79052 | 0.905 | 0.263 | 2.4E-123 | 3 | NUF2 |
| 2.05896339 | 0.7855 | 0.859 | 0.186 | 9.3E-116 | 3 | CDCA8 |
| 3.8E-108 | 0.782096 | 0.839 | 0.174 | 1.7E-103 | 3 | DEPDC1 |
| 3.9E-128 | 0.780721 | 0.991 | 0.929 | 1.7E-123 | 3 | HN1 |
| 1E-112 | 0.775653 | 0.861 | 0.214 | 4.7E-108 | 3 | TROAP |
| 1.9E-112 | 0.77346 | 0.874 | 0.304 | 8.4E-108 | 3 | KNSTRN |
| 1E-116 | 0.771124 | 0.885 | 0.324 | 4.7E-112 | 3 | KIF11 |
| 1.9239841 | 0.767579 | 0.857 | 0.221 | 8.7E-106 | 3 | KIF23 |
| 6.7E-99 | 0.752938 | 0.859 | 0.256 | 3.01E-94 | 3 | MXD3 |
| 2.9E-124 | 0.738749 | 0.912 | 0.308 | 1.3E-119 | 3 | NCAPG |
| 4.8E-117 | 0.731621 | 0.874 | 0.249 | 2.2E-112 | 3 | ECT2 |
| 4.1E-113 | 0.722313 | 0.839 | 0.159 | 1.9E-108 | 3 | BUB1 |
| 6.8E-159 | 0.719781 | 0.993 | 0.865 | 3.1E-154 | 3 | HMGB3 |
| 4.07E-96 | 0.706338 | 0.843 | 0.234 | 1.83E-91 | 3 | CASC5 |
| 2.6E-123 | 0.697603 | 0.934 | 0.528 | 1.2E-118 | 3 | KIF22 |
| 1.5E-105 | 0.689501 | 0.837 | 0.193 | 6.7E-101 | 3 | CKAP2L |
| 1.7E-184 | 0.686754 | 1 | 0.976 | 7.7937908 | 3 | CALM2 |
| 1.4E-244 | 0.68117 | 1 | 0.981 | 6.4888882 | 3 | HMGN2 |
| 1.27E-87 | 0.681087 | 0.775 | 0.146 | 5.72E-83 | 3 | KIF14 |
| 9.21E-95 | 0.667268 | 0.826 | 0.215 | 4.1518052 | 3 | KIF4A |
| 1.9603261 | 0.660157 | 1 | 0.967 | 8.8E-286 | 3 | H2AFZ |
| 9.84E-98 | 0.6505 | 0.898 | 0.515 | 4.43E-93 | 3 | CKAP5 |
| 1E-103 | 0.645618 | 0.859 | 0.3 | 4.7E-99 | 3 | SGOL1 |
| 2.2E-109 | 0.644005 | 0.956 | 0.652 | 1E-104 | 3 | LMNB1 |
| 4.2E-123 | 0.639619 | 0.969 | 0.753 | 1.9E-118 | 3 | DTYMK |
| 7.3E-129 | 0.63754 | 0.982 | 0.786 | 3.3E-124 | 3 | ANP32E |
| 1.31E-84 | 0.629003 | 0.773 | 0.214 | 5.9229261 | 3 | ARHGAP11A |
| 9.98E-91 | 0.628487 | 0.797 | 0.179 | 4.5E-86 | 3 | KIF18A |
| 8.3E-119 | 0.611645 | 0.98 | 0.764 | 3.7E-114 | 3 | RAD21 |
| 2.65E-97 | 0.606794 | 0.85 | 0.296 | 1.2E-92 | 3 | SPC25 |
| 3.8E-94 | 0.603002 | 0.817 | 0.231 | 1.71E-89 | 3 | BUB1B |
| 4.19E-78 | 0.59954 | 0.865 | 0.503 | 1.89E-73 | 3 | DBF4 |
| 3.01E-83 | 0.598425 | 0.762 | 0.191 | 1.36E-78 | 3 | HJURP |
| 3.5E-131 | 0.596056 | 1 | 0.956 | 1.6E-126 | 3 | H1FX |
| 2.16397929 | 0.595969 | 0.704 | 0.434 | 9.75E-26 | 3 | LGALS1 |
| 1.8E-86 | 0.594875 | 0.775 | 0.18 | 8.13E-82 | 3 | CDCA2 |
| 2.5791423 | 0.586169 | 0.823 | 0.274 | 1.16E-85 | 3 | KIF15 |
| 1.21E-78 | 0.58326 | 0.848 | 0.415 | 5.46E-74 | 3 | ENDOG |
| 7.4202741 | 0.580796 | 0.914 | 0.511 | 3.34E-95 | 3 | UBE2T |
| 2.7E-106 | 0.576531 | 0.96 | 0.758 | 1.2E-101 | 3 | TMPO |
| 2.39E-94 | 0.556889 | 0.989 | 0.919 | 1.08E-89 | 3 | HP1BP3 |
| 2.9E-294 | 0.554236 | 1 | 0.997 | 1.3E-289 | 3 | HMGB1 |

| | | | | | | |
|---|---|---|---|---|---|---|
| 2.91E-88 | 0.551678 | 0.786 | 0.189 | 1.31E-83 | 3 | RACGAP1 |
| 1.74E-89 | 0.548969 | 0.905 | 0.553 | 7.83E-85 | 3 | MZT1 |
| 1.13E-78 | 0.539432 | 0.781 | 0.268 | 5.11E-74 | 3 | PHF19 |
| 3.5E-77 | 0.537214 | 0.773 | 0.254 | 1.58E-72 | 3 | GPSM2 |
| 2.09E-84 | 0.535038 | 0.744 | 0.154 | 9.4256685[illegible] | 3 | CDC25C |
| 7.5E-125 | 0.533713 | 0.978 | 0.65 | 3.3755957[illegible] | 3 | CENPW |
| 1.8E-212 | 0.526222 | 1 | 0.995 | 7.9E-208 | 3 | TUBB |
| 1.7E-132 | 0.52373 | 0.998 | 0.94 | 7.7E-128 | 3 | H2AFV |
| 3.08E-35 | 0.51584 | 0.558 | 0.255 | 1.3879069[illegible] | 3 | PRAC2 |
| 1.76E-87 | 0.515032 | 1 | 0.986 | 7.93E-83 | 3 | SFPQ |
| 2.09E-61 | 0.513711 | 0.713 | 0.271 | 9.41E-57 | 3 | TUBB6 |
| 6.52E-71 | 0.508166 | 0.682 | 0.152 | 2.94E-66 | 3 | CEP55 |
| 1.37E-88 | 0.505898 | 0.956 | 0.718 | 6.18E-84 | 3 | LBR |
| 1.35E-77 | 0.505386 | 0.797 | 0.335 | 6.09E-73 | 3 | NCAPD2 |
| 9.68E-64 | 0.504514 | 0.823 | 0.491 | 4.36E-59 | 3 | CNTRL |
| 9.9E-111 | 0.501312 | 0.98 | 0.78 | 4.5E-106 | 3 | TMSB15A |

Supplementary Table 1

### DE genes for each cluster identified in tdTomato RET day 7 organoid data

Cluster 4

| p_val | avg_logFC | pct.1 | pct.2 | p_val_adj | cluster | gene |
|---|---|---|---|---|---|---|
| 5.48539968 | 1.040599 | 0.954 | 0.888 | 2.47E-85 | 4 | DNM3OS |
| 3.9E-102 | 1.028768 | 0.975 | 0.957 | 1.77E-97 | 4 | WSB1 |
| 8.3E-105 | 1.027624 | 0.992 | 0.987 | 3.7416835 | 4 | DDX17 |
| 7.6E-61 | 1.011962 | 0.83 | 0.797 | 3.42E-56 | 4 | TIA1 |
| 2.54E-89 | 1.001693 | 0.949 | 0.847 | 1.14E-84 | 4 | PDGFRA |
| 1.7706196 | 0.990064 | 0.921 | 0.899 | 7.98E-66 | 4 | NKTR |
| 3.31E-82 | 0.95155 | 0.98 | 0.929 | 1.49E-77 | 4 | COL3A1 |
| 9.11E-32 | 0.938578 | 0.65 | 0.602 | 4.1E-27 | 4 | PCDH9 |
| 1.46E-45 | 0.881071 | 0.835 | 0.785 | 6.58E-41 | 4 | TSHZ2 |
| 2.13E-26 | 0.87544 | 0.718 | 0.795 | 9.62E-22 | 4 | NEAT1 |
| 4.73E-43 | 0.841829 | 0.726 | 0.716 | 2.13E-38 | 4 | CCNL2 |
| 8.22E-52 | 0.841538 | 0.827 | 0.797 | 3.71E-47 | 4 | TBX2 |
| 4.15E-34 | 0.836271 | 0.871 | 0.889 | 1.87E-29 | 4 | IGF2 |
| 7.46E-38 | 0.83131 | 0.789 | 0.703 | 3.36E-33 | 4 | POSTN |
| 2.21E-73 | 0.815767 | 1 | 0.998 | 9.97E-69 | 4 | MT-ATP6 |
| 4.2E-29 | 0.806754 | 0.424 | 0.249 | 1.89E-24 | 4 | TENM1 |
| 1.48E-47 | 0.800121 | 0.863 | 0.85 | 6.67E-43 | 4 | SOX11 |
| 8.56E-39 | 0.791352 | 0.744 | 0.755 | 3.86E-34 | 4 | MGEA5 |
| 2.1687517 | 0.790073 | 0.863 | 0.907 | 9.77E-36 | 4 | ZCCHC11 |
| 6E-35 | 0.787684 | 0.695 | 0.726 | 2.7020785 | 4 | RBM5 |
| 1.61E-22 | 0.776542 | 0.746 | 0.842 | 7.24E-18 | 4 | VCAN |
| 1.99E-33 | 0.772583 | 0.84 | 0.812 | 8.97E-29 | 4 | NTRK2 |
| 5.58E-45 | 0.762527 | 0.985 | 0.97 | 2.5150932 | 4 | KCNQ1OT1 |
| 1.03E-32 | 0.752228 | 0.723 | 0.754 | 4.64E-28 | 4 | CCDC14 |
| 1.34E-35 | 0.74992 | 0.754 | 0.786 | 6.05E-31 | 4 | BAZ2B |
| 5.3E-28 | 0.748179 | 0.642 | 0.675 | 2.39E-23 | 4 | DST |
| 8.17E-37 | 0.742243 | 0.721 | 0.726 | 3.68E-32 | 4 | COL6A1 |
| 6.3E-34 | 0.73991 | 0.652 | 0.628 | 2.84E-29 | 4 | DDR2 |
| 3.11E-36 | 0.729065 | 0.723 | 0.743 | 1.4E-31 | 4 | MARCH6 |
| 3.42E-64 | 0.728021 | 0.992 | 0.992 | 1.54E-59 | 4 | MTRNR2L12 |
| 7.51E-37 | 0.727307 | 0.64 | 0.578 | 3.39E-32 | 4 | MLXIP |
| 2.55E-64 | 0.727196 | 1 | 0.997 | 1.15E-59 | 4 | MT-CO1 |
| 1E-52 | 0.720681 | 0.995 | 0.997 | 4.51E-48 | 4 | MT-ND1 |
| 7.39E-31 | 0.719282 | 0.797 | 0.856 | 3.33E-26 | 4 | ZNF292 |
| 1.7E-25 | 0.719122 | 0.678 | 0.719 | 7.67E-21 | 4 | SCUBE3 |
| 8.9E-19 | 0.716593 | 0.779 | 0.878 | 4.01E-14 | 4 | PBX1 |
| 1.93E-33 | 0.714129 | 0.728 | 0.75 | 8.72E-29 | 4 | ZMYM2 |
| 1.22E-17 | 0.7137 | 0.447 | 0.485 | 5.5E-13 | 4 | FSIP2 |
| 9.56E-44 | 0.71297 | 0.868 | 0.887 | 4.31E-39 | 4 | COL6A2 |
| 1.2E-34 | 0.708053 | 0.584 | 0.523 | 5.3974740 | 4 | MRC2 |
| 8.38E-45 | 0.700679 | 0.886 | 0.917 | 3.7778411 | 4 | CHD9 |
| 5.4824126 | 0.700469 | 0.66 | 0.697 | 2.47E-25 | 4 | ARHGAP5 |
| 3.8687740 | 0.699689 | 0.901 | 0.942 | 1.74E-35 | 4 | SRSF11 |
| 2.26E-24 | 0.698721 | 0.792 | 0.837 | 1.02E-19 | 4 | FN1 |
| 2.23E-31 | 0.697094 | 0.637 | 0.628 | 1.01E-26 | 4 | NF1 |
| 1.68E-35 | 0.695284 | 0.647 | 0.609 | 7.58E-31 | 4 | TNRC6A |

| | | | | | | |
|---|---|---|---|---|---|---|
| 9.07E-25 | 0.691611 | 0.655 | 0.703 | 4.0872458[illegible] | 4 | FBN1 |
| 2.37E-25 | 0.686508 | 0.632 | 0.649 | 1.0668770[illegible] | 4 | AMOT |
| 1.86E-35 | 0.685554 | 0.782 | 0.771 | 8.38E-31 | 4 | CDH11 |
| 1.75E-31 | 0.685447 | 0.751 | 0.789 | 7.87E-27 | 4 | NFIB |
| 4.13E-29 | 0.685061 | 0.668 | 0.688 | 1.86E-24 | 4 | MAP3K1 |
| 2.02E-27 | 0.681068 | 0.637 | 0.671 | 9.09E-23 | 4 | ZNF518A |
| 5.33E-38 | 0.679316 | 0.906 | 0.938 | 2.4E-33 | 4 | ATRX |
| 2.79E-24 | 0.676782 | 0.579 | 0.628 | 1.26E-19 | 4 | PRTG |
| 4.69E-39 | 0.676203 | 0.901 | 0.952 | 2.12E-34 | 4 | ARGLU1 |
| 5.99E-26 | 0.672747 | 0.822 | 0.892 | 2.7E-21 | 4 | AKAP9 |
| 5.2863484[illegible] | 0.669256 | 0.784 | 0.852 | 2.38E-25 | 4 | MACF1 |
| 8.11E-68 | 0.663327 | 1 | 0.998 | 3.65E-63 | 4 | MT-CYB |
| 4.08E-24 | 0.662662 | 0.561 | 0.55 | 1.84E-19 | 4 | EBF3 |
| 2.94E-61 | 0.662285 | 1 | 0.998 | 1.33E-56 | 4 | MT-CO3 |
| 7.09E-25 | 0.659462 | 0.533 | 0.486 | 3.1955765[illegible] | 4 | COL6A3 |
| 4.28E-24 | 0.657457 | 0.538 | 0.533 | 1.93E-19 | 4 | STOX2 |
| 1.54E-36 | 0.656459 | 0.863 | 0.91 | 6.93E-32 | 4 | IGF2BP2 |
| 8.1850734[illegible] | 0.656421 | 0.787 | 0.859 | 3.69E-25 | 4 | FSTL1 |
| 3.23E-26 | 0.655244 | 0.518 | 0.498 | 1.45E-21 | 4 | TRIO |
| 1.6E-26 | 0.653531 | 0.617 | 0.616 | 7.23E-22 | 4 | ISLR |
| 4.08E-25 | 0.651191 | 0.805 | 0.787 | 1.8392219[illegible] | 4 | ALDH1A2 |
| 7.75E-29 | 0.650487 | 0.769 | 0.847 | 3.49E-24 | 4 | ZNF638 |
| 5.83E-35 | 0.644861 | 0.797 | 0.844 | 2.6257746[illegible] | 4 | PXDN |
| 7.55E-25 | 0.642957 | 0.599 | 0.646 | 3.4037920[illegible] | 4 | ASH1L |
| 7.77E-23 | 0.639507 | 0.769 | 0.786 | 3.5E-18 | 4 | COL1A2 |
| 1.23E-29 | 0.639162 | 0.508 | 0.437 | 5.55E-25 | 4 | INPPL1 |
| 3.81E-33 | 0.639148 | 0.926 | 0.976 | 1.72E-28 | 4 | LUC7L3 |
| 3.62E-28 | 0.636846 | 0.589 | 0.581 | 1.63E-23 | 4 | KMT2C |
| 2.68E-31 | 0.636634 | 0.668 | 0.68 | 1.21E-26 | 4 | TUG1 |
| 5.11E-36 | 0.635826 | 0.871 | 0.927 | 2.3E-31 | 4 | FGFR1 |
| 7.64E-24 | 0.632132 | 0.424 | 0.343 | 3.44E-19 | 4 | FAT3 |
| 6.46E-32 | 0.631359 | 0.652 | 0.649 | 2.91E-27 | 4 | SLC35E2B |
| 3.03E-27 | 0.630323 | 0.523 | 0.486 | 1.37E-22 | 4 | AHSA2 |
| 3.84E-24 | 0.628865 | 0.766 | 0.809 | 1.73E-19 | 4 | EFNB2 |
| 3.62E-19 | 0.628395 | 0.515 | 0.577 | 1.63E-14 | 4 | AGO3 |
| 3.76E-37 | 0.627598 | 0.794 | 0.828 | 1.7E-32 | 4 | PRRC2B |
| 1.7E-21 | 0.626181 | 0.508 | 0.506 | 7.66E-17 | 4 | EBF2 |
| 1.68E-22 | 0.626018 | 0.723 | 0.814 | 7.58E-18 | 4 | CCDC88A |
| 4.18E-26 | 0.625789 | 0.571 | 0.574 | 1.88E-21 | 4 | COL4A5 |
| 4.11E-53 | 0.624705 | 0.98 | 0.968 | 1.85E-48 | 4 | MEIS2 |
| 4.38E-22 | 0.624313 | 0.533 | 0.55 | 1.97E-17 | 4 | CASC15 |
| 1.53E-22 | 0.62416 | 0.65 | 0.713 | 6.91E-18 | 4 | HOXB3 |
| 1.9281369[illegible] | 0.62084 | 0.604 | 0.607 | 8.69E-16 | 4 | COL1A1 |
| 5.68E-24 | 0.618882 | 0.51 | 0.506 | 2.56E-19 | 4 | TPT1-AS1 |
| 1.81E-25 | 0.618813 | 0.655 | 0.701 | 8.17E-21 | 4 | MSI2 |
| 2.92E-24 | 0.615944 | 0.591 | 0.614 | 1.32E-19 | 4 | MEIS1 |
| 9.75E-21 | 0.614435 | 0.596 | 0.649 | 4.39E-16 | 4 | ROBO1 |
| 7.01E-23 | 0.614278 | 0.599 | 0.629 | 3.16E-18 | 4 | BOC |
| 5.27E-29 | 0.613665 | 0.739 | 0.813 | 2.38E-24 | 4 | PKN2 |
| 8.07E-28 | 0.613172 | 0.655 | 0.658 | 3.64E-23 | 4 | UNC5B |

| | | | | | | |
|---|---|---|---|---|---|---|
| 2.72E-22 | 0.612328 | 0.619 | 0.69 | 1.23E-17 | 4 | MDM4 |
| 2.21E-25 | 0.610919 | 0.541 | 0.555 | 9.96E-21 | 4 | OGT |
| 5.17E-28 | 0.610619 | 0.716 | 0.741 | 2.33E-23 | 4 | GPC6 |
| 6.23E-21 | 0.609229 | 0.437 | 0.401 | 2.81E-16 | 4 | PLXNA4 |
| 1.85E-25 | 0.607854 | 0.53 | 0.516 | 8.34E-21 | 4 | NFAT5 |
| 1.56E-31 | 0.606377 | 0.612 | 0.578 | 7.05E-27 | 4 | ADAMTS7 |
| 2.67E-46 | 0.60529 | 0.952 | 0.984 | 1.2E-41 | 4 | GTF2I |
| 1.6E-21 | 0.60362 | 0.637 | 0.718 | 7.21E-17 | 4 | POGZ |
| 8.95E-26 | 0.603595 | 0.576 | 0.604 | 4.04E-21 | 4 | EPHB3 |
| 1.38E-21 | 0.596454 | 0.579 | 0.629 | 6.23E-17 | 4 | FRMD4A |
| 1.24E-27 | 0.596051 | 0.68 | 0.735 | 5.59E-23 | 4 | NUMA1 |
| 1.29E-21 | 0.594534 | 0.563 | 0.602 | 5.81E-17 | 4 | RUNX1T1 |
| 5.59E-27 | 0.593558 | 0.543 | 0.523 | 2.52E-22 | 4 | LENG8 |
| 6.1720736[illegible] | 0.592867 | 1 | 0.997 | 2.78E-55 | 4 | MT-ND3 |
| 6.4261971[illegible] | 0.5918 | 1 | 0.999 | 2.9E-55 | 4 | MALAT1 |
| 9.78E-27 | 0.59054 | 0.536 | 0.532 | 4.41E-22 | 4 | C16orf72 |
| 1.7987415[illegible] | 0.589975 | 0.642 | 0.73 | 8.11E-16 | 4 | KIDINS220 |
| 6.92E-25 | 0.585496 | 0.561 | 0.571 | 3.1190759[illegible] | 4 | BIRC6 |
| 3.2755343[illegible] | 0.585118 | 0.528 | 0.592 | 1.48E-15 | 4 | TAOK1 |
| 6.29E-19 | 0.581293 | 0.406 | 0.407 | 2.83E-14 | 4 | ANKRD36C |
| 5.47E-24 | 0.580887 | 0.477 | 0.452 | 2.47E-19 | 4 | ATXN1 |
| 1.18E-13 | 0.579784 | 0.444 | 0.56 | 5.3E-09 | 4 | MT-ND6 |
| 6.48E-19 | 0.579427 | 0.373 | 0.336 | 2.92E-14 | 4 | CACNA1C |
| 2.41E-23 | 0.578899 | 0.848 | 0.944 | 1.09E-18 | 4 | N4BP2L2 |
| 1.19E-23 | 0.578705 | 0.52 | 0.531 | 5.36E-19 | 4 | KLF12 |
| 9.53E-23 | 0.578588 | 0.85 | 0.931 | 4.3E-18 | 4 | BPTF |
| 1.5887800[illegible] | 0.577599 | 0.594 | 0.663 | 7.16E-16 | 4 | TNS3 |
| 2.47E-46 | 0.576021 | 0.997 | 0.998 | 1.11E-41 | 4 | MT-CO2 |
| 9.2167151[illegible] | 0.575759 | 0.739 | 0.763 | 4.15E-15 | 4 | TGFBI |
| 6.58E-24 | 0.570397 | 0.614 | 0.674 | 2.96E-19 | 4 | LTBP4 |
| 2.96E-15 | 0.569403 | 0.726 | 0.822 | 1.3346261[illegible] | 4 | COL5A2 |
| 7.85E-18 | 0.565865 | 0.647 | 0.773 | 3.54E-13 | 4 | KMT2A |
| 5.1E-22 | 0.565747 | 0.673 | 0.772 | 2.3E-17 | 4 | ZC3H11A |
| 5.8539140[illegible] | 0.565283 | 0.655 | 0.766 | 2.64E-15 | 4 | DYNC1LI2 |
| 2.19E-19 | 0.562416 | 0.396 | 0.363 | 9.86E-15 | 4 | WNT5A |
| 1.37E-19 | 0.556542 | 0.779 | 0.899 | 6.18E-15 | 4 | JMJD1C |
| 3.88E-34 | 0.554652 | 0.926 | 0.971 | 1.75E-29 | 4 | HNRNPH1 |
| 1.58E-16 | 0.553844 | 0.602 | 0.716 | 7.13E-12 | 4 | NRIP1 |
| 2.93E-24 | 0.553723 | 0.541 | 0.534 | 1.32E-19 | 4 | ZNF827 |
| 8.18E-18 | 0.552131 | 0.596 | 0.705 | 3.69E-13 | 4 | PSMA3-AS1 |
| 1.39E-19 | 0.551734 | 0.723 | 0.837 | 6.25E-15 | 4 | ZKSCAN1 |
| 7.2510321[illegible] | 0.551188 | 0.475 | 0.592 | 3.27E-05 | 4 | KCTD12 |
| 2.38E-18 | 0.549753 | 0.784 | 0.884 | 1.07E-13 | 4 | TCF4 |
| 1.07E-19 | 0.549595 | 0.485 | 0.523 | 4.83E-15 | 4 | UBR5 |
| 2.64E-23 | 0.547783 | 0.528 | 0.557 | 1.19E-18 | 4 | SENP7 |
| 6.51E-22 | 0.547365 | 0.503 | 0.515 | 2.93E-17 | 4 | KIAA1109 |
| 2.82E-18 | 0.546925 | 0.579 | 0.673 | 1.27E-13 | 4 | SUGP2 |
| 2.17E-23 | 0.546631 | 0.353 | 0.231 | 9.77E-19 | 4 | GOLGA8A |
| 1.58E-21 | 0.544672 | 0.629 | 0.724 | 7.12E-17 | 4 | ANKRD10 |
| 4.02E-22 | 0.542847 | 0.464 | 0.466 | 1.81E-17 | 4 | FO538757.2 |

| | | | | | | |
|---|---|---|---|---|---|---|
| 5.62E-19 | 0.541945 | 0.388 | 0.365 | 2.53E-14 | 4 | FAT4 |
| 7.69E-15 | 0.539117 | 0.404 | 0.441 | 3.4657416 | 4 | EPHA3 |
| 7.2E-25 | 0.538665 | 0.711 | 0.753 | 3.2430832 | 4 | EMILIN1 |
| 5.4232567 | 0.534757 | 0.736 | 0.866 | 2.44E-15 | 4 | SSBP3 |
| 2.3E-19 | 0.534562 | 0.508 | 0.558 | 1.04E-14 | 4 | SMG1 |
| 2.21E-38 | 0.5331 | 0.997 | 0.997 | 9.96E-34 | 4 | MT-ND4 |
| 1.12E-16 | 0.532761 | 0.779 | 0.913 | 5.06E-12 | 4 | ZNF207 |
| 5.2642723 | 0.532739 | 0.787 | 0.906 | 2.37E-15 | 4 | RBFOX2 |
| 2.51E-14 | 0.532688 | 0.703 | 0.824 | 1.13E-09 | 4 | TCF7L2 |
| 7.71E-17 | 0.532298 | 0.495 | 0.564 | 3.48E-12 | 4 | REV3L |
| 2.14E-14 | 0.531293 | 0.411 | 0.46 | 9.6227111 | 4 | ADGRL3 |
| 2.56E-19 | 0.530858 | 0.492 | 0.54 | 1.15E-14 | 4 | FOXO3 |
| 1.68E-15 | 0.530724 | 0.591 | 0.726 | 7.55E-11 | 4 | SENP6 |
| 1.61E-16 | 0.529521 | 0.449 | 0.472 | 7.24E-12 | 4 | DCX |
| 3.38E-15 | 0.529355 | 0.409 | 0.445 | 1.5243281 | 4 | COL25A1 |
| 2.66E-14 | 0.528166 | 0.744 | 0.853 | 1.2E-09 | 4 | TRIL |
| 5.7695574 | 0.528121 | 0.447 | 0.45 | 2.6E-15 | 4 | ZCCHC24 |
| 4.5E-15 | 0.527308 | 0.553 | 0.685 | 2.0298744 | 4 | SETD2 |
| 4.81E-16 | 0.527028 | 0.586 | 0.711 | 2.17E-11 | 4 | RBM26 |
| 2.95E-15 | 0.526526 | 0.635 | 0.782 | 1.3303627 | 4 | SREK1 |
| 2.7408447 | 0.523914 | 0.497 | 0.553 | 1.24E-15 | 4 | B4GALNT4 |
| 1.48E-22 | 0.523566 | 0.482 | 0.456 | 6.68E-18 | 4 | COL5A1 |
| 1.34E-17 | 0.522508 | 0.495 | 0.565 | 6.02E-13 | 4 | MED13L |
| 1.58E-18 | 0.517376 | 0.434 | 0.438 | 7.12E-14 | 4 | PDE10A |
| 4.5197696 | 0.517354 | 0.5 | 0.527 | 2.04E-15 | 4 | GSE1 |
| 6.2330421 | 0.516464 | 0.447 | 0.451 | 2.81E-15 | 4 | GIGYF1 |
| 3.57E-16 | 0.515779 | 0.515 | 0.584 | 1.61E-11 | 4 | ZFHX4 |
| 1.8995229 | 0.51533 | 0.561 | 0.622 | 8.56E-16 | 4 | TTC17 |
| 1.2E-15 | 0.514877 | 0.569 | 0.668 | 5.4E-11 | 4 | TRPS1 |
| 7.12E-11 | 0.514588 | 0.632 | 0.805 | 3.21E-06 | 4 | GABPB1-AS1 |
| 8.8E-15 | 0.513278 | 0.607 | 0.753 | 3.9676344 | 4 | CD46 |
| 1.36E-17 | 0.511821 | 0.86 | 0.953 | 6.12E-13 | 4 | RBM25 |
| 2.2E-11 | 0.511085 | 0.35 | 0.407 | 9.91E-07 | 4 | SLIT2 |
| 3.52E-19 | 0.509103 | 0.855 | 0.941 | 1.59E-14 | 4 | TTC3 |
| 5.84E-23 | 0.508761 | 0.485 | 0.479 | 2.63E-18 | 4 | DLG5 |
| 1.21E-13 | 0.508228 | 0.548 | 0.7 | 5.45E-09 | 4 | PHF20L1 |
| 1.7293606 | 0.508196 | 0.614 | 0.783 | 7.79E-06 | 4 | POLR2J3 |
| 2.87E-21 | 0.507161 | 0.416 | 0.371 | 1.29E-16 | 4 | FLNC |
| 1.19E-16 | 0.506928 | 0.703 | 0.838 | 5.37E-12 | 4 | TBL1XR1 |
| 8.56E-18 | 0.506581 | 0.391 | 0.4 | 3.86E-13 | 4 | ANKRD36 |
| 4.24E-19 | 0.505687 | 0.754 | 0.862 | 1.91E-14 | 4 | EWSR1 |
| 1.29E-13 | 0.504455 | 0.454 | 0.577 | 5.83E-09 | 4 | ATM |
| 2.57E-17 | 0.503857 | 0.53 | 0.619 | 1.16E-12 | 4 | QSER1 |
| 1.93E-12 | 0.503619 | 0.652 | 0.796 | 8.68E-08 | 4 | PHIP |
| 5.9E-11 | 0.501848 | 0.353 | 0.419 | 2.66E-06 | 4 | ADAMTS1 |
| 1.47E-16 | 0.501033 | 0.503 | 0.567 | 6.61E-12 | 4 | KLHL24 |
| 4.56E-17 | 0.50096 | 0.571 | 0.666 | 2.05E-12 | 4 | RSRP1 |
| 1.48E-16 | 0.500892 | 0.444 | 0.52 | 6.68E-12 | 4 | PUM2 |
| 3.8E-16 | 0.500176 | 0.495 | 0.59 | 1.71E-11 | 4 | LRP6 |
| 6.22E-16 | 0.500155 | 0.566 | 0.68 | 2.8E-11 | 4 | TTC14 |

Supplementary Table 1

### DE genes for each cluster identified in tdTomato RET day 7 organoid data

Cluster 5

| p_val | avg_logFC | pct.1 | pct.2 | p_val_adj | cluster | gene |
|---|---|---|---|---|---|---|
| 2.8E-184 | 1.301554 | 0.992 | 0.788 | 1.3E-179 | 5 | IGFBP5 |
| 3.01E-29 | 0.899722 | 0.628 | 0.373 | 1.36E-24 | 5 | G0S2 |
| 8.23E-85 | 0.746527 | 0.984 | 0.88 | 3.7113404 | 5 | IGF2 |
| 2.05E-47 | 0.715854 | 0.709 | 0.302 | 9.24E-43 | 5 | L1TD1 |
| 6.89E-54 | 0.715374 | 0.884 | 0.677 | 3.11E-49 | 5 | SERPINE2 |
| 7.9E-21 | 0.699827 | 0.553 | 0.337 | 3.56E-16 | 5 | LUM |
| 9.73E-53 | 0.651552 | 0.951 | 0.829 | 4.38E-48 | 5 | COTL1 |
| 8.37E-69 | 0.624271 | 0.995 | 0.969 | 3.77E-64 | 5 | IGFBP2 |
| 5.89E-54 | 0.615534 | 0.873 | 0.629 | 2.66E-49 | 5 | TBX3 |
| 6.56E-15 | 0.582316 | 0.372 | 0.206 | 2.9576691 | 5 | DCN |
| 3.51E-79 | 0.571409 | 0.968 | 0.873 | 1.58E-74 | 5 | GYPC |
| 1.01E-31 | 0.553326 | 0.663 | 0.436 | 4.55E-27 | 5 | TCEAL7 |
| 3.5E-39 | 0.546193 | 0.951 | 0.898 | 1.58E-34 | 5 | TPM1 |
| 4.56E-62 | 0.542766 | 0.957 | 0.848 | 2.06E-57 | 5 | IFITM2 |
| 3.08E-32 | 0.532318 | 0.771 | 0.519 | 1.39E-27 | 5 | AKAP12 |
| 2.99E-49 | 0.516309 | 0.962 | 0.849 | 1.35E-44 | 5 | SCX |
| 8.55E-16 | 0.514039 | 0.59 | 0.448 | 3.85E-11 | 5 | LGALS1 |

Supplementary Table 1

DE genes for each cluster identified in tdTomato RET day 7 organoid data

Cluster 6

| p_val | avg_logFC | pct.1 | pct.2 | p_val_adj | cluster | gene |
|---|---|---|---|---|---|---|
| 3.6E-217 | 2.073546 | 1 | 0.7 | 1.6E-212 | 6 | EGFL7 |
| 8.2E-178 | 1.944123 | 0.997 | 0.792 | 3.7E-173 | 6 | RAMP2 |
| 1.1E-101 | 1.554878 | 0.912 | 0.036 | 4.89E-97 | 6 | KDR |
| 5.9E-107 | 1.522941 | 0.921 | 0.049 | 2.7E-102 | 6 | ICAM2 |
| 4.3E-133 | 1.503341 | 0.991 | 0.647 | 1.9E-128 | 6 | IGFBP4 |
| 8.7E-109 | 1.484087 | 0.951 | 0.28 | 3.9E-104 | 6 | PLVAP |
| 9.1E-114 | 1.422265 | 0.924 | 0.027 | 4.1E-109 | 6 | ESAM |
| 4.4E-103 | 1.360373 | 0.976 | 0.787 | 1.99E-98 | 6 | GNG11 |
| 4.9E-106 | 1.358388 | 0.951 | 0.544 | 2.2E-101 | 6 | VAMP5 |
| 2E-121 | 1.350679 | 0.985 | 0.67 | 8.9E-117 | 6 | PRCP |
| 2.8E-139 | 1.301851 | 0.988 | 0.758 | 1.3E-134 | 6 | TFPI |
| 1.49E-26 | 1.265379 | 0.383 | 0.007 | 6.69E-22 | 6 | CLDN5 |
| 1.27E-94 | 1.241576 | 0.857 | 0.017 | 5.7233818[illegible] | 6 | ECSCR.1 |
| 6.07E-72 | 1.204033 | 0.793 | 0.177 | 2.74E-67 | 6 | S100A16 |
| 2.9E-127 | 1.193399 | 0.985 | 0.739 | 1.3E-122 | 6 | CCDC85B |
| 6.6743980[illegible] | 1.143574 | 0.878 | 0.288 | 3.01E-75 | 6 | RGS5 |
| 2.75E-85 | 1.129303 | 0.863 | 0.077 | 1.2413702[illegible] | 6 | SOX18 |
| 2.27E-89 | 1.075936 | 0.869 | 0.101 | 1.02E-84 | 6 | PROCR |
| 2.03E-56 | 1.075316 | 0.881 | 0.624 | 9.13E-52 | 6 | S100A6 |
| 1.9872730[illegible] | 1.03518 | 0.854 | 0.559 | 8.96E-56 | 6 | NR2F2 |
| 1.2E-114 | 1.033373 | 0.973 | 0.586 | 5.1961337[illegible] | 6 | LMO2 |
| 5.4290821[illegible] | 1.004579 | 0.845 | 0.1 | 2.45E-75 | 6 | MMRN1 |
| 2.13E-49 | 1.00252 | 0.653 | 0.031 | 9.61E-45 | 6 | KLF2 |
| 3.77E-43 | 0.989362 | 0.568 | 0.007 | 1.7E-38 | 6 | APLNR |
| 1.0279749[illegible] | 0.981157 | 0.824 | 0.035 | 4.63E-76 | 6 | PECAM1 |
| 5.1E-109 | 0.973275 | 0.994 | 0.921 | 2.3E-104 | 6 | CALM1 |
| 3.1921870[illegible] | 0.970512 | 0.757 | 0.021 | 1.44E-65 | 6 | CDH5 |
| 7.62E-45 | 0.959366 | 0.842 | 0.56 | 3.4354855[illegible] | 6 | ANXA2 |
| 3.67E-53 | 0.939647 | 0.69 | 0.059 | 1.66E-48 | 6 | CLEC14A |
| 3.0333842[illegible] | 0.936137 | 0.769 | 0.067 | 1.37E-65 | 6 | TIE1 |
| 3.06E-35 | 0.925778 | 0.495 | 0.022 | 1.3812987[illegible] | 6 | GJA4 |
| 9.92E-59 | 0.920266 | 0.821 | 0.37 | 4.47E-54 | 6 | S100A10 |
| 2.77E-92 | 0.911331 | 0.891 | 0.096 | 1.25E-87 | 6 | HHEX |
| 1.76E-61 | 0.905759 | 0.787 | 0.158 | 7.95E-57 | 6 | CALCRL |
| 2.57E-59 | 0.897598 | 0.951 | 0.815 | 1.16E-54 | 6 | HYAL2 |
| 7.61E-46 | 0.889648 | 0.86 | 0.654 | 3.43E-41 | 6 | NRP2 |
| 2.86E-77 | 0.884442 | 0.921 | 0.581 | 1.29E-72 | 6 | CTSC |
| 3.34E-88 | 0.858321 | 0.982 | 0.889 | 1.5E-83 | 6 | FKBP1A |
| 4.7E-102 | 0.852004 | 0.976 | 0.662 | 2.1E-97 | 6 | MEF2C |
| 1.84E-92 | 0.827836 | 0.915 | 0.225 | 8.28E-88 | 6 | LYL1 |
| 5.52E-81 | 0.827279 | 0.839 | 0.077 | 2.49E-76 | 6 | HAPLN3 |
| 3.3E-85 | 0.81496 | 0.96 | 0.653 | 1.4855704[illegible] | 6 | FAM107B |
| 7.16E-79 | 0.803841 | 0.948 | 0.616 | 3.23E-74 | 6 | FCGRT |
| 1.35E-46 | 0.803418 | 0.818 | 0.411 | 6.08E-42 | 6 | DAB2 |
| 2.26E-54 | 0.798595 | 0.669 | 0.056 | 1.02E-49 | 6 | KANK3 |
| 4.59E-67 | 0.795727 | 0.748 | 0.023 | 2.07E-62 | 6 | TEK |

| | | | | | | |
|---|---|---|---|---|---|---|
| 1.62E-78 | 0.792579 | 0.809 | 0.048 | 7.32E-74 | 6 | SOX7 |
| 3.7E-95 | 0.791303 | 0.991 | 0.908 | 1.6696411 | 6 | HSPB1 |
| 3.2133232 | 0.789703 | 0.821 | 0.351 | 1.45E-55 | 6 | HSPG2 |
| 9.71E-57 | 0.788389 | 0.775 | 0.202 | 4.38E-52 | 6 | PLXND1 |
| 2.22E-61 | 0.776599 | 0.833 | 0.263 | 1E-56 | 6 | HAPLN1 |
| 1.05E-46 | 0.767878 | 0.672 | 0.172 | 4.71E-42 | 6 | GBP4 |
| 3.31E-59 | 0.765916 | 0.796 | 0.291 | 1.49E-54 | 6 | TMEM88 |
| 1.47E-64 | 0.764137 | 0.751 | 0.058 | 6.6042134 | 6 | ENTPD1 |
| 1.47E-63 | 0.76208 | 0.711 | 0.027 | 6.64E-59 | 6 | MMRN2 |
| 3.29E-77 | 0.761015 | 0.951 | 0.692 | 1.48E-72 | 6 | ELK3 |
| 1.04E-54 | 0.746284 | 0.79 | 0.291 | 4.6940294 | 6 | ITM2A |
| 2.01E-59 | 0.736656 | 0.787 | 0.203 | 9.04E-55 | 6 | ADGRL4 |
| 1.92E-69 | 0.73318 | 0.772 | 0.053 | 8.68E-65 | 6 | RASIP1 |
| 5.5619804 | 0.728386 | 0.854 | 0.414 | 2.51E-55 | 6 | ARHGAP29 |
| 1.34E-64 | 0.711724 | 0.821 | 0.269 | 6.0344816 | 6 | TGFB1 |
| 1.5E-44 | 0.705277 | 0.739 | 0.314 | 6.7767835 | 6 | ARL4A |
| 4.81E-66 | 0.704713 | 0.851 | 0.279 | 2.17E-61 | 6 | MTUS1 |
| 1.28E-52 | 0.692035 | 0.644 | 0.005 | 5.77E-48 | 6 | BCL6B |
| 1.99E-71 | 0.676468 | 0.763 | 0.028 | 8.95E-67 | 6 | ERG |
| 2.22E-52 | 0.675968 | 0.821 | 0.415 | 1E-47 | 6 | EMCN |
| 2.9732022 | 0.675742 | 0.812 | 0.383 | 1.34E-45 | 6 | HLA-E |
| 5.0644414 | 0.666056 | 1 | 0.979 | 2.3E-155 | 6 | VIM |
| 6.08E-75 | 0.665846 | 0.954 | 0.699 | 2.7400757 | 6 | LIMCH1 |
| 4.46E-65 | 0.663644 | 0.973 | 0.892 | 2.0102360 | 6 | RHOC |
| 1.07E-83 | 0.652848 | 0.994 | 0.965 | 4.84E-79 | 6 | FSCN1 |
| 3.04E-66 | 0.639365 | 0.763 | 0.106 | 1.37E-61 | 6 | SHE |
| 9.85E-54 | 0.637316 | 0.763 | 0.288 | 4.44E-49 | 6 | LDB2 |
| 9.49E-58 | 0.635171 | 0.714 | 0.071 | 4.28E-53 | 6 | FLT1 |
| 1.83E-51 | 0.634785 | 0.644 | 0.011 | 8.25E-47 | 6 | GIMAP1 |
| 1.28E-66 | 0.632945 | 0.954 | 0.809 | 5.76E-62 | 6 | RDX |
| 1.29E-49 | 0.628253 | 0.839 | 0.464 | 5.82E-45 | 6 | COL4A1 |
| 1.28E-48 | 0.61293 | 0.979 | 0.86 | 5.76E-44 | 6 | BST2 |
| 4.81E-47 | 0.609625 | 0.775 | 0.41 | 2.17E-42 | 6 | FAM89A |
| 3.73E-57 | 0.608403 | 0.851 | 0.493 | 1.68E-52 | 6 | ARPC1B |
| 3.75E-44 | 0.606696 | 0.59 | 0.007 | 1.69E-39 | 6 | GIMAP4 |
| 6.6374352 | 0.602312 | 0.547 | 0.027 | 2.99E-35 | 6 | RNASE1 |
| 8.34E-47 | 0.591755 | 0.672 | 0.148 | 3.76E-42 | 6 | SDPR |
| 7.1585861 | 0.591028 | 0.954 | 0.825 | 3.23E-35 | 6 | FN1 |
| 9.16E-28 | 0.587273 | 0.456 | 0.063 | 4.13E-23 | 6 | TM4SF1 |
| 5.04E-45 | 0.57681 | 0.863 | 0.64 | 2.2714429 | 6 | FAM69B |
| 3.46E-29 | 0.558945 | 0.432 | 0.011 | 1.56E-24 | 6 | CD34 |
| 2.37E-25 | 0.552555 | 0.383 | 0.002 | 1.0669244 | 6 | CD93 |
| 7.53E-46 | 0.547969 | 0.635 | 0.077 | 3.4E-41 | 6 | NPR3 |
| 3.5941587 | 0.547605 | 0.933 | 0.852 | 1.62E-25 | 6 | RPS27L |
| 3.63E-41 | 0.545308 | 0.547 | 0.006 | 1.63E-36 | 6 | STAB1 |
| 3.48E-48 | 0.544027 | 0.714 | 0.262 | 1.57E-43 | 6 | TACC1 |
| 8.03E-53 | 0.537649 | 0.866 | 0.538 | 3.62E-48 | 6 | MSN |
| 1.55E-42 | 0.536367 | 0.611 | 0.024 | 6.98E-38 | 6 | GDPD5 |
| 6.5828782 | 0.534656 | 0.641 | 0.028 | 2.97E-45 | 6 | RASGRP3 |
| 1.41E-44 | 0.531699 | 0.881 | 0.702 | 6.3325970 | 6 | APOA1BP |

| | | | | | | |
|---|---|---|---|---|---|---|
| 1.5E-56 | 0.530713 | 0.976 | 0.872 | 6.75E-52 | 6 | SPTBN1 |
| 2.35E-47 | 0.524592 | 0.973 | 0.924 | 1.06E-42 | 6 | GNAI2 |
| 7.01153378 | 0.514112 | 0.997 | 0.994 | 3.16E-35 | 6 | TMSB4X |
| 1.46E-44 | 0.511354 | 0.76 | 0.35 | 6.5709755 | 6 | TSPAN18 |
| 3.55E-37 | 0.508745 | 0.495 | 0.004 | 1.6E-32 | 6 | NT5E |
| 1.46E-36 | 0.507947 | 0.748 | 0.422 | 6.56E-32 | 6 | ETS2 |
| 6.87E-37 | 0.507425 | 0.611 | 0.194 | 3.1E-32 | 6 | ADAM15 |
| 6.65E-34 | 0.50461 | 0.465 | 0.004 | 3E-29 | 6 | PTRF |

Supplementary Table 1

### DE genes for each cluster identified in tdTomato RET day 7 organoid data

Cluster 7

| p_val | avg_logFC | pct.1 | pct.2 | p_val_adj | cluster | gene |
|---|---|---|---|---|---|---|
| 4.13E-16 | 2.248299 | 0.944 | 0.015 | 1.86E-11 | 7 | PLSCR2 |
| 1.85E-16 | 1.857828 | 0.972 | 0.056 | 8.34E-12 | 7 | SERHL2 |
| 1.1378539 | 1.613563 | 0.917 | 0.395 | 5.13E-06 | 7 | S100A10 |
| 1E-11 | 1.5082 | 0.889 | 0.007 | 4.52E-07 | 7 | ADAMTS8 |
| 6.74E-05 | 1.39763 | 0.5 | 0.005 | 1 | 7 | GYPB |
| 5.74E-11 | 1.232811 | 0.861 | 0.048 | 2.59E-06 | 7 | RASGRP1 |
| 1.54E-09 | 1.167735 | 0.944 | 0.166 | 6.93E-05 | 7 | SERINC5 |
| 0.004754 | 1.146203 | 0.306 | 0.001 | 1 | 7 | LINC01467 |
| 8.52E-09 | 1.105988 | 0.778 | 0.011 | 0.000384 | 7 | CCDC81 |
| 4.62E-08 | 1.088309 | 0.944 | 0.679 | 0.002084 | 7 | APOE |
| 2.69E-13 | 1.088014 | 1 | 0.559 | 1.21E-08 | 7 | CFLAR |
| 1.1754739 | 1.049706 | 0.917 | 0.2 | 5.3E-06 | 7 | ALX1 |
| 2.2726928 | 1.041158 | 0.861 | 0.007 | 1.02E-05 | 7 | RP11-119D9.1 |
| 5.04E-09 | 1.03531 | 0.861 | 0.191 | 0.000227 | 7 | ERBB4 |
| 6.62E-06 | 0.995816 | 1 | 0.959 | 0.298364 | 7 | PRDX1 |
| 5.93E-07 | 0.989158 | 0.861 | 0.408 | 0.026742 | 7 | IER3 |
| 5.99E-08 | 0.980299 | 0.833 | 0.175 | 0.0027 | 7 | DGKI |
| 4.97E-12 | 0.97957 | 1 | 0.913 | 2.24E-07 | 7 | RHOBTB3 |
| 6.46E-07 | 0.957653 | 0.972 | 0.927 | 0.029103 | 7 | PKM |
| 0.003641 | 0.903459 | 0.556 | 0.107 | 1 | 7 | NPY |
| 4.79E-18 | 0.899613 | 1 | 0.954 | 2.16E-13 | 7 | LAPTM4B |
| 1.29E-08 | 0.886367 | 0.972 | 0.772 | 0.00058 | 7 | TFPI |
| 1.19E-09 | 0.862104 | 0.861 | 0.214 | 5.35E-05 | 7 | CNTNAP2 |
| 7.4344548 | 0.859573 | 0.972 | 0.571 | 3.35E-05 | 7 | NFKBIA |
| 3.58E-11 | 0.852764 | 0.944 | 0.22 | 1.61E-06 | 7 | PCDH10 |
| 2.52E-06 | 0.835681 | 0.667 | 0.034 | 0.113561 | 7 | TECRL |
| 1.39E-05 | 0.821323 | 0.528 | 0.003 | 0.628537 | 7 | PI15 |
| 2.4651289 | 0.807262 | 0.917 | 0.3 | 1.11E-05 | 7 | TGFB1 |
| 9.09E-07 | 0.785994 | 0.833 | 0.487 | 0.040983 | 7 | NRXN1 |
| 1.56E-06 | 0.78171 | 0.833 | 0.454 | 0.070119 | 7 | CKB |
| 1.01E-12 | 0.776572 | 1 | 0.552 | 4.55E-08 | 7 | LHX1 |
| 1.41E-09 | 0.758509 | 1 | 0.813 | 6.36E-05 | 7 | COL5A2 |
| 2.56E-06 | 0.740474 | 0.889 | 0.642 | 0.115321 | 7 | HTRA1 |
| 0.001963 | 0.726445 | 0.333 | 0.001 | 1 | 7 | CXCL13 |
| 9.9644346 | 0.723274 | 1 | 0.984 | 4.49E-05 | 7 | NGFRAP1 |
| 6.69E-09 | 0.710874 | 0.972 | 0.783 | 0.000302 | 7 | COL1A2 |
| 2.2E-12 | 0.690512 | 0.972 | 0.498 | 9.93E-08 | 7 | C1GALT1 |
| 1.46E-06 | 0.688813 | 0.833 | 0.333 | 0.065784 | 7 | MAP2 |
| 6.64E-06 | 0.67639 | 0.667 | 0.054 | 0.299473 | 7 | GRID2 |
| 1.9E-05 | 0.666545 | 0.583 | 0.001 | 0.855507 | 7 | TBR1 |
| 4.43E-07 | 0.660237 | 0.972 | 0.697 | 0.019965 | 7 | FBN1 |
| 6.64E-07 | 0.654844 | 0.694 | 0.08 | 0.029919 | 7 | AMIGO2 |
| 1.37E-05 | 0.650679 | 0.611 | 0.06 | 0.619491 | 7 | OLFM3 |
| 2.03E-08 | 0.634804 | 0.806 | 0.141 | 0.000916 | 7 | KCNK1 |
| 3.8E-05 | 0.618042 | 0.556 | 0.055 | 1 | 7 | C6orf141 |
| 0.000141 | 0.605703 | 0.583 | 0.231 | 1 | 7 | SULF1 |

| | | | | | | |
|---|---|---|---|---|---|---|
| 1.22E-08 | 0.595051 | 0.972 | 0.745 | 0.000549 | 7 | COL2A1 |
| 1.77E-07 | 0.594868 | 0.722 | 0.116 | 0.007991 | 7 | GABRB3 |
| 1.44E-09 | 0.594139 | 1 | 0.953 | 6.47E-05 | 7 | CRABP2 |
| 6.24E-12 | 0.593402 | 1 | 0.796 | 2.81E-07 | 7 | PHLDA1 |
| 0.006336 | 0.59184 | 0.889 | 0.751 | 1 | 7 | UBE2S |
| 1.87E-06 | 0.588951 | 0.972 | 0.768 | 0.084265 | 7 | CPE |
| 0.000373 | 0.577585 | 0.917 | 0.782 | 1 | 7 | BEX1 |
| 6.93E-07 | 0.557983 | 0.611 | 0.026 | 0.031218 | 7 | MARCH11 |
| 4.41E-05 | 0.556934 | 0.667 | 0.185 | 1 | 7 | TMEM47 |
| 1.09E-07 | 0.55465 | 0.806 | 0.175 | 0.00492 | 7 | ACOX3 |
| 2.73E-05 | 0.549362 | 0.583 | 0.001 | 1 | 7 | EOMES |
| 2.3E-05 | 0.538559 | 0.556 | 0.04 | 1 | 7 | FGF12 |
| 5.66E-08 | 0.534858 | 0.806 | 0.063 | 0.002549 | 7 | MCF2L2 |
| 8.93E-05 | 0.528632 | 0.5 | 0.034 | 1 | 7 | CCKBR |
| 0.000155 | 0.520286 | 0.472 | 0.006 | 1 | 7 | DKK4 |
| 1.14E-06 | 0.520161 | 0.75 | 0.217 | 0.051171 | 7 | SIPA1L2 |
| 2.49E-05 | 0.518023 | 0.944 | 0.806 | 1 | 7 | MAP1B |
| 5.68E-08 | 0.502601 | 0.722 | 0.02 | 0.002558 | 7 | NTN4 |

Supplementary Table 1

DE genes for each cluster identified in tdTomato RET day 7 organoid data

Cluster 8

| p_val | avg_logFC | pct.1 | pct.2 | p_val_adj | cluster | gene |
|---|---|---|---|---|---|---|
| 0.001318 | 1.375986 | 0.583 | 0.366 | 1 | 8 | CRYM |
| 1.66E-06 | 1.134905 | 0.917 | 0.678 | 0.074893 | 8 | CENPW |
| 1.97E-08 | 1.02355 | 0.958 | 0.841 | 0.00089 | 8 | FABP5 |
| 1.95E-05 | 0.997194 | 0.875 | 0.603 | 0.878792 | 8 | KIAA0101 |
| 2.02E-06 | 0.974187 | 0.875 | 0.671 | 0.090982 | 8 | NQO2 |
| 4.94E-07 | 0.951495 | 0.958 | 0.927 | 0.022261 | 8 | TUBB2B |
| 1.6E-07 | 0.935776 | 0.958 | 0.632 | 0.007198 | 8 | TYMS |
| 7.53E-05 | 0.911031 | 0.708 | 0.412 | 1 | 8 | VAMP8 |
| 3.35E-05 | 0.901805 | 0.75 | 0.337 | 1 | 8 | KRT19 |
| 3.57E-05 | 0.852066 | 0.875 | 0.68 | 1 | 8 | KRT18 |
| 5.37E-08 | 0.849618 | 0.958 | 0.944 | 0.002421 | 8 | RANBP1 |
| 6.92E-08 | 0.842768 | 1 | 0.928 | 0.003119 | 8 | ODC1 |
| 0.004708 | 0.823733 | 0.5 | 0.249 | 1 | 8 | HIST1H1A |
| 9.62E-06 | 0.803338 | 0.958 | 0.845 | 0.433776 | 8 | LDHA |
| 0.004477 | 0.778655 | 0.375 | 0.134 | 1 | 8 | FEV |
| 7.53E-05 | 0.766944 | 0.875 | 0.746 | 1 | 8 | UCHL1 |
| 0.000143 | 0.764352 | 0.875 | 0.834 | 1 | 8 | METRN |
| 2.67E-11 | 0.764327 | 1 | 0.982 | 1.2E-06 | 8 | HMGN2 |
| 4.63E-05 | 0.757342 | 0.958 | 0.961 | 1 | 8 | ENO1 |
| 6.28E-11 | 0.754118 | 1 | 0.935 | 2.83E-06 | 8 | ALDOA |
| 4.52E-06 | 0.75064 | 0.917 | 0.794 | 0.20372 | 8 | ANXA5 |
| 0.007143 | 0.733848 | 0.917 | 0.94 | 1 | 8 | HIST1H4C |
| 9.1E-05 | 0.71872 | 0.833 | 0.742 | 1 | 8 | RPA3 |
| 0.001518 | 0.710511 | 0.875 | 0.893 | 1 | 8 | DUT |
| 4.82E-05 | 0.709676 | 0.708 | 0.324 | 1 | 8 | UCP2 |
| 0.003739 | 0.704348 | 0.5 | 0.285 | 1 | 8 | C12orf75 |
| 0.00252 | 0.701889 | 0.917 | 0.923 | 1 | 8 | DBI |
| 0.008955 | 0.700213 | 0.542 | 0.325 | 1 | 8 | RGS5 |
| 3.29E-12 | 0.698011 | 1 | 0.97 | 1.48E-07 | 8 | H2AFZ |
| 3.92E-08 | 0.687259 | 1 | 0.944 | 0.001766 | 8 | SNRPG |
| 3.37E-06 | 0.685526 | 0.958 | 0.873 | 0.151922 | 8 | NT5DC2 |
| 4.03125250 | 0.682405 | 1 | 0.982 | 1.82E-05 | 8 | GSTP1 |
| 3.81E-07 | 0.678138 | 1 | 0.945 | 0.017156 | 8 | HMGA1 |
| 1.94E-05 | 0.677818 | 0.958 | 0.91 | 0.876212 | 8 | LSM3 |
| 4.06E-08 | 0.675042 | 1 | 0.956 | 0.001829 | 8 | PPP1R14B |
| 3.07E-09 | 0.674253 | 1 | 0.94 | 0.000138 | 8 | SNRPB |
| 0.001057 | 0.674113 | 0.708 | 0.563 | 1 | 8 | IRX3 |
| 0.000627 | 0.664859 | 0.917 | 0.899 | 1 | 8 | MZT2A |
| 1.21E-13 | 0.664522 | 1 | 0.998 | 5.47E-09 | 8 | PTMA |
| 0.000392 | 0.661081 | 0.917 | 0.797 | 1 | 8 | CKS1B |
| 0.000178 | 0.659808 | 0.875 | 0.794 | 1 | 8 | AKR1B1 |
| 0.001329 | 0.656572 | 0.625 | 0.479 | 1 | 8 | FEN1 |
| 2.28E-08 | 0.656145 | 1 | 0.95 | 0.001026 | 8 | HSPE1 |
| 5.19E-07 | 0.650825 | 1 | 0.927 | 0.02338 | 8 | PKM |
| 3.75E-06 | 0.647196 | 1 | 0.978 | 0.168814 | 8 | NCL |
| 0.000146 | 0.644099 | 0.958 | 0.949 | 1 | 8 | TXN |

| | | | | | | |
|---|---|---|---|---|---|---|
| 3.9960946 | 0.636847 | 1 | 0.996 | 1.8E-05 | 8 | GAPDH |
| 1.92E-09 | 0.634802 | 1 | 0.993 | 8.63E-05 | 8 | RPL27A |
| 7.2E-05 | 0.631214 | 0.958 | 0.948 | 1 | 8 | MIF |
| 0.002784 | 0.62868 | 0.792 | 0.752 | 1 | 8 | YBX3 |
| 0.000902 | 0.628476 | 0.708 | 0.501 | 1 | 8 | SVIP |
| 0.000188 | 0.628322 | 0.958 | 0.933 | 1 | 8 | SEPW1 |
| 0.002579 | 0.627893 | 0.75 | 0.75 | 1 | 8 | CCDC167 |
| 4.54E-16 | 0.627517 | 1 | 0.98 | 2.04E-11 | 8 | HINT1 |
| 2.78E-05 | 0.625045 | 0.958 | 0.899 | 1 | 8 | MRPL11 |
| 0.000369 | 0.623391 | 0.833 | 0.794 | 1 | 8 | POLD2 |
| 0.001082 | 0.622287 | 0.917 | 0.911 | 1 | 8 | C1QBP |
| 2.64E-05 | 0.619441 | 1 | 0.929 | 1 | 8 | GADD45GIP1 |
| 1.79E-12 | 0.61867 | 1 | 0.977 | 8.07E-08 | 8 | MZT2B |
| 3.2E-07 | 0.618022 | 1 | 0.915 | 0.014444 | 8 | NME1 |
| 0.002409 | 0.618001 | 0.917 | 0.898 | 1 | 8 | PA2G4 |
| 1.28E-06 | 0.616329 | 1 | 0.955 | 0.057477 | 8 | SNRPD1 |
| 2.26E-07 | 0.612343 | 1 | 0.975 | 0.010193 | 8 | ATP5I |
| 1.11E-05 | 0.610571 | 0.875 | 0.578 | 0.498421 | 8 | DMKN |
| 2.02E-08 | 0.604025 | 1 | 0.983 | 0.00091 | 8 | TUBA1B |
| 2.97E-07 | 0.603945 | 1 | 0.966 | 0.013396 | 8 | SRSF9 |
| 0.005484 | 0.603872 | 0.833 | 0.883 | 1 | 8 | ATP5G1 |
| 0.000131 | 0.603411 | 0.958 | 0.897 | 1 | 8 | TPM3 |
| 0.000156 | 0.603207 | 1 | 0.931 | 1 | 8 | PRELID1 |
| 2.12E-12 | 0.601939 | 1 | 0.998 | 9.55E-08 | 8 | RPS2 |
| 0.008876 | 0.595917 | 0.583 | 0.522 | 1 | 8 | DHFR |
| 0.000912 | 0.595398 | 0.958 | 0.964 | 1 | 8 | YWHAQ |
| 0.000891 | 0.595349 | 0.75 | 0.565 | 1 | 8 | PDLIM1 |
| 5.29E-13 | 0.589602 | 1 | 0.997 | 2.38E-08 | 8 | RPL35 |
| 0.000686 | 0.587899 | 0.958 | 0.935 | 1 | 8 | LSM4 |
| 5.73E-09 | 0.587619 | 1 | 0.983 | 0.000258 | 8 | EEF1B2 |
| 1.86E-11 | 0.587239 | 1 | 0.997 | 8.4E-07 | 8 | RPS6 |
| 1.39E-07 | 0.584883 | 1 | 0.997 | 0.006257 | 8 | HMGB1 |
| 0.005591 | 0.583194 | 0.833 | 0.825 | 1 | 8 | SNRPA1 |
| 0.001816 | 0.582485 | 0.75 | 0.722 | 1 | 8 | ALYREF |
| 1.1E-09 | 0.582179 | 1 | 0.975 | 4.96E-05 | 8 | PFN1 |
| 7.3905032 | 0.580531 | 1 | 0.984 | 3.33E-05 | 8 | COX7C |
| 0.006467 | 0.573153 | 0.792 | 0.772 | 1 | 8 | PCBD1 |
| 0.001963 | 0.57311 | 0.917 | 0.918 | 1 | 8 | ROMO1 |
| 0.005336 | 0.570834 | 0.708 | 0.678 | 1 | 8 | CYB5A |
| 3.56E-07 | 0.570058 | 1 | 0.956 | 0.016054 | 8 | SLC25A5 |
| 1.31E-05 | 0.569082 | 1 | 0.952 | 0.589254 | 8 | TPI1 |
| 0.000116 | 0.568497 | 0.833 | 0.662 | 1 | 8 | RNASEH2A |
| 1.36E-07 | 0.568364 | 1 | 0.975 | 0.006144 | 8 | RAN |
| 0.001253 | 0.567655 | 0.917 | 0.921 | 1 | 8 | CCT5 |
| 0.009794 | 0.566399 | 0.833 | 0.9 | 1 | 8 | POLR2I |
| 2.42E-07 | 0.566217 | 1 | 0.96 | 0.010899 | 8 | PPDPF |
| 3.49E-05 | 0.564437 | 1 | 0.98 | 1 | 8 | HSP90AA1 |
| 0.004537 | 0.56439 | 0.667 | 0.559 | 1 | 8 | GINS2 |
| 7.03E-05 | 0.563632 | 0.917 | 0.83 | 1 | 8 | C16orf13 |
| 8.0043454 | 0.552941 | 1 | 0.974 | 3.61E-05 | 8 | UQCRH |

| | | | | | | |
|---|---|---|---|---|---|---|
| 3.32E-06 | 0.551967 | 1 | 0.958 | 0.149468 | 8 | NDUFS6 |
| 0.004626 | 0.550827 | 0.875 | 0.891 | 1 | 8 | NR2F6 |
| 0.008225 | 0.549283 | 0.667 | 0.654 | 1 | 8 | SLBP |
| 1.24E-08 | 0.544989 | 1 | 0.97 | 0.000557 | 8 | XRCC5 |
| 8.95E-06 | 0.540167 | 1 | 0.984 | 0.403287 | 8 | LDHB |
| 0.001595 | 0.540113 | 0.875 | 0.839 | 1 | 8 | IDI1 |
| 4.36E-05 | 0.538226 | 1 | 0.982 | 1 | 8 | PRDX2 |
| 0.000225 | 0.537941 | 0.958 | 0.887 | 1 | 8 | HNRNPAB |
| 0.009695 | 0.537599 | 0.625 | 0.512 | 1 | 8 | TSTD1 |
| 1.36E-07 | 0.536266 | 1 | 0.955 | 0.006151 | 8 | USMG5 |
| 3.29E-05 | 0.534745 | 1 | 0.955 | 1 | 8 | POLR2L |
| 0.000227 | 0.533692 | 0.792 | 0.62 | 1 | 8 | CYYR1 |
| 4.23E-05 | 0.532646 | 0.792 | 0.557 | 1 | 8 | MGAT4B |
| 0.000785 | 0.532211 | 0.792 | 0.723 | 1 | 8 | STIP1 |
| 3.46E-08 | 0.530773 | 1 | 0.9 | 0.001558 | 8 | FKBP3 |
| 1.53E-05 | 0.528783 | 1 | 0.918 | 0.689609 | 8 | CDK4 |
| 0.001433 | 0.52811 | 0.833 | 0.788 | 1 | 8 | CARHSP1 |
| 0.00736 | 0.524051 | 0.792 | 0.803 | 1 | 8 | CHCHD3 |
| 0.001293 | 0.523961 | 0.875 | 0.866 | 1 | 8 | CHCHD1 |
| 0.003554 | 0.521373 | 0.917 | 0.936 | 1 | 8 | CCT2 |
| 0.001667 | 0.518593 | 0.875 | 0.863 | 1 | 8 | PSMC3 |
| 0.003924 | 0.512789 | 0.833 | 0.801 | 1 | 8 | COMMD4 |
| 4.62E-05 | 0.508442 | 0.958 | 0.909 | 1 | 8 | GTF3C6 |
| 1.26E-05 | 0.508415 | 1 | 0.954 | 0.568907 | 8 | TRMT112 |
| 0.002487 | 0.507462 | 0.792 | 0.753 | 1 | 8 | SLC25A39 |
| 0.001738 | 0.505075 | 0.958 | 0.959 | 1 | 8 | ERH |
| 3.79E-06 | 0.500513 | 1 | 0.959 | 0.170919 | 8 | PRDX1 |
