## Supplementary Table 2 for "*In vitro* plasticity between ureteric epithelial and distal nephron identity and maturity is controlled by extracellular signals"

DE gene lists for each subcluster within epithelial population of tdTomato RET day 7 organoid data

Cluster 0

| p_val | avg_logFC | pct.1 | pct.2 | p_val_adj | cluster | gene |
|---|---|---|---|---|---|---|
| 2.92E-94 | 0.845527 | 0.991 | 0.703 | 1.32E-89 | 0 | NEFM |
| 4.1E-132 | 0.812635 | 0.998 | 0.869 | 1.8E-127 | 0 | PTN |
| 1E-49 | 0.736838 | 0.946 | 0.685 | 4.51E-45 | 0 | IGFBP5 |
| 7.69E-93 | 0.690957 | 1 | 0.933 | 3.47E-88 | 0 | VIM |
| 2.07117210 | 0.555615 | 0.981 | 0.645 | 9.33E-86 | 0 | PDGFRA |
| 1.93E-75 | 0.544913 | 0.932 | 0.448 | 8.71E-71 | 0 | POSTN |
| 2.77E-75 | 0.543619 | 0.939 | 0.531 | 1.24679158 | 0 | ALDH1A2 |
| 3.99300368 | 0.514831 | 0.972 | 0.79 | 1.8E-45 | 0 | IGF2 |
| 3.4E-53 | 0.512929 | 0.979 | 0.785 | 1.53E-48 | 0 | SFRP1 |

Supplementary Table 2

DE gene lists for each subcluster within epithelial population of tdTomato RET day 7 organoid data

Cluster 1

| p_val | avg_logFC | pct.1 | pct.2 | p_val_adj | cluster | gene |
|---|---|---|---|---|---|---|
| 1.58E-95 | 0.681097 | 1 | 0.992 | 7.14E-91 | 1 | LAPTM4B |
| 4.63E-33 | 0.675161 | 0.78 | 0.578 | 2.09E-28 | 1 | RGS5 |
| 2.03E-24 | 0.642935 | 0.643 | 0.472 | 9.1282707! | 1 | RSPO3 |
| 3.44E-38 | 0.63987 | 0.891 | 0.802 | 1.55E-33 | 1 | SLC35G1 |
| 4.03E-22 | 0.619144 | 0.711 | 0.504 | 1.82E-17 | 1 | SPP1 |
| 7.7968696 | 0.55352 | 0.363 | 0.268 | 3.51E-05 | 1 | NPY |
| 1.97E-31 | 0.509818 | 0.845 | 0.674 | 8.86E-27 | 1 | TCF24 |

Supplementary Table 2

DE gene lists for each subcluster within epithelial population of tdTomato RET day 7 organoid data

Cluster 2

| p_val | avg_logFC | pct.1 | pct.2 | p_val_adj | cluster | gene |
|---|---|---|---|---|---|---|
| 8.54E-86 | 1.489242 | 0.949 | 0.496 | 3.85E-81 | 2 | PNOC |
| 5.61E-62 | 1.275186 | 0.875 | 0.59 | 2.53E-57 | 2 | IGFBP7 |
| 5.5E-76 | 1.05749 | 0.939 | 0.456 | 2.48E-71 | 2 | MAL |
| 5.04E-52 | 0.967865 | 0.861 | 0.524 | 2.27E-47 | 2 | ALDH1A1 |
| 5.87E-41 | 0.873011 | 0.932 | 0.872 | 2.64E-36 | 2 | GNG11 |
| 1.52639043 | 0.821664 | 0.864 | 0.409 | 6.88E-56 | 2 | ARL4C |
| 1.7722371 | 0.788917 | 0.98 | 0.823 | 7.99E-86 | 2 | PAX2 |
| 6.54E-33 | 0.771525 | 0.953 | 0.842 | 2.95E-28 | 2 | HBD |
| 1.4E-31 | 0.717153 | 0.929 | 0.84 | 6.31E-27 | 2 | ID1 |
| 8.82E-45 | 0.708628 | 0.908 | 0.69 | 3.9742265 | 2 | KIAA1551 |
| 2.0295951 | 0.65341 | 0.746 | 0.386 | 9.15E-36 | 2 | GATA3 |
| 4.35E-52 | 0.628256 | 0.902 | 0.702 | 1.96E-47 | 2 | BCAM |
| 6.44E-56 | 0.622244 | 0.895 | 0.581 | 2.9E-51 | 2 | TFAP2A |
| 1.09E-54 | 0.609322 | 0.956 | 0.775 | 4.9008546 | 2 | ANXA2 |
| 4.5E-71 | 0.580003 | 1 | 0.978 | 2.03E-66 | 2 | CD24 |
| 5.85E-42 | 0.535524 | 0.732 | 0.26 | 2.64E-37 | 2 | GRB14 |
| 3.64E-41 | 0.52164 | 0.98 | 0.884 | 1.64E-36 | 2 | SAT1 |
| 1.91E-39 | 0.510688 | 0.902 | 0.654 | 8.62E-35 | 2 | RDH10 |
| 1.16E-49 | 0.509994 | 0.98 | 0.84 | 5.21E-45 | 2 | EPCAM |

Supplementary Table 2
DE gene lists for each subcluster within epithelial population of tdTomato RET day 7 organoid data
Cluster 3

| p_val | avg_logFC | pct.1 | pct.2 | p_val_adj | cluster | gene |
|---|---|---|---|---|---|---|
| 6.54E-65 | 1.208531 | 1 | 0.982 | 2.9481323 | 3 | HIST1H4C |
| 3.2E-124 | 0.960221 | 0.996 | 0.652 | 1.4E-119 | 3 | KIAA0101 |
| 1.1E-115 | 0.959761 | 0.996 | 0.689 | 5E-111 | 3 | TYMS |
| 4.66E-96 | 0.839394 | 0.996 | 0.933 | 2.1E-91 | 3 | DUT |
| 7.8E-117 | 0.654452 | 1 | 0.99 | 3.5E-112 | 3 | TUBA1B |
| 2.1E-136 | 0.642453 | 1 | 0.998 | 9.3E-132 | 3 | H2AFZ |
| 1.7783980 | 0.61189 | 0.974 | 0.701 | 8.02E-66 | 3 | PCNA |
| 7.95E-48 | 0.587849 | 0.985 | 0.786 | 3.58E-43 | 3 | CENPW |
| 1.42E-74 | 0.587069 | 0.936 | 0.414 | 6.4197163 | 3 | CLSPN |
| 2.17E-66 | 0.554149 | 0.951 | 0.516 | 9.78E-62 | 3 | ORC6 |
| 3.45E-72 | 0.539803 | 0.993 | 0.982 | 1.56E-67 | 3 | RANBP1 |
| 2.52E-59 | 0.537153 | 0.959 | 0.56 | 1.14E-54 | 3 | GINS2 |
| 4.33E-32 | 0.533313 | 0.76 | 0.363 | 1.95E-27 | 3 | HIST1H1D |
| 1.25E-56 | 0.522407 | 0.948 | 0.534 | 5.65E-52 | 3 | ZWINT |
| 5.4E-54 | 0.520573 | 0.858 | 0.272 | 2.43E-49 | 3 | RRM2 |
| 2.6E-53 | 0.51573 | 0.985 | 0.853 | 1.17E-48 | 3 | MCM7 |
| 7.85E-59 | 0.505048 | 0.978 | 0.745 | 3.54E-54 | 3 | HELLS |
| 1.51E-66 | 0.503931 | 0.963 | 0.567 | 6.79E-62 | 3 | RRM1 |
| 1.78E-59 | 0.502611 | 0.899 | 0.265 | 8.02E-55 | 3 | MYBL2 |

Supplementary Table 2

DE gene lists for each subcluster within epithelial population of tdTomato RET day 7 organoid data

Cluster 4

| p_val | avg_logFC | pct.1 | pct.2 | p_val_adj | cluster | gene |
|---|---|---|---|---|---|---|
| 3.4E-119 | 1.683244 | 1 | 0.621 | 1.5E-114 | 4 | CENPF |
| 1.87E-82 | 1.559948 | 0.995 | 0.51 | 8.42E-78 | 4 | UBE2C |
| 6.1E-104 | 1.450561 | 1 | 0.434 | 2.7E-99 | 4 | TOP2A |
| 7.94E-96 | 1.412867 | 0.995 | 0.555 | 3.58E-91 | 4 | CCNB1 |
| 2.5E-157 | 1.283284 | 1 | 0.873 | 1.1E-152 | 4 | HMGB2 |
| 1.51E-81 | 1.262962 | 0.99 | 0.823 | 6.81E-77 | 4 | KPNA2 |
| 2E-122 | 1.252133 | 1 | 0.754 | 9E-118 | 4 | PTTG1 |
| 2.05E-94 | 1.234647 | 0.995 | 0.945 | 9.2351834! | 4 | TUBB4B |
| 1.16E-89 | 1.172178 | 0.98 | 0.494 | 5.24E-85 | 4 | CCNB2 |
| 9.15E-75 | 1.167175 | 0.966 | 0.398 | 4.1221882! | 4 | MKI67 |
| 3.19E-97 | 1.149455 | 1 | 0.824 | 1.44E-92 | 4 | UBE2S |
| 2.51E-87 | 1.146548 | 0.985 | 0.384 | 1.13E-82 | 4 | CDC20 |
| 2.23E-77 | 1.126507 | 0.971 | 0.328 | 1.01E-72 | 4 | ASPM |
| 1.1541967: | 1.115175 | 0.985 | 0.387 | 5.2E-86 | 4 | TPX2 |
| 1.18E-63 | 1.06984 | 0.985 | 0.718 | 5.3E-59 | 4 | TUBA1C |
| 1.04E-84 | 1.020507 | 0.99 | 0.316 | 4.6953873! | 4 | DLGAP5 |
| 1.97E-87 | 0.974085 | 1 | 0.515 | 8.9E-83 | 4 | BIRC5 |
| 1.42E-79 | 0.970419 | 0.985 | 0.681 | 6.39E-75 | 4 | ARL6IP1 |
| 3.52E-88 | 0.966725 | 0.995 | 0.413 | 1.59E-83 | 4 | NUSAP1 |
| 3.98E-69 | 0.950476 | 0.956 | 0.328 | 1.79E-64 | 4 | CENPE |
| 1.42E-81 | 0.937746 | 0.98 | 0.754 | 6.4E-77 | 4 | CKS2 |
| 7.32E-91 | 0.877241 | 0.995 | 0.85 | 3.3E-86 | 4 | CKS1B |
| 6.14E-68 | 0.855999 | 0.966 | 0.345 | 2.77E-63 | 4 | CDKN3 |
| 4.7E-126 | 0.786526 | 1 | 0.991 | 2.1E-121 | 4 | TUBA1B |
| 1.54E-69 | 0.774226 | 0.995 | 0.658 | 6.93E-65 | 4 | H2AFX |
| 1.31E-62 | 0.758238 | 0.927 | 0.294 | 5.92E-58 | 4 | SGOL2 |
| 7.12E-64 | 0.748862 | 0.951 | 0.364 | 3.21E-59 | 4 | PRC1 |
| 4.62E-45 | 0.747837 | 0.893 | 0.377 | 2.0834034( | 4 | CDK1 |
| 4.15E-74 | 0.739222 | 0.98 | 0.357 | 1.87E-69 | 4 | AURKB |
| 6.02E-55 | 0.738507 | 0.912 | 0.26 | 2.7133195: | 4 | HMMR |
| 2.31E-63 | 0.736991 | 0.917 | 0.207 | 1.04E-58 | 4 | PLK1 |
| 1.4E-64 | 0.736663 | 0.961 | 0.66 | 6.3200599( | 4 | SMC4 |
| 2.55E-55 | 0.69846 | 0.902 | 0.238 | 1.1507317! | 4 | CDCA3 |
| 4.47E-62 | 0.696894 | 0.946 | 0.439 | 2.01E-57 | 4 | KIF20B |
| 6.75E-58 | 0.680124 | 0.956 | 0.497 | 3.04E-53 | 4 | CKAP2 |
| 2.25E-63 | 0.677581 | 0.941 | 0.256 | 1.02E-58 | 4 | GTSE1 |
| 6.6497256- | 0.670847 | 0.941 | 0.269 | 3E-55 | 4 | FAM64A |
| 3.6154469( | 0.659867 | 0.976 | 0.539 | 1.63E-55 | 4 | MAD2L1 |
| 2.82E-57 | 0.650218 | 0.927 | 0.271 | 1.27E-52 | 4 | CCNA2 |
| 1.26E-55 | 0.64302 | 0.912 | 0.278 | 5.69E-51 | 4 | NDC80 |
| 7.44E-66 | 0.628934 | 0.961 | 0.333 | 3.35E-61 | 4 | TACC3 |
| 2.7E-52 | 0.615757 | 0.951 | 0.578 | 1.22E-47 | 4 | MIS18BP1 |
| 4.93E-54 | 0.607663 | 0.893 | 0.153 | 2.22E-49 | 4 | CENPA |
| 3.78E-47 | 0.606379 | 0.839 | 0.163 | 1.7E-42 | 4 | AURKA |
| 1.08E-54 | 0.58671 | 0.902 | 0.286 | 4.8489841: | 4 | PSRC1 |
| 6.44E-36 | 0.583818 | 0.741 | 0.095 | 2.9E-31 | 4 | PIF1 |

| | | | | | | |
|---|---|---|---|---|---|---|
| 8.93E-52 | 0.582003 | 0.912 | 0.254 | 4.02E-47 | 4 | NUF2 |
| 9.06E-61 | 0.574824 | 1 | 0.983 | 4.08E-56 | 4 | HN1 |
| 3.86E-52 | 0.564041 | 0.888 | 0.246 | 1.74E-47 | 4 | KIF2C |
| 7.77E-55 | 0.563945 | 0.912 | 0.234 | 3.5033221[illegible] | 4 | TROAP |
| 8.88E-45 | 0.562258 | 0.971 | 0.77 | 4.0013459[illegible] | 4 | BUB3 |
| 4.58E-46 | 0.558476 | 0.844 | 0.196 | 2.06E-41 | 4 | CDCA8 |
| 2.74E-42 | 0.552957 | 0.985 | 0.796 | 1.23E-37 | 4 | CENPW |
| 2.21E-66 | 0.549367 | 1 | 0.841 | 9.96E-62 | 4 | DTYMK |
| 1.64E-46 | 0.544394 | 0.844 | 0.213 | 7.37E-42 | 4 | TTK |
| 2.01E-46 | 0.536236 | 0.878 | 0.342 | 9.04E-42 | 4 | KIF11 |
| 5.07E-45 | 0.534895 | 0.883 | 0.272 | 2.2837013[illegible] | 4 | ECT2 |
| 9.56E-71 | 0.52356 | 1 | 0.989 | 4.31E-66 | 4 | NUCKS1 |
| 7.99E-42 | 0.522476 | 0.824 | 0.16 | 3.6E-37 | 4 | KIF20A |
| 1.08E-45 | 0.520185 | 0.893 | 0.293 | 4.87E-41 | 4 | NCAPG |
| 1.45E-45 | 0.5116 | 0.888 | 0.287 | 6.56E-41 | 4 | KIFC1 |
| 2.05E-86 | 0.510813 | 1 | 0.998 | 9.22E-82 | 4 | H2AFZ |
| 1.25E-42 | 0.50035 | 0.805 | 0.146 | 5.63E-38 | 4 | DEPDC1 |
