## Supplementary Table 3 for "*In vitro* plasticity between ureteric epithelial and distal nephron identity and maturity is controlled by extracellular signals"

DE genes for each cluster identified in GATA3 mCherry day 7+14 organoids cultured with RSPO1

Cluster 0

| p_val | avg_logFC | pct.1 | pct.2 | p_val_adj | cluster | gene |
|---|---|---|---|---|---|---|
| 0 | 2.196206 | 0.986 | 0.315 | 0 | 0 | WFDC2 |
| 0 | 1.735963 | 0.97 | 0.101 | 0 | 0 | EPCAM |
| 0 | 1.631594 | 0.93 | 0.149 | 0 | 0 | IGFBP7 |
| 0 | 1.575296 | 0.908 | 0.06 | 0 | 0 | KRT19 |
| 0 | 1.499527 | 0.607 | 0.074 | 0 | 0 | TFPI2 |
| 0 | 1.436456 | 0.559 | 0.134 | 0 | 0 | DKK1 |
| 0 | 1.420281 | 0.884 | 0.088 | 0 | 0 | MAL |
| 0 | 1.412173 | 0.964 | 0.18 | 0 | 0 | EMX2 |
| 0 | 1.296583 | 0.976 | 0.546 | 0 | 0 | SAT1 |
| 0 | 1.280144 | 0.948 | 0.074 | 0 | 0 | PAX2 |
| 0 | 1.277932 | 0.463 | 0.039 | 0 | 0 | PNOC |
| 0 | 1.238794 | 0.939 | 0.479 | 0 | 0 | GNG11 |
| 0 | 1.216464 | 0.736 | 0.049 | 0 | 0 | PCSK1 |
| 0 | 1.216086 | 0.983 | 0.377 | 0 | 0 | SPINT2 |
| 0 | 1.165797 | 0.934 | 0.253 | 0 | 0 | PLEKHA1 |
| 0 | 1.161376 | 0.979 | 0.326 | 0 | 0 | KRT18 |
| 0 | 1.157445 | 0.907 | 0.231 | 0 | 0 | MECOM |
| 0 | 1.118634 | 0.973 | 0.562 | 0 | 0 | CAMK2N1 |
| 0 | 1.102699 | 0.891 | 0.051 | 0 | 0 | CLDN3 |
| 0 | 1.100913 | 0.9 | 0.047 | 0 | 0 | DCDC2 |
| 0 | 1.087214 | 0.965 | 0.6 | 0 | 0 | APOE |
| 0 | 1.059158 | 0.856 | 0.068 | 0 | 0 | LHX1 |
| 0 | 1.050045 | 0.941 | 0.176 | 0 | 0 | DMKN |
| 0 | 1.044972 | 0.656 | 0.033 | 0 | 0 | TMEM52B |
| 0 | 1.043647 | 0.864 | 0.055 | 0 | 0 | CLDN4 |
| 0 | 0.993915 | 0.886 | 0.047 | 0 | 0 | CLDN7 |
| 0 | 0.923117 | 0.725 | 0.187 | 0 | 0 | SCUBE3 |
| 0 | 0.89067 | 0.857 | 0.226 | 0 | 0 | HOXD8 |
| 0 | 0.870419 | 0.954 | 0.585 | 0 | 0 | MPC2 |
| 0 | 0.867747 | 0.921 | 0.414 | 0 | 0 | ATP1B1 |
| 0 | 0.862197 | 0.95 | 0.239 | 0 | 0 | KRT8 |
| 0 | 0.861726 | 0.971 | 0.583 | 0 | 0 | CYB5A |
| 0 | 0.852125 | 0.868 | 0.278 | 0 | 0 | PTBP3 |
| 0 | 0.833084 | 0.523 | 0.033 | 0 | 0 | ALDH1A1 |
| 0 | 0.822745 | 0.872 | 0.154 | 0 | 0 | VAMP8 |
| 0 | 0.818592 | 0.993 | 0.889 | 0 | 0 | ITM2B |
| 0 | 0.814663 | 0.573 | 0.045 | 0 | 0 | GATA3 |
| 0 | 0.807993 | 0.81 | 0.035 | 0 | 0 | RAB25 |
| 0 | 0.774632 | 0.777 | 0.035 | 0 | 0 | CLDN6 |
| 0 | 0.757007 | 0.942 | 0.479 | 0 | 0 | CYBA |
| 0 | 0.756619 | 0.863 | 0.178 | 0 | 0 | BCAM |
| 0 | 0.755541 | 0.988 | 0.709 | 0 | 0 | PCBD1 |
| 5.4E-227 | 0.753437 | 0.32 | 0.037 | 1.8E-222 | 0 | FGF8 |
| 0 | 0.740911 | 0.995 | 0.877 | 0 | 0 | CD24 |
| 1.9E-103 | 0.732733 | 0.193 | 0.011 | 6.5E-99 | 0 | DEFB1 |
| 0 | 0.729304 | 0.871 | 0.188 | 0 | 0 | DSP |

| | | | | | | |
|---|---|---|---|---|---|---|
| 0 | 0.726521 | 0.774 | 0.219 | 0 | 0 | LINC01116 |
| 0 | 0.718529 | 0.845 | 0.337 | 0 | 0 | BST2 |
| 0 | 0.691146 | 0.754 | 0.051 | 0 | 0 | CEBPD |
| 0 | 0.690403 | 0.788 | 0.031 | 0 | 0 | AP1M2 |
| 0 | 0.683141 | 1 | 0.999 | 0 | 0 | GSTP1 |
| 0 | 0.681231 | 0.812 | 0.068 | 0 | 0 | LINC00958 |
| 0 | 0.680338 | 0.825 | 0.059 | 0 | 0 | SPINT1 |
| 0 | 0.674967 | 0.788 | 0.215 | 0 | 0 | FAM181B |
| 0 | 0.663772 | 0.77 | 0.032 | 0 | 0 | HPN |
| 0 | 0.661411 | 0.926 | 0.354 | 0 | 0 | COL18A1 |
| 0 | 0.646359 | 0.487 | 0.064 | 0 | 0 | HTRA1 |
| 0 | 0.640291 | 0.825 | 0.254 | 0 | 0 | BMP7 |
| 0 | 0.628642 | 0.985 | 0.809 | 0 | 0 | S100A13 |
| 0 | 0.622391 | 0.78 | 0.033 | 0 | 0 | LLGL2 |
| 0 | 0.620705 | 0.717 | 0.031 | 0 | 0 | CYS1 |
| 0 | 0.613457 | 0.877 | 0.307 | 0 | 0 | AGPAT2 |
| 0 | 0.602557 | 0.858 | 0.314 | 0 | 0 | EGFL7 |
| 0 | 0.600925 | 0.87 | 0.4 | 0 | 0 | UTRN |
| 0 | 0.599303 | 0.978 | 0.815 | 0 | 0 | PTGR1 |
| 0 | 0.597874 | 0.997 | 0.943 | 0 | 0 | ITM2C |
| 0 | 0.594057 | 0.727 | 0.219 | 0 | 0 | JAG1 |
| 0 | 0.589305 | 0.832 | 0.285 | 0 | 0 | EMID1 |
| 0 | 0.588588 | 0.784 | 0.08 | 0 | 0 | HOOK1 |
| 0 | 0.583691 | 0.835 | 0.247 | 0 | 0 | ANXA11 |
| 0 | 0.578696 | 0.716 | 0.042 | 0 | 0 | ITGA2 |
| 9.9E-181 | 0.568095 | 0.278 | 0.01 | 3.3E-176 | 0 | MMP7 |
| 0 | 0.566399 | 0.809 | 0.197 | 0 | 0 | MFHAS1 |
| 0 | 0.56637 | 0.523 | 0.022 | 0 | 0 | S100A14 |
| 0 | 0.563002 | 0.644 | 0.051 | 0 | 0 | COBLL1 |
| 0 | 0.557329 | 0.811 | 0.207 | 0 | 0 | POU3F3 |
| 0 | 0.55443 | 0.763 | 0.365 | 0 | 0 | ARL4C |
| 0 | 0.550374 | 0.755 | 0.12 | 0 | 0 | LAMA1 |
| 0 | 0.549431 | 0.999 | 0.999 | 0 | 0 | MT-ND4 |
| 0 | 0.54519 | 0.882 | 0.461 | 0 | 0 | ATP1A1 |
| 0 | 0.542663 | 0.699 | 0.034 | 0 | 0 | GPR160 |
| 0 | 0.542138 | 0.984 | 0.759 | 0 | 0 | NR2F6 |
| 0 | 0.540415 | 0.79 | 0.14 | 0 | 0 | GPRC5C |
| 0 | 0.539507 | 0.836 | 0.257 | 0 | 0 | MYO6 |
| 0 | 0.532851 | 0.954 | 0.627 | 0 | 0 | APLP2 |
| 0 | 0.532384 | 0.807 | 0.287 | 0 | 0 | ID4 |
| 0 | 0.529757 | 0.721 | 0.091 | 0 | 0 | EMX2OS |
| 0 | 0.52615 | 0.835 | 0.335 | 0 | 0 | GNAI1 |
| 0 | 0.519154 | 0.698 | 0.043 | 0 | 0 | GRB14 |
| 0 | 0.518166 | 0.799 | 0.3 | 0 | 0 | SMIM14 |
| 0 | 0.517788 | 0.804 | 0.342 | 0 | 0 | NKAIN4 |
| 0 | 0.516712 | 0.702 | 0.132 | 0 | 0 | CD9 |
| 0 | 0.514998 | 0.997 | 0.876 | 0 | 0 | HMGA1 |
| 0 | 0.510982 | 0.699 | 0.061 | 0 | 0 | CRACR2B |
| 0 | 0.502438 | 0.687 | 0.206 | 0 | 0 | S100A4 |
| 0 | 0.501988 | 0.754 | 0.123 | 0 | 0 | S100A16 |

Supplementary Table 3

DE genes for each cluster identified in GATA3 mCherry day 7+14 organoids cultured with RSPO1

Cluster 1

| p_val | avg_logFC | pct.1 | pct.2 | p_val_adj | cluster | gene |
|---|---|---|---|---|---|---|
| 0 | 2.065958 | 0.921 | 0.367 | 0 | 1 | LGALS1 |
| 0 | 2.061804 | 0.774 | 0.374 | 0 | 1 | IGFBP5 |
| 0 | 1.504341 | 0.889 | 0.111 | 0 | 1 | COL3A1 |
| 0 | 1.300191 | 0.851 | 0.114 | 0 | 1 | TWIST1 |
| 0 | 1.240351 | 0.979 | 0.897 | 0 | 1 | VIM |
| 0 | 1.171948 | 0.549 | 0.054 | 0 | 1 | LUM |
| 0 | 1.108007 | 0.988 | 0.71 | 0 | 1 | CALD1 |
| 0 | 1.081474 | 0.731 | 0.162 | 0 | 1 | SCX |
| 0 | 1.060016 | 0.958 | 0.77 | 0 | 1 | TPM1 |
| 0 | 1.058763 | 0.972 | 0.594 | 0 | 1 | COL1A2 |
| 0 | 1.057707 | 0.797 | 0.276 | 0 | 1 | L1TD1 |
| 0 | 1.05367 | 0.944 | 0.837 | 0 | 1 | PEG10 |
| 0 | 1.030388 | 0.57 | 0.056 | 0 | 1 | DCN |
| 0 | 1.008496 | 0.728 | 0.189 | 0 | 1 | COL1A1 |
| 0 | 0.969674 | 0.908 | 0.525 | 0 | 1 | PTN |
| 0 | 0.953301 | 0.82 | 0.109 | 0 | 1 | DNM3OS |
| 0 | 0.952754 | 0.894 | 0.449 | 0 | 1 | COL6A2 |
| 0 | 0.947941 | 0.878 | 0.632 | 0 | 1 | GPC3 |
| 0 | 0.932872 | 0.861 | 0.3 | 0 | 1 | PCOLCE |
| 0 | 0.924003 | 0.951 | 0.788 | 0 | 1 | TCF4 |
| 0 | 0.902503 | 0.936 | 0.708 | 0 | 1 | CRABP2 |
| 0 | 0.884314 | 0.647 | 0.152 | 0 | 1 | SFRP1 |
| 0 | 0.842485 | 0.747 | 0.282 | 0 | 1 | NFIB |
| 0 | 0.83206 | 0.579 | 0.197 | 0 | 1 | TBX3 |
| 0 | 0.81056 | 0.769 | 0.329 | 0 | 1 | RAMP2 |
| 0 | 0.804657 | 0.539 | 0.053 | 0 | 1 | PRRX1 |
| 0 | 0.791243 | 0.918 | 0.426 | 0 | 1 | MEIS2 |
| 0 | 0.783433 | 0.758 | 0.195 | 0 | 1 | IGF2.1 |
| 0 | 0.766095 | 0.679 | 0.106 | 0 | 1 | EDNRA |
| 0 | 0.765454 | 0.631 | 0.053 | 0 | 1 | MEOX2 |
| 0 | 0.73547 | 0.927 | 0.617 | 0 | 1 | CDC42EP5 |
| 0 | 0.71933 | 0.591 | 0.059 | 0 | 1 | HIC1 |
| 1.5100648[illegible] | 0.713267 | 0.325 | 0.043 | 5.1E-196 | 1 | DLK1 |
| 6E-308 | 0.693216 | 0.4 | 0.046 | 2E-303 | 1 | NPR3 |
| 0 | 0.681738 | 0.895 | 0.601 | 0 | 1 | TPM2 |
| 0 | 0.677042 | 0.775 | 0.306 | 0 | 1 | LRRC17 |
| 0 | 0.670796 | 0.631 | 0.121 | 0 | 1 | TBX2 |
| 0 | 0.668727 | 0.844 | 0.395 | 0 | 1 | TIMP1 |
| 0 | 0.663778 | 0.927 | 0.771 | 0 | 1 | SEPT6 |
| 0 | 0.663693 | 0.746 | 0.339 | 0 | 1 | ZFHX4 |
| 0 | 0.662407 | 0.625 | 0.196 | 0 | 1 | PLK2 |
| 0 | 0.6613 | 0.602 | 0.143 | 0 | 1 | NFIA |
| 0 | 0.657525 | 0.571 | 0.066 | 0 | 1 | OLFML3 |
| 0 | 0.654481 | 0.904 | 0.613 | 0 | 1 | VCAN |
| 0 | 0.648979 | 0.691 | 0.227 | 0 | 1 | IL11RA |
| 0 | 0.636065 | 0.707 | 0.326 | 0 | 1 | THY1 |

| | | | | | | |
|---|---|---|---|---|---|---|
| 0 | 0.635193 | 0.576 | 0.114 | 0 | 1 | PDGFRB |
| 0 | 0.633579 | 0.656 | 0.125 | 0 | 1 | PDGFRA |
| 0 | 0.632979 | 0.969 | 0.843 | 0 | 1 | LAPTM4A |
| 0 | 0.628778 | 0.566 | 0.054 | 0 | 1 | GAS2 |
| 1.1E-234 | 0.62461 | 0.431 | 0.119 | 3.5684604[illegible] | 1 | POSTN |
| 0 | 0.616415 | 0.821 | 0.44 | 0 | 1 | GYPC |
| 0 | 0.613782 | 0.642 | 0.228 | 0 | 1 | TSHZ2 |
| 2.4E-183 | 0.613463 | 0.282 | 0.024 | 8.1E-179 | 1 | IGF1 |
| 0 | 0.608576 | 0.623 | 0.117 | 0 | 1 | PCDH18 |
| 0 | 0.605705 | 0.545 | 0.203 | 0 | 1 | ITPR2 |
| 0 | 0.598453 | 0.824 | 0.345 | 0 | 1 | MDFI |
| 0 | 0.590611 | 0.594 | 0.247 | 0 | 1 | SVIL |
| 0 | 0.588439 | 0.762 | 0.36 | 0 | 1 | EMP3 |
| 0 | 0.580402 | 0.863 | 0.717 | 0 | 1 | PBX1 |
| 0 | 0.577787 | 0.997 | 0.982 | 0 | 1 | MAGED2 |
| 0 | 0.564622 | 0.873 | 0.629 | 0 | 1 | RBFOX2 |
| 1.7E-267 | 0.554599 | 0.424 | 0.125 | 5.8E-263 | 1 | DHRS3 |
| 0 | 0.54959 | 0.604 | 0.102 | 0 | 1 | EMILIN1 |
| 0 | 0.544592 | 0.625 | 0.106 | 0 | 1 | INKA1 |
| 0 | 0.54063 | 0.595 | 0.102 | 0 | 1 | DDR2 |
| 0 | 0.535476 | 0.487 | 0.128 | 0 | 1 | PRRX2 |
| 2.1E-267 | 0.526477 | 0.981 | 0.946 | 7.1E-263 | 1 | KCNQ1OT1 |
| 4E-135 | 0.519808 | 0.216 | 0.047 | 1.3547634[illegible] | 1 | CHODL |
| 0 | 0.514051 | 0.687 | 0.292 | 0 | 1 | GAS1 |
| 0 | 0.509891 | 0.86 | 0.662 | 0 | 1 | MFAP2 |
| 5E-233 | 0.504949 | 0.586 | 0.331 | 1.7E-228 | 1 | SNCG |

Supplementary Table 3
DE genes for each cluster identified in GATA3 mCherry day 7+14 organoids cultured with RSPO1
Cluster 2

| p_val | avg_logFC | pct.1 | pct.2 | p_val_adj | cluster | gene |
|---|---|---|---|---|---|---|
| 0 | 1.978176 | 0.91 | 0.623 | 0 | 2 | NEFM |
| 0 | 1.566773 | 0.994 | 0.714 | 0 | 2 | METRN |
| 0 | 1.535344 | 0.934 | 0.209 | 0 | 2 | TTYH1 |
| 0 | 1.46931 | 0.92 | 0.137 | 0 | 2 | SOX2 |
| 0 | 1.262576 | 0.858 | 0.512 | 0 | 2 | NEFL |
| 0 | 1.197114 | 0.882 | 0.41 | 0 | 2 | FGFBP3 |
| 0 | 1.022779 | 0.998 | 0.979 | 0 | 2 | HMGN2 |
| 0 | 0.982507 | 0.849 | 0.277 | 0 | 2 | ZIC2 |
| 0 | 0.980861 | 0.791 | 0.492 | 0 | 2 | AP1S2 |
| 0 | 0.928955 | 0.675 | 0.336 | 0 | 2 | SSFA2 |
| 8.5E-256 | 0.862005 | 0.478 | 0.102 | 2.9E-251 | 2 | MSX1 |
| 0 | 0.831088 | 0.976 | 0.724 | 0 | 2 | CKB |
| 0 | 0.830392 | 0.726 | 0.113 | 0 | 2 | POU3F2 |
| 2.2E-242 | 0.815168 | 0.552 | 0.187 | 7.3E-238 | 2 | AC103702.2 |
| 0 | 0.797844 | 0.979 | 0.661 | 0 | 2 | HOXB9 |
| 0 | 0.763178 | 1 | 0.987 | 0 | 2 | TUBA1A |
| 1.4E-209 | 0.74589 | 0.839 | 0.705 | 4.8E-205 | 2 | HMGB2 |
| 0 | 0.745311 | 1 | 0.996 | 0 | 2 | H2AFZ |
| 0 | 0.738357 | 0.865 | 0.372 | 0 | 2 | CDH2 |
| 0 | 0.729948 | 0.999 | 0.994 | 0 | 2 | TUBA1B |
| 7.8E-199 | 0.720912 | 0.36 | 0.029 | 2.6E-194 | 2 | S100B |
| 0 | 0.720256 | 0.777 | 0.349 | 0 | 2 | SFRP2 |
| 3.9E-295 | 0.71045 | 0.608 | 0.216 | 1.2984279! | 2 | NTRK2 |
| 0 | 0.705036 | 0.751 | 0.205 | 0 | 2 | HOXC9 |
| 0 | 0.699674 | 0.754 | 0.152 | 0 | 2 | FEZ1 |
| 1.9E-135 | 0.694694 | 0.688 | 0.516 | 6.4E-131 | 2 | CENPF |
| 0 | 0.69434 | 0.757 | 0.331 | 0 | 2 | AC004540.2 |
| 0 | 0.6899 | 0.668 | 0.236 | 0 | 2 | DUSP4 |
| 0 | 0.686987 | 0.985 | 0.839 | 0 | 2 | COTL1 |
| 0 | 0.686278 | 0.641 | 0.089 | 0 | 2 | GRID2 |
| 0 | 0.67085 | 0.533 | 0.061 | 0 | 2 | HOXC10 |
| 1.4E-235 | 0.665508 | 0.893 | 0.816 | 4.8E-231 | 2 | CCND2 |
| 0 | 0.656148 | 0.974 | 0.906 | 0 | 2 | LAPTM4B |
| 0 | 0.651164 | 0.742 | 0.357 | 0 | 2 | TUBB2A |
| 0 | 0.646741 | 0.948 | 0.717 | 0 | 2 | CENPV |
| 1.2E-243 | 0.64537 | 0.876 | 0.711 | 3.9E-239 | 2 | IGDCC3 |
| 0 | 0.644482 | 0.576 | 0.053 | 0 | 2 | GDF10 |
| 0 | 0.636737 | 0.656 | 0.191 | 0 | 2 | ZEB2 |
| 7.9E-282 | 0.614641 | 0.865 | 0.624 | 2.7E-277 | 2 | NR2F1 |
| 1.6E-154 | 0.61133 | 0.87 | 0.781 | 5.4442348! | 2 | UBE2S |
| 5.2260635! | 0.606936 | 0.556 | 0.401 | 1.75E-75 | 2 | UBE2C |
| 5.8E-158 | 0.599156 | 0.327 | 0.046 | 1.9E-153 | 2 | SLC18A3 |
| 2.5704540! | 0.596651 | 0.793 | 0.583 | 8E-306 | 2 | WLS |
| 0 | 0.589905 | 0.777 | 0.31 | 0 | 2 | BOC |
| 9.9E-172 | 0.587037 | 0.972 | 0.938 | 3.3E-167 | 2 | ID3 |
| 0 | 0.585801 | 0.531 | 0.046 | 0 | 2 | PTPRZ1 |

| | | | | | | |
|---|---|---|---|---|---|---|
| 3.1E-141 | 0.584409 | 0.346 | 0.079 | 1E-136 | 2 | FABP7 |
| 3.3E-166 | 0.576405 | 0.62 | 0.389 | 1.1E-161 | 2 | BIRC5 |
| 8.9145249: | 0.574461 | 0.915 | 0.898 | 2.99E-95 | 2 | TUBB4B |
| 2.32E-82 | 0.574394 | 0.188 | 0.041 | 7.8E-78 | 2 | HES5 |
| 3.5E-253 | 0.571708 | 0.451 | 0.047 | 1.2E-248 | 2 | AL359091.1 |
| 1.2225019! | 0.571088 | 0.563 | 0.396 | 4.1E-96 | 2 | TOP2A |
| 2.57E-72 | 0.568912 | 0.492 | 0.413 | 8.62E-68 | 2 | CCNB1 |
| 0 | 0.566578 | 0.57 | 0.093 | 0 | 2 | LINC00461 |
| 1.7E-271 | 0.56374 | 0.661 | 0.353 | 5.9E-267 | 2 | NLRP1 |
| 0 | 0.5619 | 1 | 0.971 | 0 | 2 | MARCKS |
| 6.2E-84 | 0.560056 | 0.728 | 0.671 | 2.08E-79 | 2 | PTTG1 |
| 0 | 0.559035 | 0.996 | 0.794 | 0 | 2 | RBP1 |
| 0 | 0.55721 | 0.529 | 0.045 | 0 | 2 | RFX4 |
| 4.9E-184 | 0.553299 | 0.7 | 0.53 | 1.6E-179 | 2 | H2AFX |
| 4.1E-227 | 0.549909 | 0.414 | 0.063 | 1.4E-222 | 2 | C1QTNF3 |
| 6.4249007! | 0.548221 | 0.841 | 0.622 | 2.2E-235 | 2 | TYMS |
| 1.1E-141 | 0.547253 | 0.569 | 0.381 | 3.5E-137 | 2 | NUSAP1 |
| 0 | 0.543156 | 0.913 | 0.771 | 0 | 2 | CSRP2 |
| 9E-304 | 0.537704 | 0.795 | 0.515 | 3E-299 | 2 | COL2A1 |
| 5.8E-126 | 0.535457 | 0.564 | 0.36 | 1.9E-121 | 2 | CCNB2 |
| 0 | 0.535012 | 0.937 | 0.754 | 0 | 2 | TMSB15A |
| 0 | 0.532726 | 0.513 | 0.083 | 5E-305 | 2 | COL14A1 |
| 5.08E-86 | 0.532231 | 0.782 | 0.735 | 1.7E-81 | 2 | KPNA2 |
| 7.66E-21 | 0.532185 | 0.967 | 0.984 | 2.57E-16 | 2 | HIST1H4C |
| 5.16E-22 | 0.531617 | 0.421 | 0.406 | 1.73E-17 | 2 | TGFBI |
| 4.6E-212 | 0.530856 | 0.727 | 0.516 | 1.6E-207 | 2 | CENPW |
| 8.5E-229 | 0.525211 | 0.488 | 0.141 | 2.8E-224 | 2 | SLC1A3 |
| 0 | 0.523357 | 1 | 0.993 | 0 | 2 | TUBB2B |
| 2.35E-88 | 0.52186 | 0.468 | 0.324 | 7.87E-84 | 2 | CDC20 |
| 2.4E-248 | 0.505474 | 0.977 | 0.959 | 7.9E-244 | 2 | FDPS |
| 2.4E-208 | 0.504255 | 0.522 | 0.189 | 8.1E-204 | 2 | APCDD1 |
| 1.8E-261 | 0.500194 | 0.467 | 0.084 | 6E-257 | 2 | EDNRB |

Supplementary Table 3
DE genes for each cluster identified in GATA3 mCherry day 7+14 organoids cultured with RSPO1
Cluster 3

| p_val | avg_logFC | pct.1 | pct.2 | p_val_adj | cluster | gene |
|---|---|---|---|---|---|---|
| 4.3457531 | 2.40217 | 0.969 | 0.598 | 1.5E-115 | 3 | CRABP1 |
| 4.6E-101 | 2.397334 | 0.96 | 0.662 | 1.55E-96 | 3 | HES6 |
| 6.32E-45 | 2.189548 | 0.554 | 0.075 | 2.1207874 | 3 | STMN2 |
| 3.5E-117 | 2.021562 | 0.918 | 0.102 | 1.2E-112 | 3 | TAGLN3 |
| 3.8E-108 | 1.85083 | 0.977 | 0.653 | 1.3E-103 | 3 | MAP1B |
| 6.98E-55 | 1.67712 | 0.706 | 0.121 | 2.3422065 | 3 | GADD45G |
| 3.77E-72 | 1.574138 | 0.808 | 0.278 | 1.27E-67 | 3 | DLL3 |
| 1.6227340 | 1.449118 | 0.78 | 0.028 | 5.44E-76 | 3 | INSM1 |
| 1.2E-125 | 1.412175 | 0.992 | 0.781 | 4.1E-121 | 3 | CKB |
| 3.9E-143 | 1.382444 | 1 | 0.99 | 1.3E-138 | 3 | TUBA1A |
| 4.6E-104 | 1.355446 | 0.879 | 0.075 | 1.5E-99 | 3 | ELAVL2 |
| 3.88E-57 | 1.315933 | 0.667 | 0.1 | 1.3E-52 | 3 | SSTR2 |
| 2.95E-91 | 1.308987 | 0.958 | 0.641 | 9.88E-87 | 3 | MLLT11 |
| 2.81E-28 | 1.270372 | 0.89 | 0.932 | 9.41E-24 | 3 | CCND1 |
| 4.0574758 | 1.251292 | 0.887 | 0.227 | 1.36E-85 | 3 | MAP2 |
| 3.33E-75 | 1.23725 | 0.763 | 0.015 | 1.1164413 | 3 | NHLH1 |
| 1.5E-116 | 1.218593 | 0.994 | 0.777 | 5E-112 | 3 | SOX11 |
| 4.44E-85 | 1.206473 | 0.782 | 0.037 | 1.4905905 | 3 | ELAVL3 |
| 2.16E-48 | 1.202434 | 0.605 | 0.089 | 7.25E-44 | 3 | RGS16 |
| 1.91E-77 | 1.166415 | 0.881 | 0.333 | 6.42E-73 | 3 | CRMP1 |
| 1.26E-91 | 1.153364 | 0.814 | 0.048 | 4.24E-87 | 3 | IGFBPL1 |
| 6.8E-154 | 1.129163 | 1 | 0.841 | 2.3E-149 | 3 | RBP1 |
| 4.44E-54 | 1.10479 | 0.924 | 0.819 | 1.49E-49 | 3 | CDKN1C |
| 1.07E-61 | 1.056337 | 0.766 | 0.24 | 3.59E-57 | 3 | PHLDA1 |
| 8.36E-35 | 1.019653 | 0.477 | 0.059 | 2.8032051 | 3 | PPP1R17 |
| 3.28E-79 | 1.019484 | 0.751 | 0.006 | 1.1E-74 | 3 | NEUROD4 |
| 5.52E-34 | 1.00259 | 0.446 | 0.068 | 1.85E-29 | 3 | TMEM176A |
| 7.8E-119 | 1.002266 | 1 | 0.982 | 2.6E-114 | 3 | BASP1 |
| 3.2E-106 | 0.991459 | 1 | 0.995 | 1.1E-101 | 3 | TUBB2B |
| 1.2E-128 | 0.98579 | 0.997 | 0.734 | 4.2E-124 | 3 | HOXB9 |
| 3.85E-62 | 0.970309 | 0.661 | 0.028 | 1.29E-57 | 3 | ELAVL4 |
| 5.33E-73 | 0.965388 | 0.743 | 0.038 | 1.79E-68 | 3 | MAP6 |
| 7.22E-37 | 0.962475 | 0.449 | 0.009 | 2.42E-32 | 3 | VXN |
| 1.21E-43 | 0.955404 | 0.743 | 0.44 | 4.07E-39 | 3 | GADD45A |
| 7.17E-59 | 0.937582 | 0.624 | 0.019 | 2.4E-54 | 3 | SCG3 |
| 8.48E-55 | 0.935694 | 0.621 | 0.011 | 2.8449693 | 3 | NEUROG2 |
| 3.28E-27 | 0.919081 | 0.842 | 0.691 | 1.1E-22 | 3 | NEFM |
| 6.6E-79 | 0.917759 | 0.873 | 0.425 | 2.21E-74 | 3 | PCBP4 |
| 9.5E-54 | 0.910584 | 0.621 | 0.072 | 3.19E-49 | 3 | DCX |
| 3.93E-33 | 0.904548 | 0.432 | 0.006 | 1.32E-28 | 3 | NEUROD1 |
| 7.32E-72 | 0.903722 | 0.845 | 0.352 | 2.45E-67 | 3 | DST |
| 2.33E-64 | 0.896285 | 0.763 | 0.179 | 7.8231021 | 3 | RASL11B |
| 2.42E-96 | 0.883401 | 0.963 | 0.597 | 8.13E-92 | 3 | BEX1 |
| 1.64E-39 | 0.881687 | 0.621 | 0.264 | 5.49E-35 | 3 | KIF5C |
| 6.34E-73 | 0.876887 | 0.777 | 0.097 | 2.12E-68 | 3 | DCC |
| 1.11E-75 | 0.86971 | 1 | 0.999 | 3.72E-71 | 3 | TMSB4X |

| | | | | | | |
|---|---|---|---|---|---|---|
| 4.13E-76 | 0.866193 | 0.997 | 0.983 | 1.39E-71 | 3 | SOX4 |
| 1.84E-72 | 0.853678 | 0.904 | 0.631 | 6.17E-68 | 3 | CHD7 |
| 2.24E-29 | 0.851652 | 0.41 | 0.065 | 7.5E-25 | 3 | TMEM176B |
| 1.92E-43 | 0.842658 | 0.669 | 0.269 | 6.45E-39 | 3 | THSD7A |
| 2.03E-35 | 0.836805 | 0.466 | 0.058 | 6.8E-31 | 3 | POU2F2 |
| 1.57E-27 | 0.814607 | 0.347 | 0.015 | 5.25E-23 | 3 | CNTN2 |
| 3.2E-129 | 0.809689 | 1 | 0.999 | 1.1E-124 | 3 | STMN1 |
| 2.33E-59 | 0.80339 | 0.847 | 0.512 | 7.83E-55 | 3 | CPE |
| 6.33E-52 | 0.789769 | 0.712 | 0.265 | 2.12E-47 | 3 | NCAM1 |
| 1.07E-42 | 0.77529 | 0.986 | 0.951 | 3.58E-38 | 3 | CALM1 |
| 2.55E-32 | 0.767251 | 0.418 | 0.006 | 8.55E-28 | 3 | NEUROG1 |
| 7.91E-58 | 0.75796 | 0.734 | 0.268 | 2.65E-53 | 3 | TOX3 |
| 2.31E-42 | 0.755309 | 0.579 | 0.138 | 7.76E-38 | 3 | DRAXIN |
| 3.45E-28 | 0.754867 | 0.918 | 0.8 | 1.16E-23 | 3 | UBE2S |
| 1.3E-23 | 0.739236 | 0.316 | 0.024 | 4.35E-19 | 3 | KLHL35 |
| 2.58E-43 | 0.72985 | 0.559 | 0.105 | 8.66E-39 | 3 | MIAT |
| 6.05E-19 | 0.727581 | 0.249 | 0.01 | 2.03E-14 | 3 | STMN4 |
| 1.96E-19 | 0.727449 | 0.347 | 0.063 | 6.57E-15 | 3 | LAMP5 |
| 1.14E-65 | 0.726485 | 0.994 | 0.964 | 3.81E-61 | 3 | RTN4 |
| 1.64E-34 | 0.726166 | 0.432 | 0.007 | 5.5158467 | 3 | RASD1 |
| 1.67E-41 | 0.72571 | 0.514 | 0.038 | 5.6E-37 | 3 | CHRNA3 |
| 8.25E-74 | 0.724065 | 0.989 | 0.873 | 2.77E-69 | 3 | COTL1 |
| 2.98E-18 | 0.723809 | 0.362 | 0.118 | 1E-13 | 3 | GAP43 |
| 1.9E-125 | 0.708526 | 1 | 0.977 | 6.2E-121 | 3 | MARCKS |
| 8.3223268 | 0.705313 | 0.565 | 0.14 | 2.79E-35 | 3 | PDPN |
| 2.6E-35 | 0.703315 | 0.74 | 0.426 | 8.71E-31 | 3 | RND3 |
| 3.29E-98 | 0.697575 | 1 | 0.966 | 1.1E-93 | 3 | JPT1 |
| 3.9728059 | 0.680753 | 0.692 | 0.228 | 1.33E-45 | 3 | TSPAN18 |
| 3.34E-36 | 0.679641 | 0.706 | 0.325 | 1.12E-31 | 3 | SOX2 |
| 2.89E-59 | 0.677603 | 0.805 | 0.431 | 9.69E-55 | 3 | GPC2 |
| 8.58E-25 | 0.669742 | 0.667 | 0.448 | 2.8768163 | 3 | TUBB2A |
| 1.86E-37 | 0.6609 | 0.571 | 0.17 | 6.23E-33 | 3 | PPP1R14A |
| 4.3042768 | 0.652404 | 0.616 | 0.192 | 1.44E-35 | 3 | BTBD17 |
| 4.3E-26 | 0.65157 | 0.593 | 0.27 | 1.44E-21 | 3 | AC103702.2 |
| 2.53E-29 | 0.644155 | 0.415 | 0.066 | 8.47E-25 | 3 | MYBL1 |
| 1.41E-37 | 0.64379 | 0.624 | 0.271 | 4.74E-33 | 3 | TFAP2A |
| 2.2E-37 | 0.642469 | 0.475 | 0.044 | 7.39E-33 | 3 | RASGEF1B |
| 1.13E-49 | 0.640958 | 0.636 | 0.096 | 3.79E-45 | 3 | BAALC |
| 2.73E-57 | 0.636938 | 0.715 | 0.224 | 9.16E-53 | 3 | RNF165 |
| 1.33E-22 | 0.63254 | 0.458 | 0.184 | 4.47E-18 | 3 | MFAP4 |
| 1.2E-42 | 0.630585 | 0.715 | 0.395 | 4.02E-38 | 3 | SRGAP3 |
| 1.25E-32 | 0.629216 | 0.675 | 0.42 | 4.18E-28 | 3 | ENC1 |
| 4.95E-35 | 0.628486 | 0.562 | 0.205 | 1.6600151 | 3 | RGMB |
| 8.68E-19 | 0.625861 | 0.367 | 0.148 | 2.91E-14 | 3 | NSG1 |
| 3.82E-36 | 0.622746 | 0.799 | 0.604 | 1.28E-31 | 3 | EPB41 |
| 6.46E-44 | 0.614 | 0.734 | 0.254 | 2.17E-39 | 3 | POU3F2 |
| 8.88E-32 | 0.612916 | 0.412 | 0.006 | 2.98E-27 | 3 | TRDC |
| 4.89E-22 | 0.612535 | 0.469 | 0.21 | 1.64E-17 | 3 | DOK5 |
| 5.88E-43 | 0.608915 | 0.859 | 0.564 | 1.97E-38 | 3 | HOXA9 |
| 2.31E-21 | 0.60566 | 0.624 | 0.404 | 7.73E-17 | 3 | PCSK1N |

| | | | | | | |
|---|---|---|---|---|---|---|
| 4.22E-18 | 0.60433 | 0.288 | 0.061 | 1.42E-13 | 3 | RTN1 |
| 5.67E-39 | 0.603598 | 0.955 | 0.904 | 1.9E-34 | 3 | SH3BGRL3 |
| 4.05E-51 | 0.601045 | 0.709 | 0.263 | 1.36E-46 | 3 | SHF |
| 6.74E-25 | 0.599786 | 0.548 | 0.247 | 2.2602839( | 3 | SYT1 |
| 4.8E-51 | 0.596432 | 0.819 | 0.499 | 1.61E-46 | 3 | AUTS2 |
| 6.28E-44 | 0.595675 | 0.825 | 0.465 | 2.11E-39 | 3 | HOXA10 |
| 4.5350836: | 0.593635 | 0.853 | 0.511 | 1.52E-55 | 3 | FYN |
| 1.77E-52 | 0.590566 | 0.76 | 0.328 | 5.93E-48 | 3 | KCNQ2 |
| 1.95E-34 | 0.586864 | 0.531 | 0.171 | 6.5557549( | 3 | DPYSL3 |
| 4.7E-43 | 0.58654 | 0.992 | 0.987 | 1.58E-38 | 3 | CALM2 |
| 1.36E-43 | 0.585846 | 0.514 | 0.025 | 4.57E-39 | 3 | KLHDC8A |
| 1.3E-14 | 0.575582 | 0.251 | 0.024 | 4.3731713! | 3 | ACTC1 |
| 1.68E-44 | 0.575424 | 0.867 | 0.586 | 5.6184352! | 3 | HOXB8 |
| 2.12E-46 | 0.575242 | 0.703 | 0.264 | 7.11E-42 | 3 | ZEB1 |
| 1.35E-36 | 0.573892 | 0.655 | 0.305 | 4.52E-32 | 3 | APLP1 |
| 2.98E-25 | 0.569639 | 0.444 | 0.135 | 1E-20 | 3 | CACNA1A |
| 4.82E-34 | 0.565713 | 0.455 | 0.059 | 1.62E-29 | 3 | ONECUT1 |
| 8.18E-29 | 0.564266 | 0.766 | 0.601 | 2.74E-24 | 3 | DAAM1 |
| 2.4296257( | 0.564021 | 0.384 | 0.008 | 8.15E-26 | 3 | NHLH2 |
| 4.36E-38 | 0.561357 | 0.873 | 0.757 | 1.46E-33 | 3 | TERF2IP |
| 3.97E-24 | 0.558939 | 0.328 | 0.012 | 1.33E-19 | 3 | INA |
| 3.13E-47 | 0.554105 | 0.771 | 0.443 | 1.05E-42 | 3 | GDI1 |
| 2.06E-18 | 0.552398 | 0.294 | 0.091 | 6.9E-14 | 3 | GNG3 |
| 4.17E-26 | 0.55045 | 0.342 | 0.003 | 1.4E-21 | 3 | TLX3 |
| 4.64E-28 | 0.550248 | 0.359 | 0.007 | 1.56E-23 | 3 | POU4F1 |
| 4.13E-06 | 0.550114 | 0.475 | 0.44 | 0.138647 | 3 | UBE2C |
| 3.02E-24 | 0.546568 | 0.415 | 0.115 | 1.01E-19 | 3 | PAX3 |
| 1.7E-27 | 0.545963 | 0.59 | 0.347 | 5.71E-23 | 3 | NOVA1 |
| 1.75E-51 | 0.543154 | 0.918 | 0.762 | 5.87E-47 | 3 | MAP4K4 |
| 8.86E-27 | 0.541468 | 0.514 | 0.246 | 2.97E-22 | 3 | CDKN2D |
| 1.59E-32 | 0.540967 | 0.415 | 0.017 | 5.33E-28 | 3 | NPTX2 |
| 1.01E-39 | 0.540931 | 0.455 | 0.003 | 3.37E-35 | 3 | LINC00599 |
| 9.11E-33 | 0.538974 | 0.415 | 0.004 | 3.06E-28 | 3 | ST18 |
| 1.23E-34 | 0.533188 | 0.531 | 0.157 | 4.1302146 | 3 | ANK2 |
| 1.47E-32 | 0.530603 | 0.788 | 0.603 | 4.93E-28 | 3 | HOXB2 |
| 3.5E-37 | 0.530193 | 0.542 | 0.137 | 1.18E-32 | 3 | EBF1 |
| 4.81E-47 | 0.529209 | 0.681 | 0.228 | 1.61E-42 | 3 | DPYSL4 |
| 6.42E-23 | 0.527344 | 0.333 | 0.032 | 2.15E-18 | 3 | ONECUT2 |
| 4.53E-19 | 0.523148 | 0.285 | 0.04 | 1.52E-14 | 3 | RGS4 |
| 7.49E-47 | 0.522415 | 0.763 | 0.435 | 2.51E-42 | 3 | RCOR2 |
| 1.01E-42 | 0.520931 | 0.867 | 0.639 | 3.4E-38 | 3 | PLPPR3 |
| 1.22E-42 | 0.519005 | 0.797 | 0.428 | 4.08E-38 | 3 | AC004540.2 |
| 6.06E-45 | 0.514306 | 0.927 | 0.827 | 2.0326204! | 3 | DYNC1H1 |
| 2.93E-34 | 0.509607 | 0.661 | 0.376 | 9.8103852: | 3 | RND2 |
| 1.05E-45 | 0.504601 | 0.554 | 0.094 | 3.53E-41 | 3 | PHF21B |
| 5.03E-36 | 0.50457 | 0.582 | 0.171 | 1.69E-31 | 3 | NGFR |
| 1.14E-14 | 0.504453 | 0.251 | 0.05 | 3.8137181( | 3 | POU3F1 |
| 2.94E-35 | 0.502525 | 0.596 | 0.257 | 9.85E-31 | 3 | ROBO2 |
